# Mutations and structural variants arising during double-strand break repair

**DOI:** 10.1101/2025.02.28.640809

**Authors:** Simona Dalin, Sophie Webster, Neal Sugawara, Qiuqin Wu, Shu Zhang, Carmen Macias, Elena Sapède, Tracy Cui, Victoria Liang, Laura Tran, Rameen Beroukhim, James E. Haber

## Abstract

Double-strand break (DSB) repair is highly mutagenic compared to normal replication. In budding yeast, repair of an HO endonuclease-induced DSB at *MATα* can be repaired by using a transcriptionally silent *HMR::Kl-URA3* donor. During repair, -1 deletions in homonucleotide runs are strongly favored over +1 insertions, whereas during replication, spontaneous +1 and -1 events are equal. Microhomology-bounded, repair-associated intragenic deletions (IDs) are recovered 12 times more frequently than tandem duplications (TDs). IDs have a mean length of 56 bp, while TDs average 22 bp. These data suggest a picture of the structure of the repair replication fork: IDs and TDs occur within the open structure of a migrating D-loop, where the 3’ end of a partly copied new DNA strand can dissociate and anneal with a single-stranded region of microhomology that lies either ∼80 bp ahead or ∼40 bp behind the 3’ end. Another major class of repair-associated mutations (∼10%) are interchromosomal template switches (ICTS), even though the *K. lactis URA3* sequence in *HMR* is only 72% identical (homeologous) with *S. cerevisiae ura3-52*. ICTS events begin and end at regions of short (∼7 bp) microhomology; however, ICTS events are constrained to the middle of the copied sequence. Whereas microhomology usage in intragenic deletions is not influenced by adjacent homeology, we show that extensive pairing of adjacent homeology plays a critical role in ICTS. Thus, although by convention, structural variants are characterized by the precise base pairs at their junction, microhomology-mediated template switching actually requires alignment of extensive adjacent homeology.

**Significance statement:** DNA synthesis during repair of a double-strand chromosome break by homologous recombination exhibits a high rate of mutation compared to normal replication. Using a genetic system in budding yeast, we isolated thousands of mutations occurring during repair. We conclude that the repair replication fork appears to have the two DNA strands open ∼80 bp ahead of the DNA polymerase, but the strands re-anneal rapidly behind the polymerase. Additionally, we analyzed interchromosomal template switching, in which the partially copied DNA strand dissociates and pairs with a new template at a short stretch of perfectly matching bases (microhomology), and resumes copying. We show that these apparent microhomology-mediated template switching events in fact require the pairing of ∼200 bp of imperfectly matching bases (homeology).

## Introduction

Double-strand breaks (DSBs) pose an existential threat to cells, which have evolved a variety of repair mechanisms to restore genome integrity. In mammalian cells, where spontaneous DSBs occur multiple times per DNA replication cycle, genes that encode the homologous recombination (HR) repair machinery are essential. In budding yeast, with its much smaller genome, such breaks occur only about one every 10 cell cycles, but when they arise, their repair also requires the homologous recombination proteins such as Rad51 or Rad52 (1). The most accurate of the HR mechanisms is gene conversion, in which both ends of a DSB engage with a homologous template – a sister chromatid, a homolog, or an ectopic sequence – to repair the break with a short patch of new DNA synthesis (synthesis dependent strand annealing, SDSA). A related mechanism is break-induced replication (BIR), in which only one end of the DSB successfully engages the homologous template sequences.

DNA synthesis during DSB repair is strikingly different from normal DNA replication. In SDSA, the predominant mechanism employed in mitotic cells (**Figure 1a**), the two strands copied into the repaired locus are synthesized sequentially and not simultaneously, as they would during normal replication. Moreover, the newly copied DNA is not semi-conservatively replicated; instead, all the newly synthesized DNA is found at the repair site, leaving the donor template unaltered (2,3). Additionally, both strands appear to be synthesized by DNA polymerase δ, which normally is confined to copying short Okazaki-length segments on the lagging strand.

**Figure 1:**
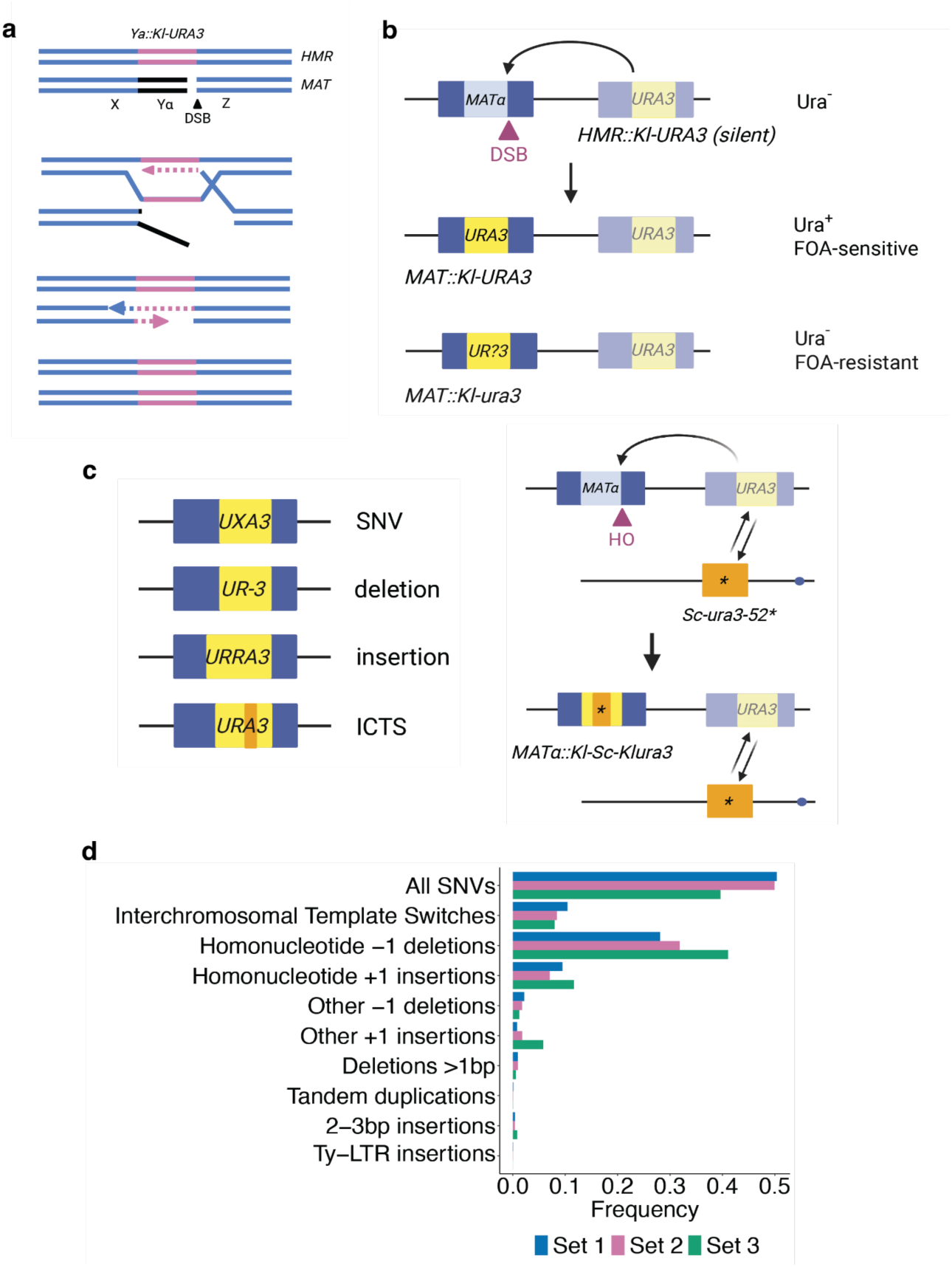
The spectrum of mutational events arising during repair replication. **a**, Schematic of synthesis dependent strand annealing (SDSA) used to repair DSBs induced at the *MAT* locus. After strand invasion of *MAT-Z* into the homologous *HMR-Z* region, new DNA synthesis extends the end of the invading strand to copy Y**a**::*Kl-URA3* (pink). Dissociation of the newly-copied strand (dotted line) allows second-end capture and copying of the second strand to complete the replacement of *MATα* by *MAT*::*Kl-URA3* (pink). **b**, Repair of HO-induced DSB at the *MAT* locus repaired by copying from *HMR::Kl-URA3* results predominantly in Ura^+^ colonies, but some FOA-resistant Ura^-^ cells are recovered. **c**, Types of mutational events observed, including intrachromosomal deletions/tandem duplications and interchromosomal template switches (ICTS). **d**, Normalized frequency of all mutational events observed during repair replication across three independent replicates.

In our previous work, we studied differences between normal and repair replication by modifying the yeast mating-type switching system. In this system, an inducible HO endonuclease creates a DSB at the *MAT* locus that is repaired via SDSA from unexpressed homologous regions, *HML* and *HMR*, located near the two ends of the same chromosome (4). By replacing the **a**1 coding sequences within *HMR***a**, with a *K. lactis URA3* gene (*Kl-URA3*), we could select for highly efficient switching events in which the *Kl-URA3* sequences are copied into the *MAT* locus, replacing Yα and expressing the *Kl-URA3* gene (**Figure 1b**) (5). After DSB induction and the 5’ to 3’ resection of the DSB ends, the right end of the DSB at *MAT* (*MAT-Z1*) strand-invades the perfectly matched 230 bp of homologous sequences at *HMR*. Strand invasion is followed by the initiation of new DNA synthesis, with polymerase δ extending the invading strand by copying *Kl-URA3* sequences. In normal switching events, the newly copied DNA extends into the left homology region, *HML-X* and the displaced, newly copied strand pairs with the resected second end of the DSB (called second-end capture), allowing replacement of the original Yα sequences by *Kl-URA3*. However, if copying of *HMR::Kl-URA3* is interrupted, the partly copied sequence can apparently dissociate from its template and undergo a number of different mutagenic fates, in which the end of the dissociated strand can pair via short microhomologies to produce a variety of mutations that are recovered at *MAT* (5). Virtually all of these Ura^-^ mutants have alterations within the *Kl-URA3* sequences that were copied into the *MAT* locus, while the unexpressed donor sequence remains unaltered. That these mutations arose during switching can be demonstrated by un-silencing *HMR::Kl-URA3* in such mutants, by addition of the Sir2 inhibitor, nicotinamide, whereby Ura^-^ cells become Ura^+^, as the *HMR::Kl-URA3* donor sequence is unaltered.

Compared to mutations arising spontaneously during normal DNA replication of the same sequences, the mutation rate accompanying DSB repair is 1000-fold higher and the mutants display a remarkably different spectrum. Among spontaneous mutations the great majority of events are base pair substitutions, whereas repair-replication events often appear to involve some sort of slippage or dissociation of the partly synthesized new strand. These include -1 frameshifts in homonucleotide runs, intragenic deletions, quasipalindrome mutations, and interchromosomal template switches (ICTSs) that we have documented previously (5–7). Most of these alterations involve the apparent use of microhomologies at the rearrangement junctions.

The most complex events – interchromosomal template switching (ICTS) – require two template “jumps” into and from a 72% identical *S. cerevisiae URA3* sequence (*Sc-ura3-52*) on a different chromosome. After a few hundred bases of *Sc-ura3* are copied, there must be a second template switch back to the original *Kl-URA3* template, where copying proceeds until the shared homology between *HMR-X* and *MAT-X* is reached, so that these sequences can be captured at *MAT* and a second strand synthesis can complete the process. Despite their divergence at the DNA level, most jumps between *Kl-URA3* and *Sc-ura3-52* are in frame, at small microhomologies, so that most events produce a *Kl-Sc-Kl* chimera that is fully functional. Initially, we only recovered Ura^-^ ICTS mutants that misaligned the *Kl* and *Sc* sequences at one particular 5-bp out-of-frame microhomology, resulting in frameshifts (5). In subsequent work, we enriched for ICTS events by demanding correction of a 32-bp deletion in the *Kl-ura3* sequence via copying the deleted bases from *Sc-ura3-52* (5,6). The rate of ICTS is strongly limited by the degree of divergence. When the *Sc-ura3-52* sequences were replaced by *K. lactis* sequences (i.e. the jumps were between identical sequences, correcting the 32-bp deletion), the rate of ICTS increased more than 500-fold, to a level of > 1 in 10^3^ repair events (6).

Our prior work was based on the individual sequencing of independent events, so we were only able to examine the sequence of approximately 50-100 mutational events per condition. Here we report PacBio sequencing and computational analysis of a large population of repair-associated mutations. We ensured recovery of most ICTS events by introducing a -1 frameshift into *Sc-ura3-52* so that most of these template switches would be Ura^-^. From these larger samples we could identify many types of mutations including ICTS events as well as intragenic deletions and insertions, also bounded by microhomologies. We find that DNA synthesis during DNA repair is distinctly different from normal replication. For example, that there are many more -1 deletions than +1 insertions in homonucleotide runs arising during repair suggests either differences in DNA polymerase δ activity or else reflects differences in the mismatch repair system compared to normal replication. Similarly, there are many more, and larger, intragenic deletions, bounded by microhomology, than tandem duplications. This spectrum of mutations reflects the instability of the repair replication fork and its architecture. Specifically, we deduce that the D-loop at the donor containing the newly replicating strand is more open ahead of the DNA polymerase while it is rapidly closed behind the polymerase, as the donor strands reanneal.

Intragenic deletions are not apparently influenced by the degree of homology in adjacent sequences. In contrast, we find that ICTS events are strongly dependent on the alignment of ∼200 bp of homeologous sequence before the polymerase can successfully initiate copying of the new, homeologous template. These results suggest that the common practice of defining structural variants by the precise number of base pairs at their junction mis-represents the way apparent microhomology-mediated recombination events occur.

## Results

### Isolation of DSB repair-associated mutations

We isolated several independent sets of >1000 mutations arising after induction of *MAT*α switching using the transcriptionally silenced *HMR::Kl-URA3* donor. Mutations were selected as independent Ura^-^ (5-FOA-resistant) colonies after galactose-induced recombination (**Figure 1b**; see Materials and Methods). As noted above, ICTS events create chimeric *Kl-Sc-Kl URA3* genes that are generally in-frame and functional. To obtain sufficient numbers of ICTS events in this context, we created a -1 frameshift mutation and a single base pair change in *Sc-ura3-52* (designated *Sc-ura3-52\**) so that any gene conversion that copied the middle segment of this donor template would create a *ura3 Kl-Sc-Kl* chimera after switching to *MAT*. ICTS events that copied this region would then be readily recovered along with other 5-FOA-resistant outcomes such as base pair substitutions, -1 frameshifts, intragenic deletions and other events (**Figure 1c**). PacBio sequencing of the PCR-amplified *MAT* locus detected thousands of independent FOA-resistant mutations that grew as 5-FOA-resistant colonies, revealing the spectrum of mutations arising during repair replication (**<u>Table S1</u>** and **Figure 1d**). We performed this experiment with three independent biological replicates (annotated as sets 1-3; the second biological replicate also comprised two technical replicates that we averaged to calculate metrics for set 2). While set 1 had more reads than sets 2 and 3, all sets largely showed the same spectrum of mutations.

Overall, these results agree with those obtained previously by sequencing individual colonies selected for mutational events (5,6). Approximately half of the Ura^-^ colonies arose from base-pair substitutions and 30% were -1 frameshifts, the great majority of which were in homonucleotide runs. About 10% of mutations were +1 frameshifts, which were also more common in homonucleotide runs. A small fraction consisted of ∼300 bp insertions that proved to be sigma sequences that are the LTRs of yeast retrotransposon Ty elements. About 10% of mutations were ICTS events.

### Repair replication leads to many more -1 than +1 frameshifts, in contrast to normal replication

As we had observed previously, a frequent class of mutations arising during DSB repair were -1 frameshifts, 94% of which were in homonucleotide runs. In contrast, +1 insertions were much less frequent, and 83% occurred in homonucleotide runs (Fisher’s test; p-value < 2.2e-16). Among mutations in homonucleotide runs, -1s were found 3 times more often than +1s. To compare these results to the spectrum of +1 and -1 frameshifts among spontaneous mutations, we analyzed the whole-genome sequencing results obtained by Liu and Zhang (**<u>Table S2</u>**) (8). Among the spontaneous mutations, 95% of all +1 and -1 mutations arose in homonucleotide runs (binomial test; p-value < 2.2e-16). Moreover, during replication, the median number of +1 insertions in homonucleotide runs was only one to two times the median number of -1 deletions in such runs (“YPD” and “YPL” **Figure 2a**). The strong bias towards -1 frameshifts in HO-induced DNA repair synthesis (**Figure 2a**) most likely reflects intrinsic differences in the DNA replication machinery present during normal chromosome duplication and which is assembled to copy donor sequences during DSB repair; however, it is also possible that the difference arises from the apparent lack of mismatch repair proofreading during DSB repair (5). It is important to note that -1s in homonucleotide runs were distributed uniformly across the *URA3* sequence, as were SNVs in homonucleotide runs (**Figure 2b-c, S1a-b**).

**Figure 2:**
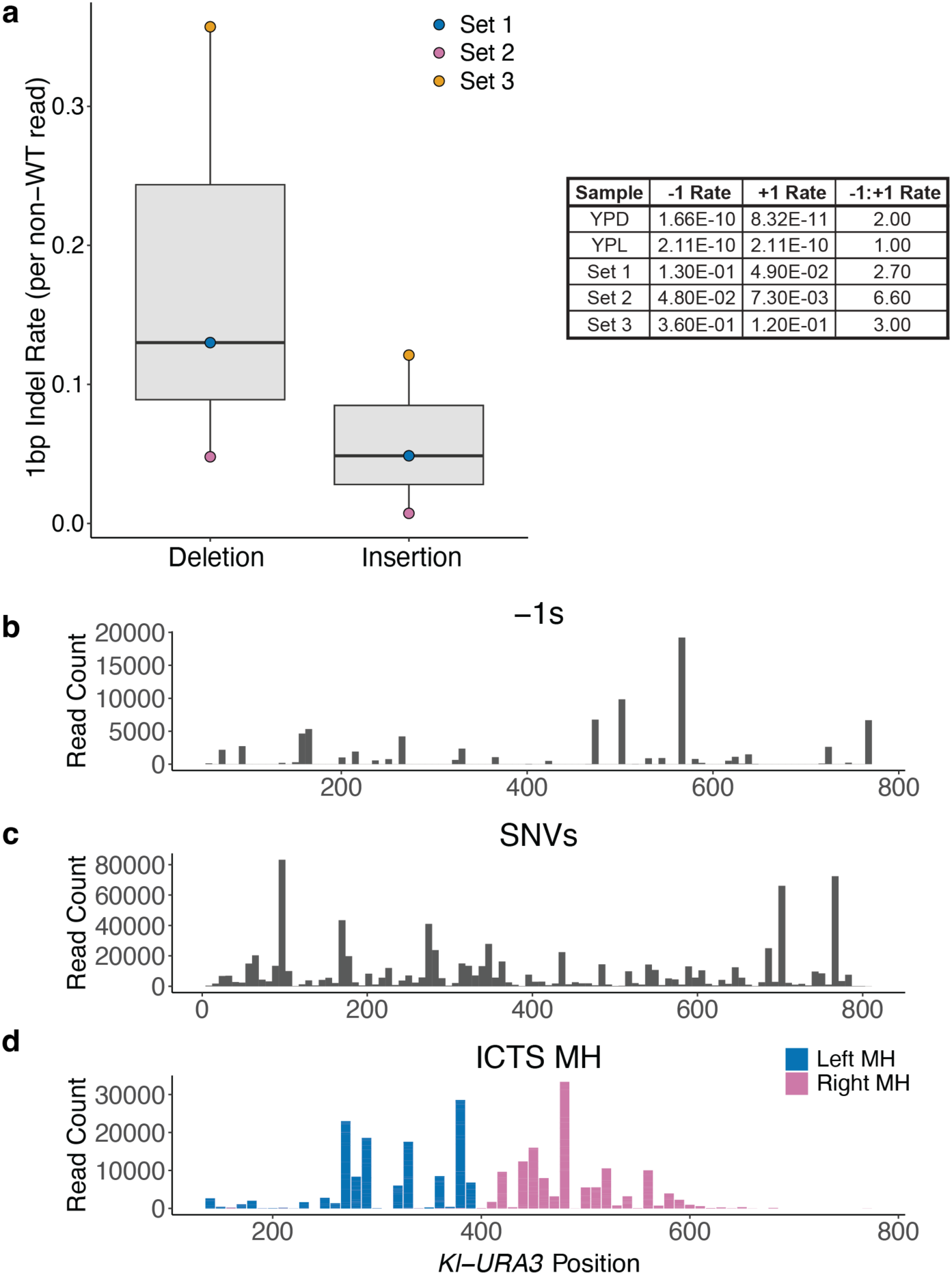
-1 indels are more common in repair replication than normal replication. **a**, Rate of 1bp insertions and deletions during normal and repair replication. The plot represents the rates of 1bp indels in sets 1, 2, and 3 during repair replication. The table includes the rates and ratios of +1:-1 indels in these three sets as well as in normal replication on YPD and YPL as measured by Liu and Zhang (8). Rates in our sets 1, 2, and 3 are per non-WT read. Rates in the Liu and Zhang data (YPD and YPL) are per nucleotide per cell division. **b** and **c**, The read counts of -1 IDs occurring within 3 bp homonucleotide runs (**b**) or SNVs (**c**) summed across all three sets. **d**, The read counts of MHs used in ICTS events were plotted vs the gene coordinate summed across all three sets. Note that the Ty retrotransposon insertion that prevents *Sc-ura3-52* expression removes the first 115 bp of homology between *HMR-Kl-URA3* and *Sc-ura3-52*.

### Interchromosomal template switches are frequent in replication repair, even with a highly divergent donor

Even though the *Sc-ura3-52\** sequence is only 72% identical to the *HMR::Kl-URA3* template that is initially engaged by the DSB end, 10% of all the Ura^-^ events we recovered were ICTS events, >99.5% of which copied the introduced -1 mutation and base substitution into the chimeric sequence. The few ICTS events that did not include this region displayed some other frameshift alteration that accounted for the Ura^-^ phenotype (**<u>Table S1</u>**).

Overall, the start and end of ICTS events were constrained to the middle of the *Kl-URA3* sequence, occurring primarily within about 150 bp from the -1 frameshift that defines *Sc-ura3-52\** (**Figure 2d, S1c**). On average, ICTS events included 165 bp copied from *Sc-ura3-52\** (**Figure 3a**). Each of the “jumps” between *Kl-URA3* and *Sc-ura3-52\** was bounded by a region of microhomology (MH) (on average 7.69 bp) shared between *Kl-URA3* and *Sc-ura3-52\** (**Figure 3b**). The most frequently used MHs tended to be the longest MHs on either side of the -1 mutation (**Figure 3c-e, S2, S3**), with closer MHs favored (**Figure 3f**). However, some very short (2 bp) MHs were used much more often than would be expected based on their length. Similarly, some long MHs (e.g. an 11-bp sequence underlined in **Figure 3c**) were used 10 times less often than many 6-bp MHs (**Figure 3e**). We note that we define MH usage by the standard practice in the field, counting only the number of matches at each *Kl-URA3/Sc-ura3-52\** junction, ignoring the contribution of additional homeologous base-pairing further from the junction. As noted above, the distribution of SNVs and -1 frameshifts in homonucleotide runs are not constrained; they are found across the entire *Kl-URA3* sequence (**Figure 2b-c** and **S1a-b**).

**Figure 3:**
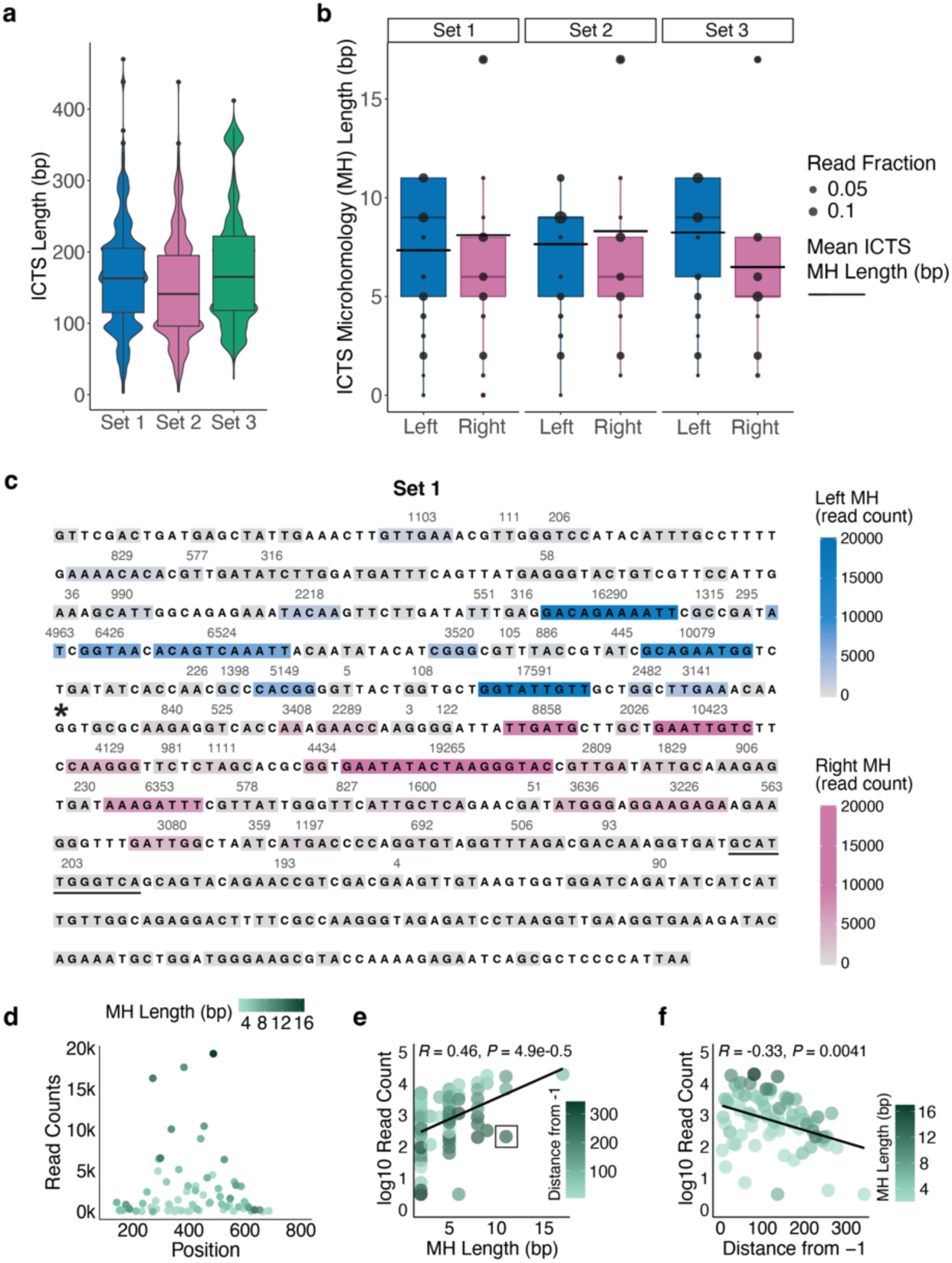
Microhomology used in ICTS events. **a**, The length distribution of ICTS events observed during replication repair. **b**, The length distribution of MHs flanking ICTS events. Right MH represents the jump from *Kl-URA3* to *Sc-ura3-52\** while left MH shows the jump back to the original *Kl-URA3* template. The bottom edge, central line, and top edge of the boxplots represent the first quartile, median, and third quartile of the data, respectively. **c**, The frequency with which each MH between *Kl-URA3* and *Sc-ura3-52\** was used for ICTS events. Grey boxes indicate unused MHs, blue boxes indicate MHs used to jump “into” *Sc-ura3-52\**, and pink boxes indicate MHs used to jump “out” of *Sc-ura3-52\**. Darker colors indicate more frequent usage. An 11-bp MH used much less frequently than several 6-bp MHs is underlined. The black asterisk indicates the introduced -1 frameshift in *Sc-ura3-52\**. *Sc-ura3-52\** lacks the first 115 bp of homology with *Kl-URA3*. **d**, MH position along *Kl-URA3* versus its usage in ICTS events, measured in read counts. Color scale indicates MH length in bp. **e**, Linear regression of ICTS MH length versus its usage in logged read counts. Black box highlights 11-bp MH used far less frequently than some 2-bp MH sequences. Rho and p-value calculated with Spearman’s rank-order correlation. **f,** Linear regression of a MH’s distance from -1 in *Sc-ura3-52\** and its usage in ICTS events. Rho and p-value calculated with Spearman’s rank-order correlation.

We formed two hypotheses to explain the paucity of ICTS events beginning closer to the start of repair replication (at the 3’ end of the *Kl-URA3* sequence). The first is that initiation of template switching is determined by topological stress during the copying to the *Kl-URA3* template that promotes the dissociation of the partly-synthesized DNA. Thus, there would be few jumps to microhomologies near the 3’ end. The second potential explanation is that the jumps are not simply between microhomologies *per se* but require substantial base-pairing between the partly-copied new strand and *Sc-ura3-52\** sequences (**Figure 4a-d**). The initiation of template switching would therefore be determined by the length of homeologous sequence copied and available for annealing to a new template.

**Figure 4:**
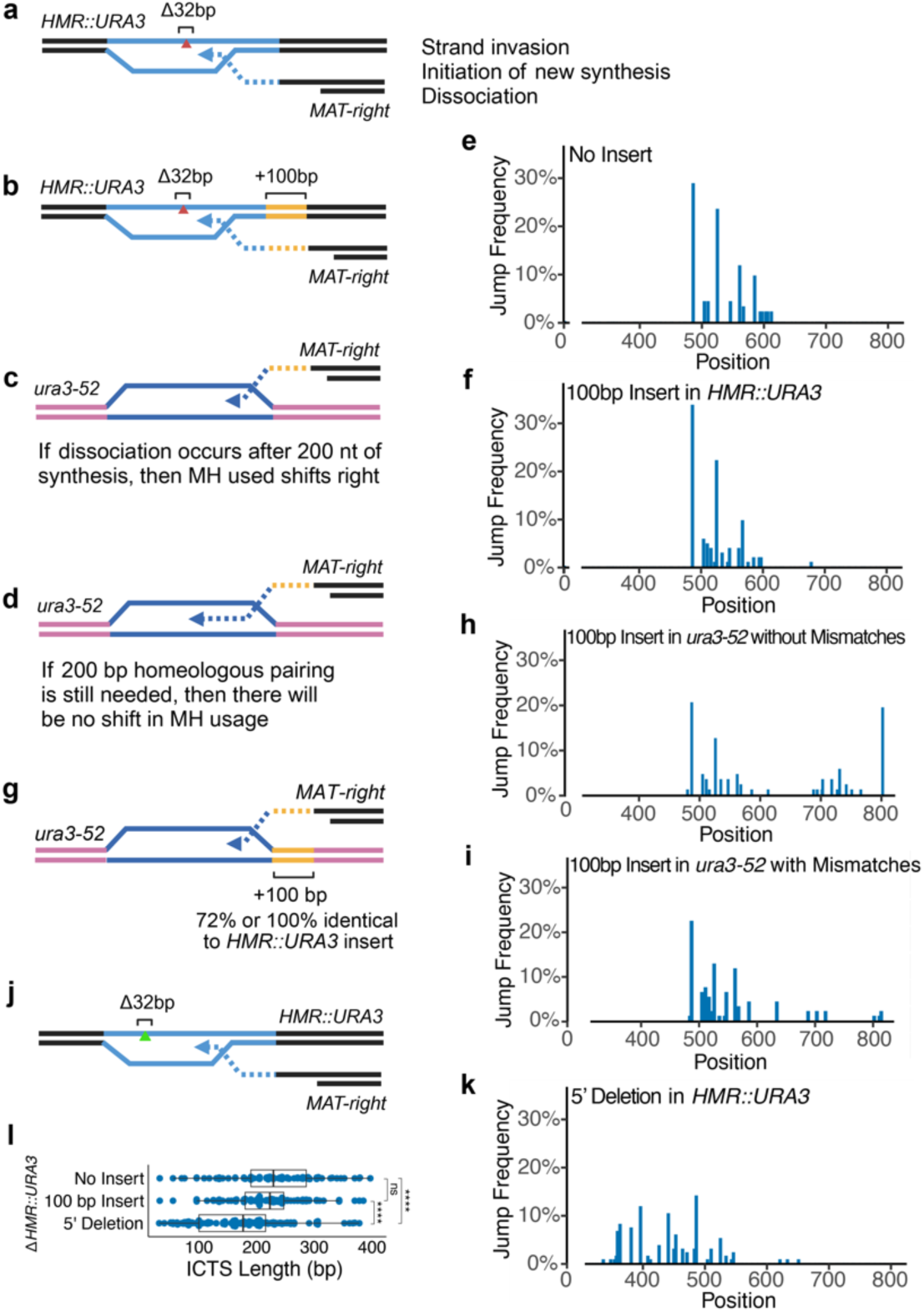
ICTS events require pairing ∼200 bp of homeologous sequence with *Sc-ura3-52\** before initiating new DNA synthesis. **a**, Diagram of strand invading into *Kl-URA3* (blue) at *HMR* and also dissociating from the template. **b**, Strand invasion and dissociation as in **a** in a strain that contains an extra 100-bp of unrelated DNA (yellow). **c.** If the nascent strand in **b** dissociates after 200 bp has been synthesized (**b**), then the MH usage will be shifted 100-bp to the right. **d.** If homeologous pairing is important, then there will be no shift in the MH usage. **e** and **f.** Comparison of template jumps into *Sc-ura3-52\** in the original strain (**e**) and in a strain with 100-bp of extra DNA (**f**) as diagrammed in panels **a** and **b**. Graphs plot the MH usage as a percentage of ICTS events vs. their location in the *Kl-URA3* open reading frame. A p-value of 0.77 was calculated from a KS-test comparing these two distributions. **g.** ICTS in strains with either the same 100 bp as in **b** or a homeologous version (72% identical) inserted adjacent to *Sc-ura3-52\**. **h** and **i.** Comparison of 3’ template jumps in the strains with an insert either 100% (**h**) or 72% (**i**) identical to that in **b** adjacent to *Sc-ura3-52\**. **k.** Locations of 3’ template jumps into *Sc-ura3-52\** in the strain diagrammed in **j**, where the 32-bp deletion was moved 109 bp 5’ of the 32-bp deletion in the other two strains. **l**, Distribution of ICTS gene conversion tract lengths for each strain. Wilcoxon Rank Sum test; **** indicates p-values are < 2.2e-16. P-values were corrected for multiple hypotheses using the Benjamini-Hochberg procedure.

To distinguish between these hypotheses, we compared two strains: one where the copy of *Kl-URA3* in *HMR* contains a 32-bp deletion and another containing this deletion as well as 100 bp of a randomly-generated DNA sequence inserted between the end of *Kl-URA3* and the start of *HMR-Z*, where strand invasion occurs (**Figure 4a-b**). Both strains carry *Sc-ura3-52* so that ICTS will produce Ura^+^ recombinants (6). In the latter strain an additional 100 bp of DNA would need to be synthesized before encountering the *Kl-URA3* sequences. In the topological stress and dissociation model, these added sequences would cause the MH usage to shift 100 bp to the right. Instead, we found that the spectrum of MH-usage did not differ between these two strains, suggesting that MH usage is not dependent on the extent of DNA synthesis at *Kl-URA3*. Rather, our results indicate that this process is strongly homology-dependent suggesting that at least 200 bp of homeologous DNA is needed to align the two 72% identical *Kl* and *Sc ura3* sequences (**Figure 4c-f**).

We then created two more strains, in which an additional 100 bp sequence was inserted in the *Sc-ura3-52* gene, such that the additional 100 bp copied at *HMR::Kl-URA3* could pair with sequences at *Sc-ura3-52*. In one case the sequences were identical to those inserted at *Kl-URA3*, while in the second case these sequences were about 72% identical – the same level of homeology seen between *Kl-URA3* and *Sc-URA3* (**Figure 4g**). In both these cases we saw, for the first time, ICTS events that could begin in the last 200 bp of the *Sc-ura3-52* open reading frame, strongly supporting the idea that what appear to be microhomology-mediated events actually require extensive adjacent base-pairing. Nevertheless, the preference for joining at longer microhomologies is still evident (**Figure 4h-i**). It is also possible that some events initiated in the first 200 bp will have jumped back before encompassing the 32 bp needed to create a Ura^+^ outcome.

We also examined apparent microhomology use in an additional strain, in which a 32-bp deletion in *Kl-URA3* was created 109 bp more 5’. Here, we again found that there were almost no ICTS events in which the initial jump to *Sc-ura3-52* occurred in the last 200 bp, but now MH usage ranged over an additional 100 bp (**Figure 4j-k**). The average length of the ICTS gene conversion tracts were also comparable across the three conditions, ranging from 172 bp in the strain with the 5’-shifted deletion to 231 bp in the case of the 3’-ward deletion and 220 bp in the strain with the latter deletion and 100 bp insert. While some of these length distributions are statistically different from each other, the magnitude of those differences (47 and 53 bp between median lengths) is small (**Figure 4l**). Taken together, these data argue that ICTS events, which have been called microhomology-mediated recombination (9, 10), are not in fact driven solely by base-pairing at often very short microhomologies, but depend on extensive (∼200 bp) alignment of the newly copied *Kl-URA3* strand with homeologous sequences at *Sc-ura3-52*.

### Longer flanking homology sequences increase the frequency of interchromosomal template switching

As a further characterization of the need for substantial base pairing to effect ICTS, we created a series of additional strains in which *Sc-ura3-52* was deleted and different-sized segments of *Kl-URA3* were inserted in four different locations (**Figure 5a-c**). Here we compared the need for adjacent *homologous* sequences when both *HMR-Kl-ura3-*Δ*32* and the second template have *Kl-URA3* sequences. The four constructs have the 32 bp region which is deleted in the *HMR-Kl-URA3* donor surrounded by 50, 100, or 150 base pairs on each side or the full length (but promoterless) *Kl-URA3* ORF (452 and 320 bp on each side of the deletion). Each of these constructs was inserted into four different chromosomes. Three of these sites (Chr 6, 7, and 16) had predicted 3D proximities to *MAT* and to *HMR* comparable to *URA3*, while the site on Chr 8 was predicted to be significantly closer to *MAT* and *HMR* (11). After *GAL::HO* induction, we measured the frequency of Ura^+^ colonies, which select for ICTS. As shown in **Figure 5d**, ICTS frequencies ranged from 1.5 x 10^-5^ to 3.0 x 10^-4^, the average of 2.5 x 10^-4^ being comparable to the original strain carrying *Sc-ura3-52* on Chr 5 (2.0 x 10^-4^) (6).

**Figure 5:**
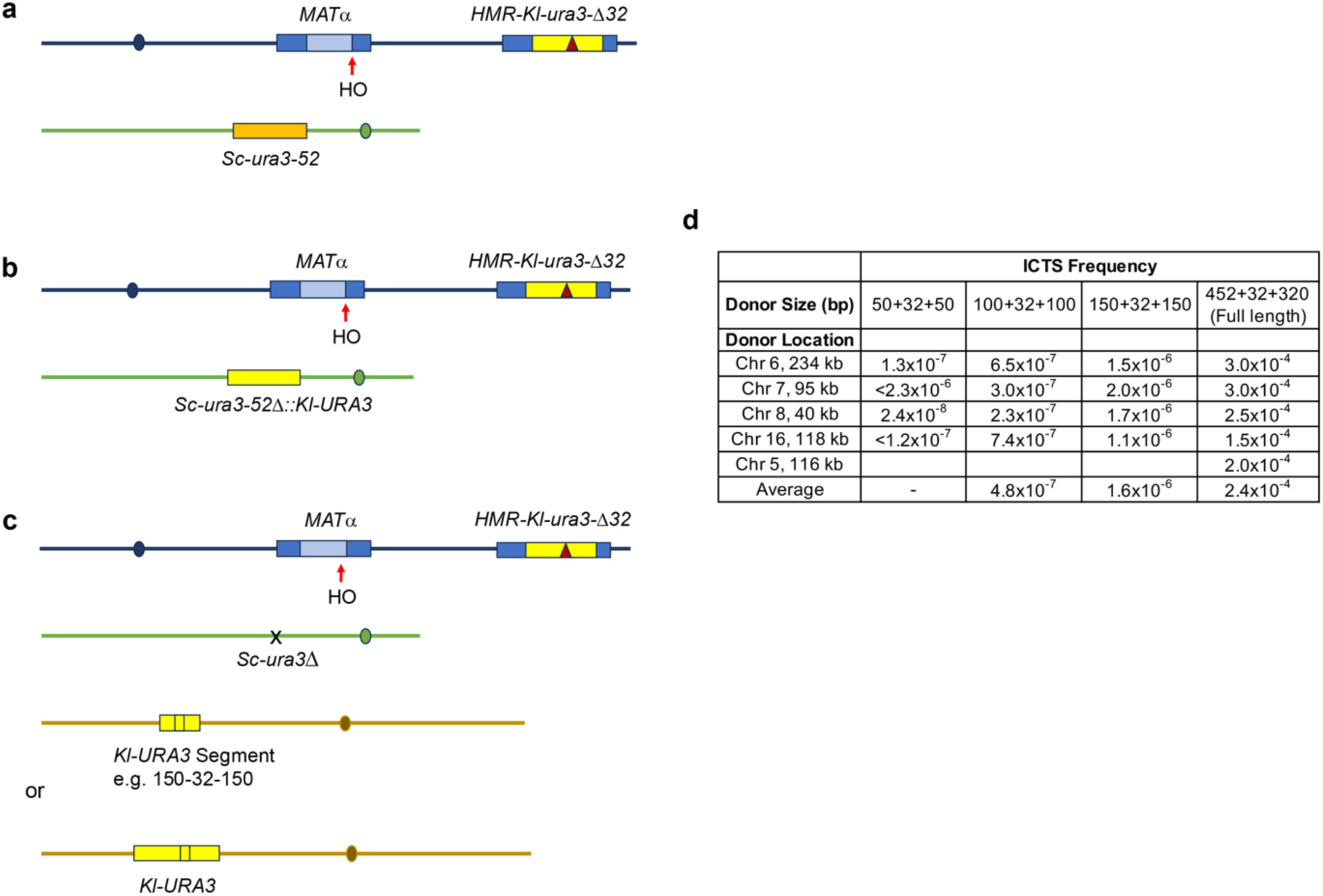
Effect of homology length on ICTS. ICTS induced by HO cleavage at *MAT*a was assayed (**a**), with *Sc-ura3-52* on Chr 5, (**b**), when the *Sc-ura3-52* donor on Chr 5 was replaced by a promoterless *Kl-URA3* or (**c**), when *Sc-ura3-52* on Chr 5 was fully deleted and either the full-length *Kl-URA3* or segments in which the 32-bp region is flanked 50, 100 or 150 bp (shown) of *Kl-URA3* sequence were inserted onto a different chromosome (Chr 6, 7, 8, or 16). **d**, The frequency of ICTS in each of these strains is shown. No average is given for the 50+32+50 case, as two samples were only determined to be less than the number of cells plated.

Compared to the rates of ICTS when there was >300 bp of homology on either side of the 32-bp deletion, strains with reduced flanking homologies were significantly lower. With 150 bp on each side of the 32 bp, the rates of ICTS averaged 1.6 x 10^-6^, i.e. more than 100-fold lower than ICTS rates with the full-length sequence. With 100 bp on either side, the rates fell to an average of 4.8 x 10^-7^. With 50 bp flanking the necessary 32 bp, we failed to recover ICTS in two strains (Chr 7 and 16) among > 2 x10^7^ cells plated and the rate for the other two strains was 2.5 x 10^-7^. Given that the average ICTS gene conversion tract length for correcting the 32 bp deletions ranged from 231 bp (6) to 220 bp (**Figure 4l**), (i.e. about 100 bp on either side of the necessary 32 bp), we might expect that having 150 bp on each side (a total of 332 bp including the necessary 32 bp) would be sufficient to capture a significant fraction of ICTS, but this is not the case. These data support the idea that efficient ICTS depends on aligning 200 or more bp of homologous or homeologous sequences on at least the “invading” side of the template jump.

### Microhomology-bounded intragenic deletions are more abundant than intragenic tandem duplications

Within the copied *MAT::Kl-URA3* sequences we identified both MH-bound intragenic deletions (IDs) and intragenic tandem duplications (TDs). IDs were approximately 12 times more abundant than TDs (**Figure 1d**) and were on average 56 bp long. In contrast, TDs were quite short, averaging only 22 bp (**Figure 6a**). Both IDs and TDs were usually bounded by MHs: across all three sets, IDs had a mean MH length of 6 bp while TDs had mean MH length of 3 bp (**Figure 6b**). These characteristics suggest a picture of the repair replication fork in which the polymerase can jump further forward than backward when it dissociates (**Figure 6c**, see Discussion).

**Figure 6:**
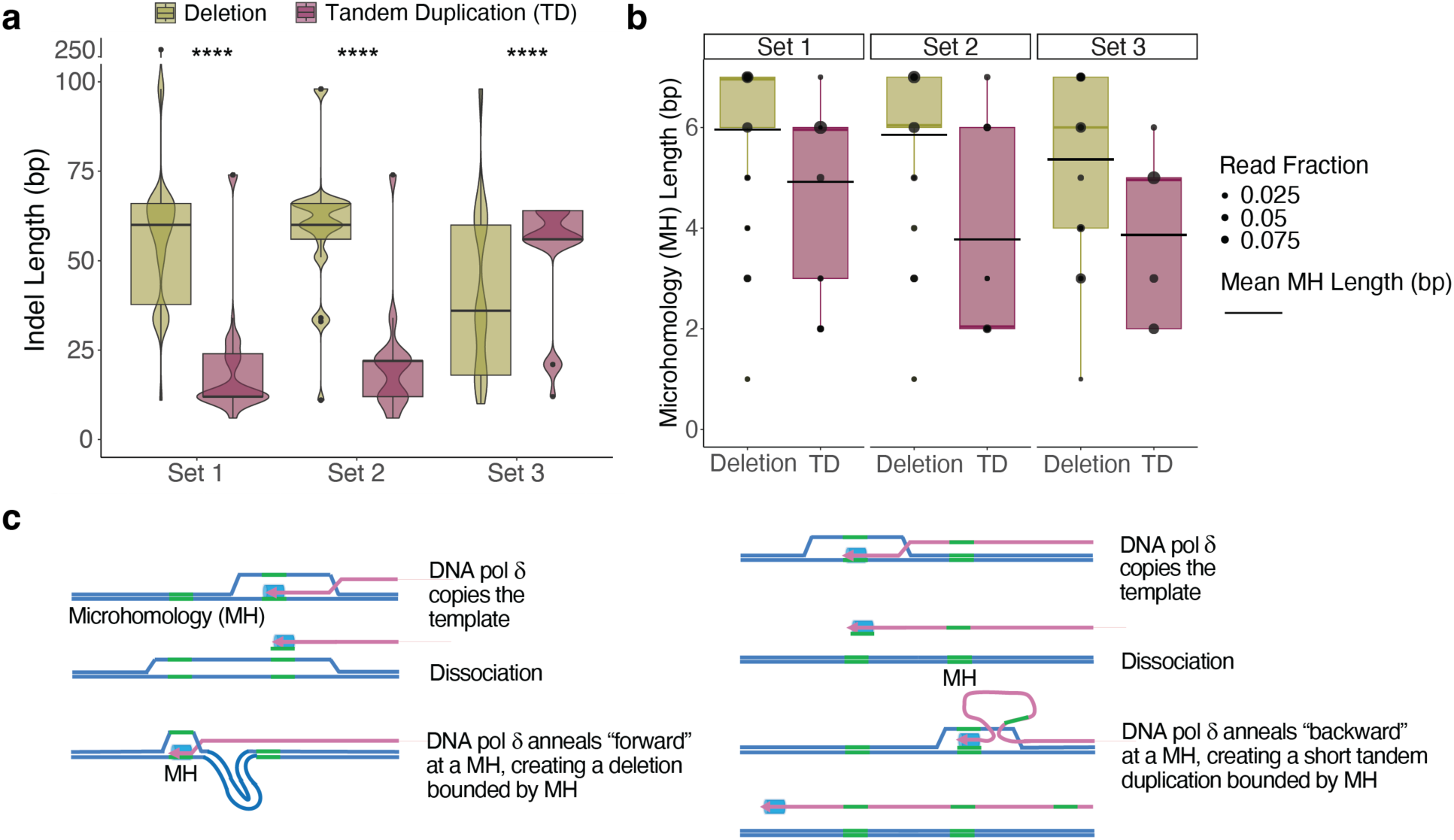
Microhomology-bounded intragenic deletions are longer than tandem duplications. **a**, The length distribution of MH-bounded deletions and tandem duplications in repair replication. Deletions are significantly longer than tandem duplications, Wilcoxon Rank Sum test. **** indicates p-values are < 2.2e-16. Several exceptionally long deletions in Set 3 greater than 400 bp amounting to 2% of its deletion reads were omitted from the figure but included in the statistical test. **b**, Distribution of MH length for MH-bounded deletions and tandem duplications. The bottom edge, central line, and top edge of the boxplots represent the first quartile, median, and third quartile of the data, respectively. **c**, Schematic of the mechanism of formation of MH-bounded deletions (left) and tandem duplications (right).

### Microhomology-bounded intragenic deletions are not aided by additional adjacent microhomeology

We next set out to further characterize the nature of IDs arising during the copying of *HMR::Kl-URA3* into *MAT*. Using the *Sc-ura3-52\** strain, we isolated and pooled 50,000 FOA-resistant colonies and purified DNA. We then extensively digested the DNA with the *Bst*EII and *Fsp*I restriction endonucleases that each cleave one site near each other in the *Kl-URA3* sequences and do not cleave in *ura3-52*. Thus only mutational events that remove the *Bst*EII and *Fsp*I sites, such as ICTS events and IDs, should remain. We sequenced the digested DNA by PacBio sequencing. The vast majority of events identified in this sample were ICTS events, while 2% were IDs (**Figure 7a**). One-bp indels in homonucleotide runs likely arose from residual undigested DNA. Similar to sets 1, 2, and 3 (**Figures 3c, S2, S3**), ICTS events most frequently used the longest MHs on either side of the -1 deletion that allowed selection of these events (**Figure S4**). ICTS events were on average 166 bp long and used MHs averaging 7.4 bp (**Figure 7b)**. IDs were tightly distributed around a mean length of 62 bp, with average MH of 6 bp (**Figure 7c**).

**Figure 7:**
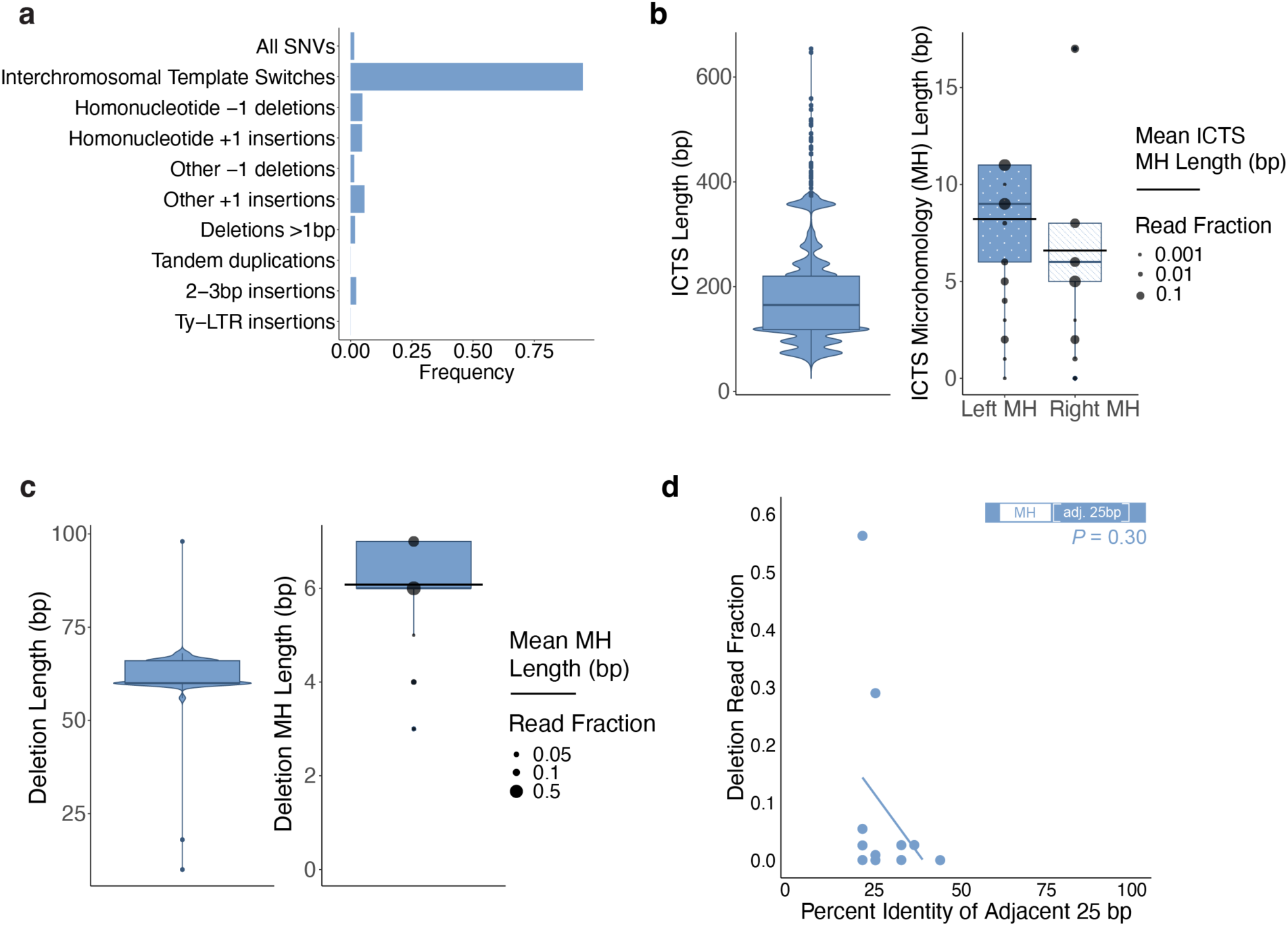
Adjacent homeology does not contribute to the choice of MH used for intragenic deletions or ICTS events. **a**, Spectrum of mutations observed with *BstE*II and *Fsp*I digestion. **b**, ICTS events centered around 177 bp in length and used an average of 8.2 bp of MH on the left and 6.6 bp on the right. The bottom edge, central line, and top edge of the boxplots represent the first quartile, median, and third quartile of the data, respectively. **c**, Intragenic deletions averaged 62 bp in length and used an average of 6 bp of MH. The bottom edge, central line, and top edge of the boxplots represent the first quartile, median, and third quartile of the data, respectively. **d**, Comparison of frequency of specific MH usage and neighboring homeology across deletions identified with *BstE*II and *Fsp*I digestion. A linear regression was fit to the read fractions vs. percent homeology values of the 25 bp adjacent to the MH shared between *Kl-URA3* and *Sc-ura3-52\**. A one-sample two-tailed t-test calculates the probability that the slope of the line is different from 0.

We entertained the hypothesis that the IDs used most frequently shared more adjacent homeology that could assist in stabilizing the intermediate leading to a deletion. However, when we calculated the percent base identity for the 25 bases upstream of the MH in each intragenic deletion event, we observed no significant association between adjacent homeology and frequency of MH usage (**Figure 7d**). A similar analysis of the 25 bp adjacent to ICTS MHs suggests that after ∼200 bp of homeologous sequence has been copied and used for alignment, local microhomologies dictate the exact location of the junction between the divergent sequences (**Figure S5**). We suggest that the 3’ end of the aligned sequence is trimmed back to a well-base-paired junction, likely by the 3’ to 5’ exonuclease proofreading activity of DNA polymerase δ, as we have shown in break-induced replication between mismatched substrates (12).

## Discussion

By sequencing a large number of structural variants arising as mutations during DSB repair, we have identified several aspects of the repair process that are distinctive for repair-induced mutations compared to those arising during simple replication of the same sequences.

As we have shown previously, a large proportion of the mutation repair events involve some form of dissociation of the partly-copied initial strand. These dissociations enable the formation of IDs and TDs, quasipalindromes, and interchromosomal template switches. The simplest example of this “slippage” are -1 frameshifts in homonucleotide runs, which greatly outnumber +1 insertions. In normal replication -1 and +1 frameshifts occur at equal rates. This difference makes clear that the repair replication apparatus differs in distinct ways from the normal replication fork. In normal replication, leading and lagging-strand synthesis are coordinated and apparently closely associated with proofreading by the MSH-MLH mismatch repair system (13, 14). However, DSB repair using synthesis-dependent strand annealing occurs in two separate steps. A first strand is copied and only later – after second end capture – is the first strand used as a template to complete the repair event (15). Moreover, the mismatch repair machinery appears to play little or no role in correcting the errors arising during DSB repair (5), possibly because this proofreading apparatus does not follow closely behind the repair replication fork as it apparently does when there is a complete simultaneous leading- and lagging-strand replication fork. The many more -1 deletions arising during repair may reflect some intrinsic feature of DNA polymerase δ when it is not part of a coordinated replication fork; alternatively, the ratio of +1 and -1 events may be influenced during normal replication by the mismatch repair system. That DNA polymerase δ is the principal DNA polymerase used in repair is evident from the fact that a proofreading-defective mutation (*pol3-01*) of Pol δ dramatically reduces the appearance of all classes of structural variants in our repair system (5). We attribute the absence of the frequent slips and jumps to the observation that a proofreading-defective Pol δ is more “processive” and less likely to dissociate from its template, as demonstrated *in vitro* (16).

The pattern of MH-bound IDs and TDs that we observe suggests a picture of the repair replication fork (**Figure 8**). After the DSB end is resected and bound by Rad51, strand invasion occurs at the region of homology on the right side of *HMR::Kl-URA3*, creating a D-loop. In the *MAT* switching system, the presence of 700-bp nonhomology to the left of the HO cleavage site biases the initial strand invasion to use the *HMR-Z* region (17). The 3’ end of the invading strand then is used by DNA Pol δ to copy the donor, by a moving replication “bubble” in which the region ahead of the fork opened either by the movement of the polymerase itself or by a helicase; however, we have previously shown that gene conversion events such as *MAT* switching are not dependent on the normal replicative CMG helicase (18). We suggest that the D-loop is kept open by the binding of the single-strand DNA binding protein complex, RPA (18). An intrinsic feature of this repair process is that the newly-copied strand does not remain semi-conservatively bound to its template; rather, the strand is displaced so that eventually all the newly-copied DNA is found at the recipient locus. Inherent in this process, the previously copied region behind the fork soon “closes up.” Because of the displacement of the new strand, the repair replication fork is unstable; indeed, if a helicase driving the new-strand displacement is more active than the rate of polymerization, the newly-copied strand may be completely dissociated. Such dissociations are necessary for events such as ICTS, quasipalindrome mutagenesis, and both IDs and TDs.

**Figure 8:**
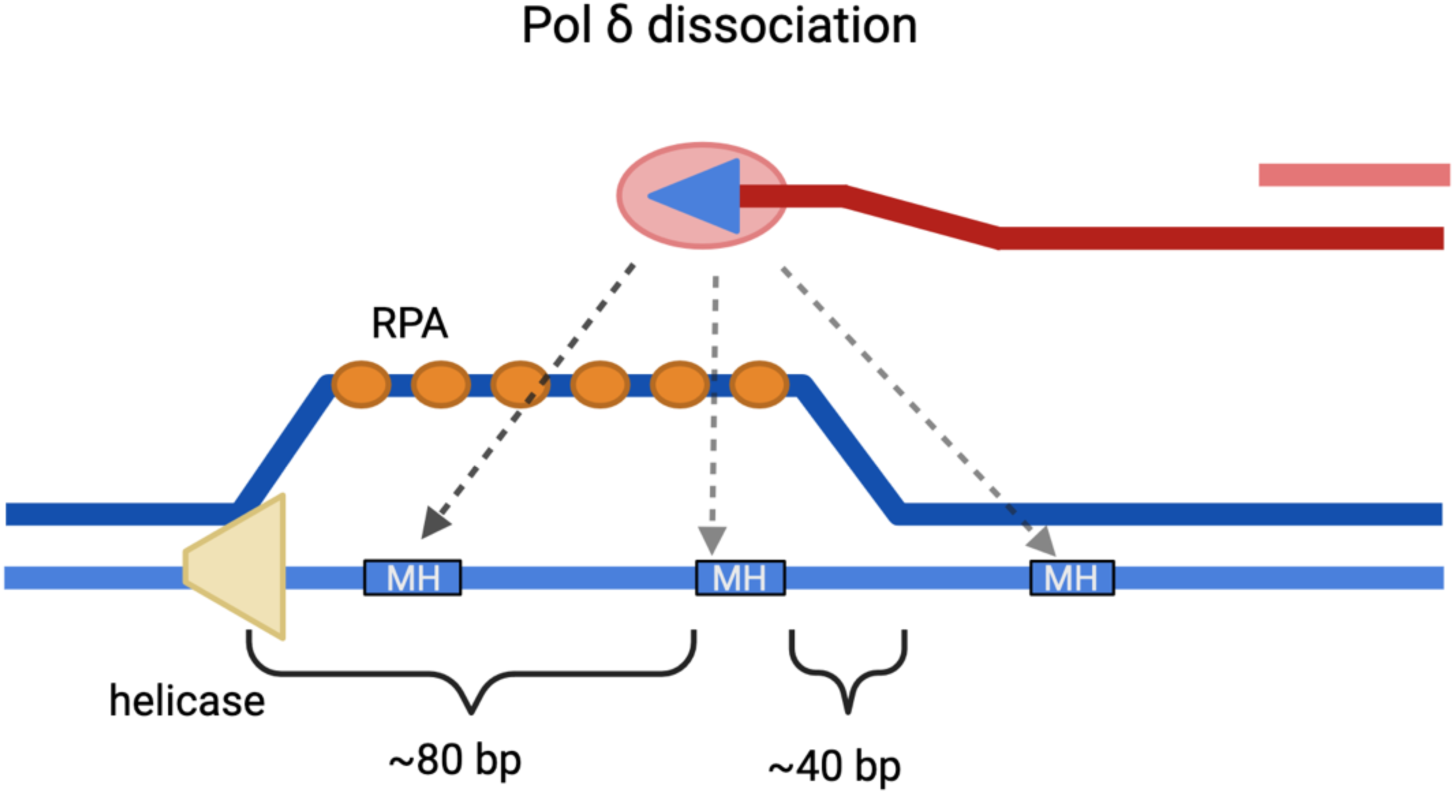
Schematic of the repair replication fork. A helicase precedes the migrating D-loop ∼80 bp ahead of the polymerase. The DNA re-anneals starting about 40 bp behind the polymerase. These values are based on the mean indel length plus one standard deviation. Whether re-invasion of the dissociated strand occurs still attached to DNA polymerase is not known.

Our results suggest that dissociation can result in a successful template switch only after ∼200 bp of homeologous sequence has been copied. The endpoints of the ICTS events show clearly that longer MHs are strongly favored, but the evidence also shows that significantly longer homology or homeology is needed to align the sequences. This suggests that the convention of categorizing structural variants by the number of precisely matching base pairs at their junction is overly simplistic and not fully representative of the repair mechanism involved. The length of breakpoint-adjacent sequence that appears to be important is consistent with sequence requirements for Rad51-mediated repair of HO-mediated events, where a minimum of 70-100 bp is needed but the efficiency improves up to about 1-2 kb (17,19-20). We suggest that, after the alignment of the dissociated, partly copied new strand with the homeologous *Sc-ura3* sequences, 3’ to 5’ exonuclease activity of Polδ or 3’ flap clipping by Rad1-Rad10 (12,21) will trim back the end preferentially to locations with the longer MHs. Dissociation may also be responsible even for the slippage in -1 frameshifts, although we observed these events uniformly across the entire length of *URA3* suggesting that factors other than homeology contribute to the formation of these mutations. Indeed, we find no evidence of any role of the use of homeology in the 25 bp adjacent to the MH used in IDs.

We observed that IDs are longer than TDs. We suggest that the creation of IDs or TDs requires that the template strand remain single-stranded and accessible to polymerase-mediated annealing either in front of the point of dissociation (creating an ID) or behind that point (creating a TD) (**Figure 6c**). The presence of RPA in the non-template strand should help maintain this open structure, but the sequences behind the fork will likely be “closed up” as the strand is dissociated, leaving only a very short region where a TD might be generated. This picture of the repair fork is supported by *in vitro* and *in vivo* mapping of the size of a D-loop generated by Rad51 in yeast (22).

Further refinement of this picture may be gained by examining the consequences of mutations that alter the processivity of DNA polymerase or various helicases as well as factors associated with loading and unloading PCNA and other replication factors. This investigation is ongoing.

Many aspects of this dissociation are not known. Does the strand become detached from the DNA polymerase? Do ICTS events require the participation of Rad51? Because Rad51 is required for the initial strand invasion, it is difficult to assess its role in secondary invasion events. Further experimentation will yield important insight into the mechanics of and participants in repair replication.

## Materials and Methods

### Yeast Strains and Plasmids

Strains used in this work are derivatives of WH50 (5) that were constructed by Cas9 mediated gene editing or by conventional transplacement strategies. Details can be found in the Supplemental Materials section. **<u>Table S3</u>** lists the strains and genotypes used in this work.

### Isolation of Mutations During DSB Repair

Gene conversion was initiated by plating cells onto medium containing galactose to induce *Gal::HO* and then isolating independent Ura^-^ cells using 5-FOA medium. Genomic DNA was prepared from pools of approximately 33,000, 1,100 or 50,000 Ura^-^ colonies, for libraries 1, 2, and 3 respectively. In addition, we enriched for intragenic deletion and ICTS events by digesting DNA from library 3 with *BstE*II and *Fsp*I which should eliminate most SNVs, -1 frameshifts and TDs. The *Kl-ura3* sequence was PCR-amplified using primers specific for the *MAT* locus and the products were submitted for NGS or PacBio sequencing. Preparations of these libraries are described in detail in the Supplemental section.

We induced Ura^+^ ICTS events in YCM1, tNS2921, tNS2922 or tNS2923 by replica-plating colonies to SD+galactose-uracil plates and isolating papillae growing within the imprints. The sequences were obtained from these colonies by PCR-amplifying the *Kl-ura3* sequence with *MAT*-specific primers and determining the sequences by Sanger sequencing.

### Homology Length Dependence

To determine the effect of homology length and chromosome location, *Kl-URA3* sequences were inserted in 4 chromosome locations by Cas9-mediated gene editing, using guide RNAs for Chr 6 (nt 233939-233958), Chr 7 (95180-95199), Chr 8 (39657-39676) and Chr 16 (118053-118072).

Segments of *Kl-URA3* sequence were inserted by using forward and reverse primers that had 32 bp of homology to the chromosome and 20 bp of homology to *Kl-ura3* appropriate for the size of each insert. Locations and sizes of inserts for strains yQW852 – yQW871 are shown in **Figure 5d**. Euclidean distances between each insertion site and *HMR* were determined by the methods of Duan *et al* (11). ICTS frequencies were determined after growing strains in YEP-raffinose overnight and then adding 2% galactose to logarithmically growing cultures for 4 h before plating on YEPD and uracil dropout plates.

### Alignment and mutation calling of *MAT::Kl-URA3* sequence from Ura^+^ colonies

Due to the presence of ICTS events, alignment to *Sc-URA3* was impossible with existing methods, preventing calling of mutations. Therefore, we created a custom alignment algorithm based on Smith-Waterman (23) and Breakpointer methods (24).

First, to determine if each sequence contains an ICTS event, they were aligned to *Kl-URA3* using a global Needleman-Wunsch algorithm using the following scores:

| Initial Alignment |  |
| --- | --- |
| Match | 1 |
| Mismatch | -80 |
| Gap open | -500 |
| Gap extend | -10 |

The score of this alignment was normalized to the length of the aligned query (as a proxy for highest possible score). Alignments with a normalized score higher than -2 were kept. Alignments with a lower normalized score were passed on to the modified alignment algorithm that allows jumping between *Kl-URA3* and *Sc-ura3-52.* All scores and thresholds were determined empirically. In all alignment matrices, the sequence to be aligned is along the top of the matrix (aka in the columns) and the reference sequence (*Kl-URA3* or *Sc-ura3-52\**) is along the side (aka in the rows).

The first step of the modified alignment algorithm is to do a global/local alignment of each sequence to *Kl-URA3*. To force inclusion of the left side of *Kl-URA3*, the alignment matrix is initialized as a Needleman-Wunsch matrix, with 0 only at [1,1], and gap penalties applied for all other [1,] and [,1] cells. Scores for this alignment, called M1, are set to optimally find the best matching left-most chunk of each sequence, so it has moderate mismatch score, but high gap scores:

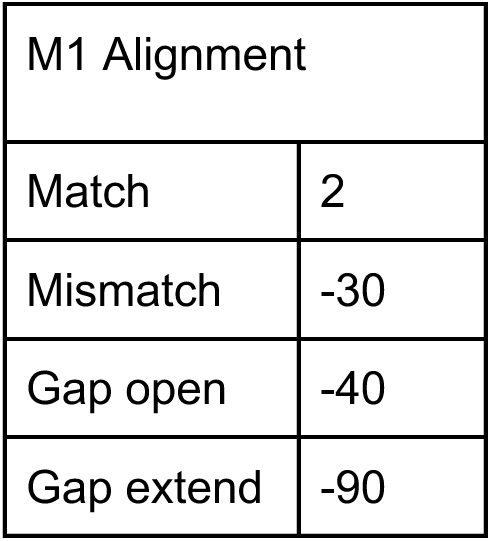

After finding the best left-most alignment between each ICTS-containing sequence and *Kl-URA3*, we perform a local Smith-Waterman alignment between each sequence and *Sc-ura3-52\**. Since this will be the middle of the final alignment it is not required to include all of the sequence, so the alignment matrix is initialized with 0s in [,1] and [1,]. Now mismatch, deletion and insertion scores are lowered to allow for some errors within the gene conversion. Additionally, this alignment, called M2, is also allowed to ‘jump’ back to the M1 alignment matrix (i.e., template switch to *Kl-URA3*). Specifically, at each cell of the alignment matrix, there is an option for the alignment to jump to the maximum cell in the previous column of M1. This corresponds to keeping the same index of the sequence but allowing a jump to anywhere in *Kl-URA3*. M2 does not allow for affine gaps.

| M2 Alignment |  |
| --- | --- |
| Match | 2 |
| Mismatch | -6 |
| Deletion | -20 |
| Insertion | -3 |
| Jump to M1 | -1 |

This step recovered sequences that were erroneously rejected due to the strictness of the cutoff for accepting the Initial alignment. These sequences lack ICTS events, so the M2 alignment will not extend very far beyond the M1 alignment. Therefore, any M2 alignments that extend M1 less than 15 bases are rejected in favor of the Initial alignment.

If the M2 alignment is accepted, the last step is to create an M3 alignment of the sequence to *Kl-URA3*, allowing jumping back to M2. To keep the location of the jump between M1 and M2, we first create a matrix to combine M1 and M2 into one alignment matrix to allow M3 to jump back to the previously completed Kl-Sc alignment. Because we are constructing a reference for this intermediate “new reference” alignment, this alignment is between the sequence and the aligned reference sequence from M2. This new alignment matrix needs to have a higher match score to encourage the third and final alignment to jump to it. That correspondingly requires increasing the other scores as well:

| MNewRef Alignment |  |
| --- | --- |
| Match | $805 = \text{nchar}(KI-URA3 + 1)$ |
| Mismatch | $-67 = -1 * \text{nchar}(KI-URA3) / 12$ |
| Deletion | $-844 = -1 * (\text{nchar}(KI-URA3) + 40)$ |
| Insertion | $-844 = -1 * (\text{nchar}(KI-URA3) + 40)$ |
| Jump to M2 | -1 |

This alignment is initialized to force inclusion of the left-most edge of the query and reference (initialized as a Needleman-Wunsch matrix, with 0 only at [1,1], and gap penalties applied for all other [1,] and [,1] cells). Traceback starts at the max cell in the bottom row, forcing inclusion of the entire M2 reference alignment.

The final alignment is then constructed with the M3 alignment between the sequence and *Kl-URA3*, allowing jumping to the MNewRef reference alignment (which is essentially the *Kl-URA3* and *Sc-ura3-52\** reference sequences pasted together at the first microhomology of each sequence’s ICTS event). This is a global alignment, specifically the M3 alignment matrix is initialized as a Needleman-Wunsch matrix, with 0 only at [1,1], and gap penalties applied for all other [1,] and [,1] cells and the traceback is initialized at the bottom right cell, to force inclusion of the entire reference and query sequence. Scores for this matrix are lower, to allow jumping back to MNewRef & tolerance of any SNVs:

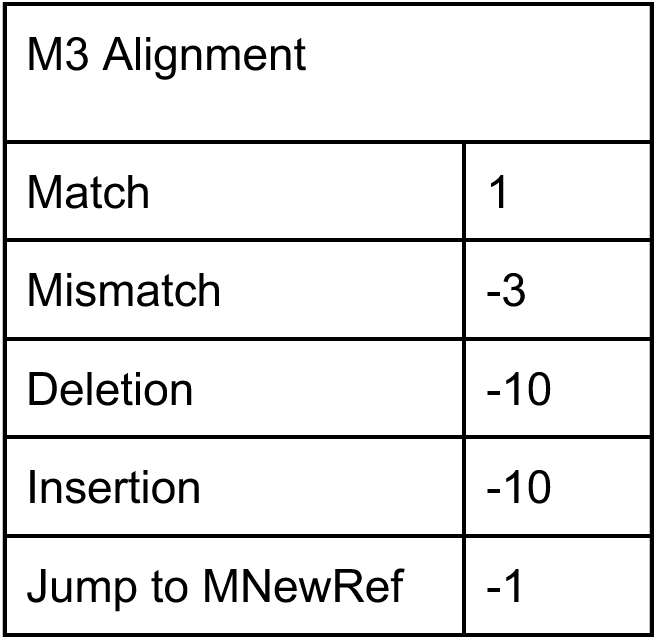

Mutations were identified by comparing between the aligned sequences resulting from this final alignment or the Initial alignment and the reference *Kl-URA3* and *Sc-ura3-52\** sequences.

### Statistical Methods

Analyses were performed in R version 4.2.1. Statistical tests are notated on figures and in figure legends when relevant.

## Acknowledgments

Initial DNA sequencing and analysis was performed by Dr. Alex Ferrazzoli, who passed away in 2020. We would like to thank Yuko Nakajima and Randall Tyler for their detailed advice and assistance in the preparation of the PacBio samples. Figures 1A-C and 7 were created with <u>BioRender.com</u>. Dr. Beroukhim was supported by the Gray Matters Brain Cancer Foundation, the Pediatric Brain Tumor Foundation, NCI grants R01 CA188228 and R01 CA262462, and Break Through Cancer. Research in the Haber lab is supported by NIH grant R35 GM127029. Summer research support for V.L. was through a Frederick W. Alt Biology Summer Research Fellowship. S.D. was supported by an NIH NCI NRSA award (1F32CA261024) and is currently supported by an NIH NIGMS career development award (K99/R00), 1K99GM155595.

## Author contributions

N.S., Q.W., T.C., V.L., L.T., C.M., and E.S. performed the recombination experiments. S.D., S.W., and S.Z. analyzed the data. J.E.H. and S.D. wrote the manuscript with input from all authors. S.D., S.W., and J.E.H. created the figures. S.D., R.B., and J.E.H. conceived the project and designed the experiments and analyses.

## Supplementary Figures & Supplementary Figure Legends

**Figure S1:**
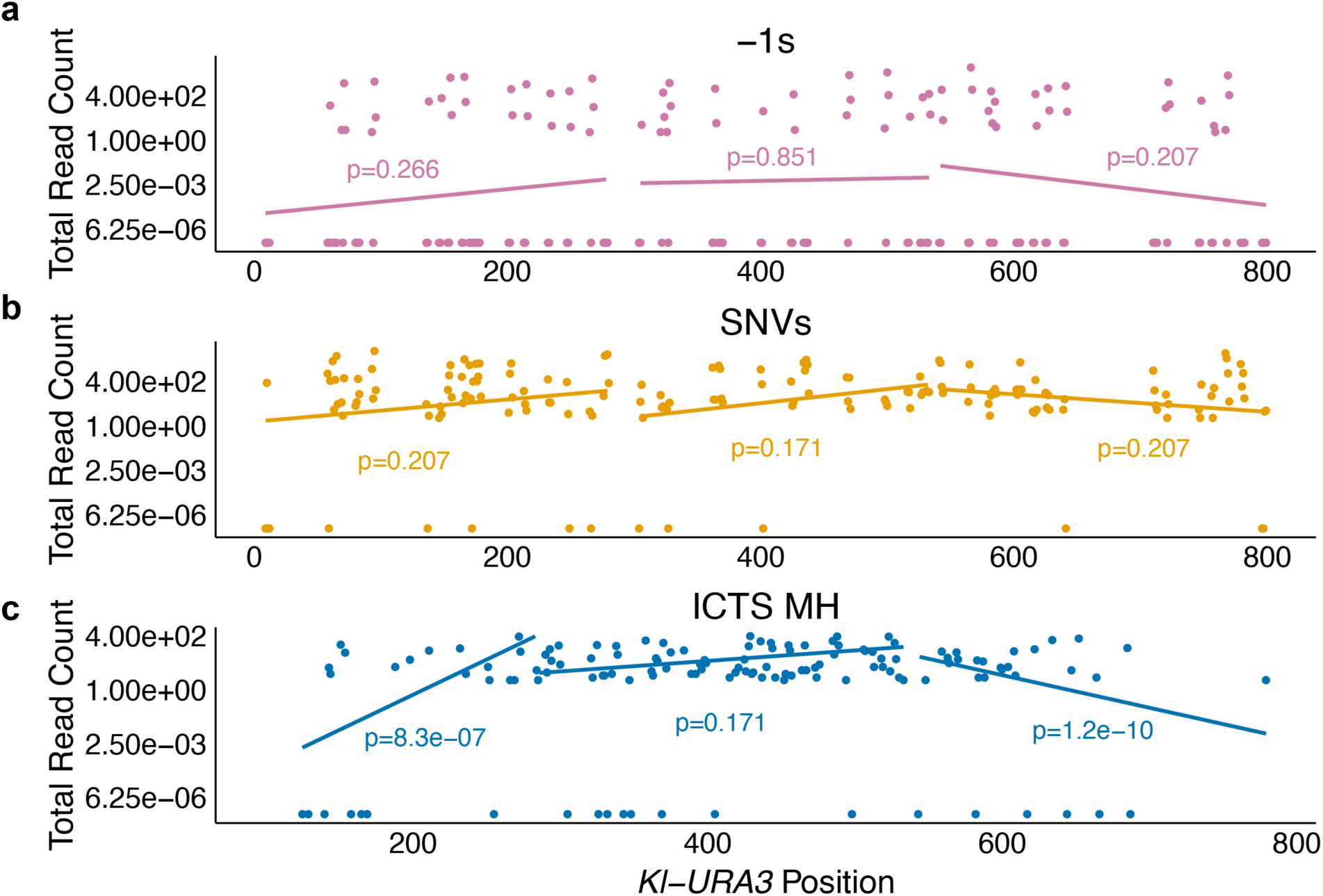
SNVs and -1s are uniformly distributed across *Kl-URA3*, while ICTS MH sequences are exclusive to the center. **a** and **b**, The read counts summed across all three sets of SNVs (**a**) or - 1 deletions (**b**) occurring within 3 bp homonucleotide runs. **c**, The read counts summed across all three sets of MHs used in the ICTS events were plotted vs the gene coordinate. Loci with no events are annotated with a read count of 0. The *Kl-URA3* sequence was split into thirds and a linear regression was fitted to the read counts of each event type within each third of the sequence. A one-sample two-tailed t-test calculates the probability that the slope of each line is different from 0. Annotated p-values were FDR-corrected. An annotated p-value less than 0.05 indicates that events of that type did not occur uniformly in the annotated region of *Kl-URA3*.

**Figure S2:**
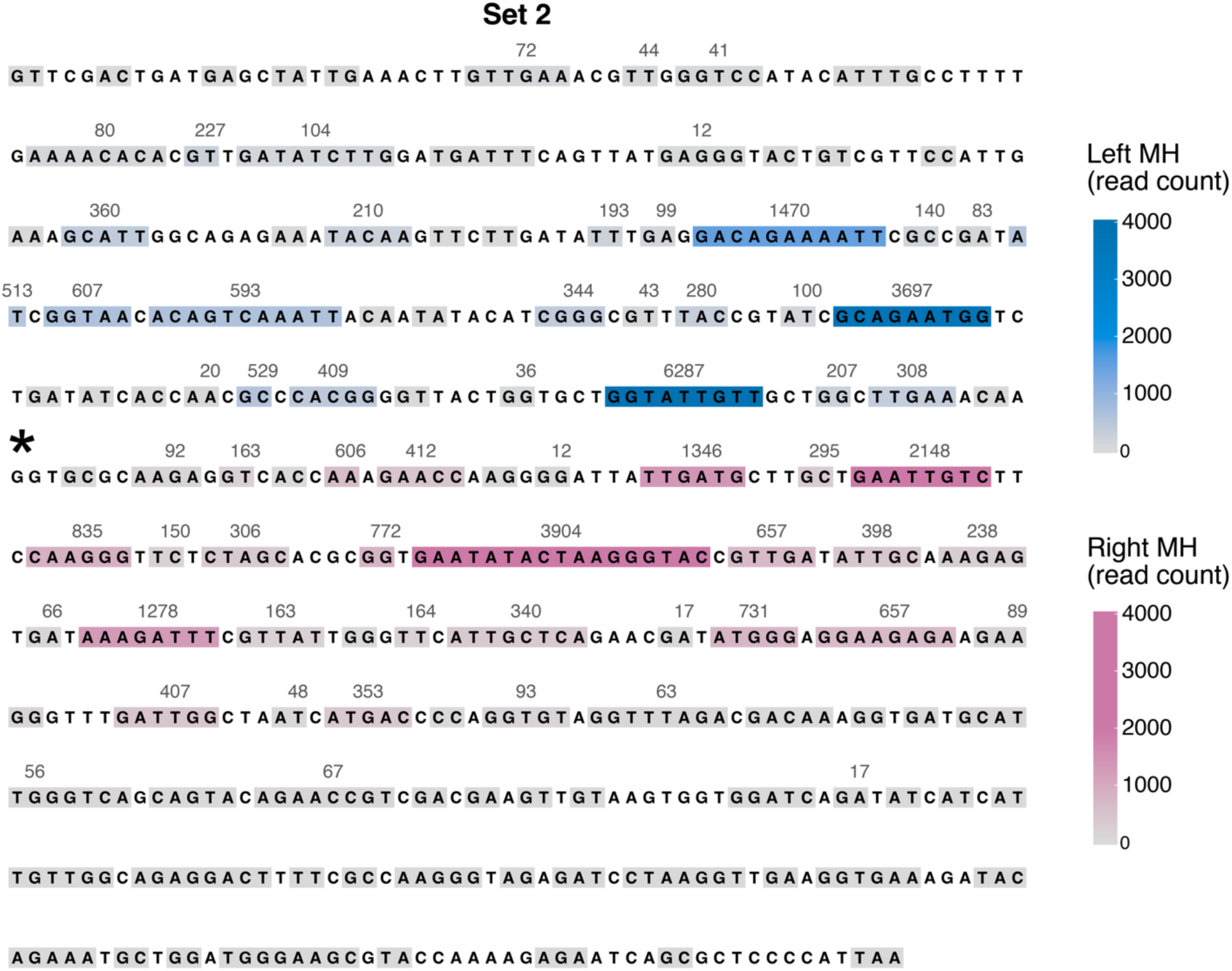
Frequency of usage of MHs between *Kl-URA3* and *Sc-ura3-52\** to form ICTS events in set 2. Grey boxes indicate unused MHs, blue boxes indicate MHs used to jump “in” to *Sc-ura3-52\** and pink boxes indicate MHs used to jump “out” of *Sc-ura3-52\**. Darker colors indicate more frequent usage. Black asterisk indicates the introduced -1 frameshift.

**Figure S3:**
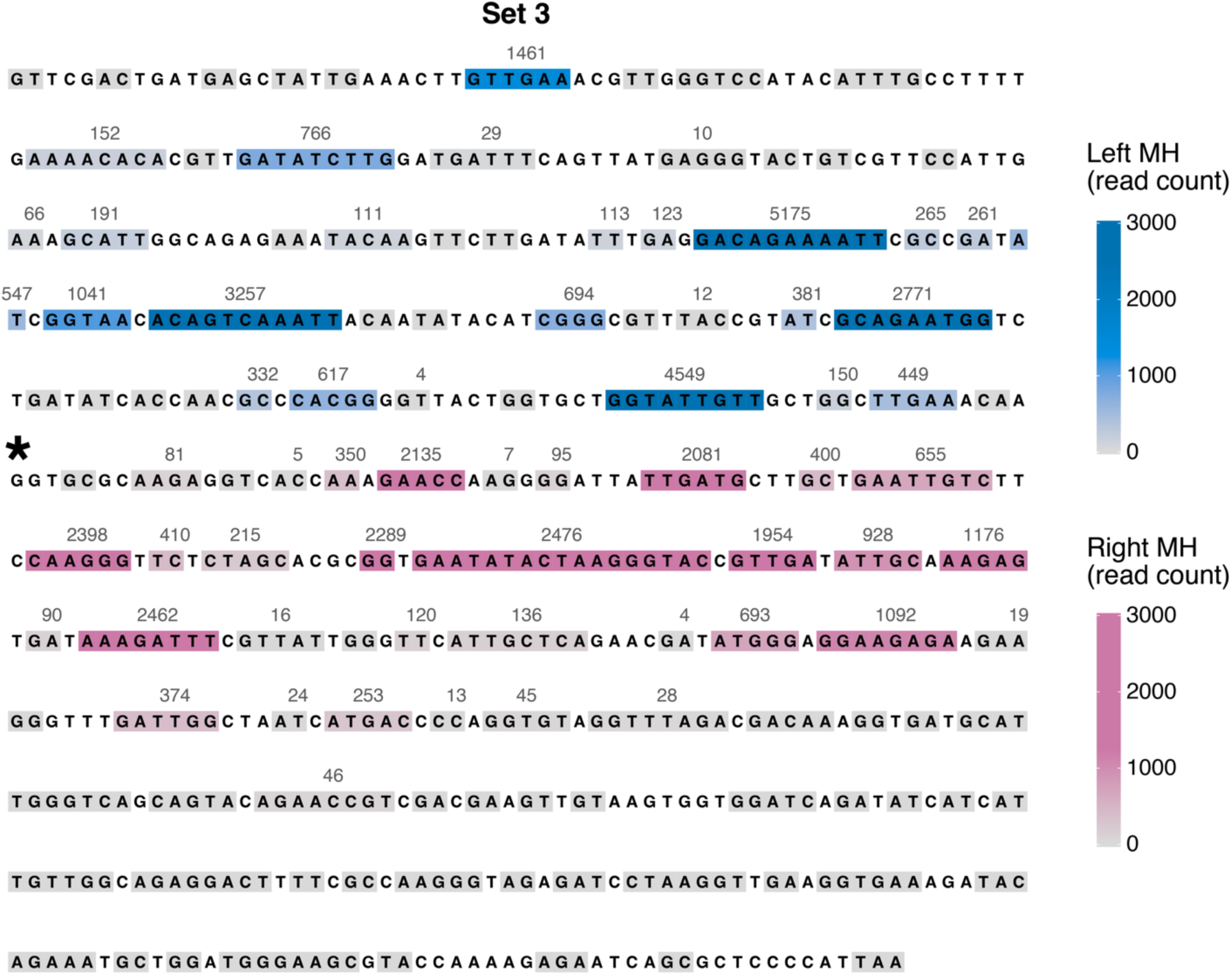
Frequency of usage of MHs between *Kl-URA3* and *Sc-ura3-52\** to form ICTS events in set 3. Grey boxes indicate unused MHs, blue boxes indicate MHs used to jump “in” to *Sc-ura3-52\** and pink boxes indicate MHs used to jump “out” of *Sc-ura3-52\**. Darker colors indicate more frequent usage. Black asterisk indicates the introduced -1 frameshift.

**Figure S4:**
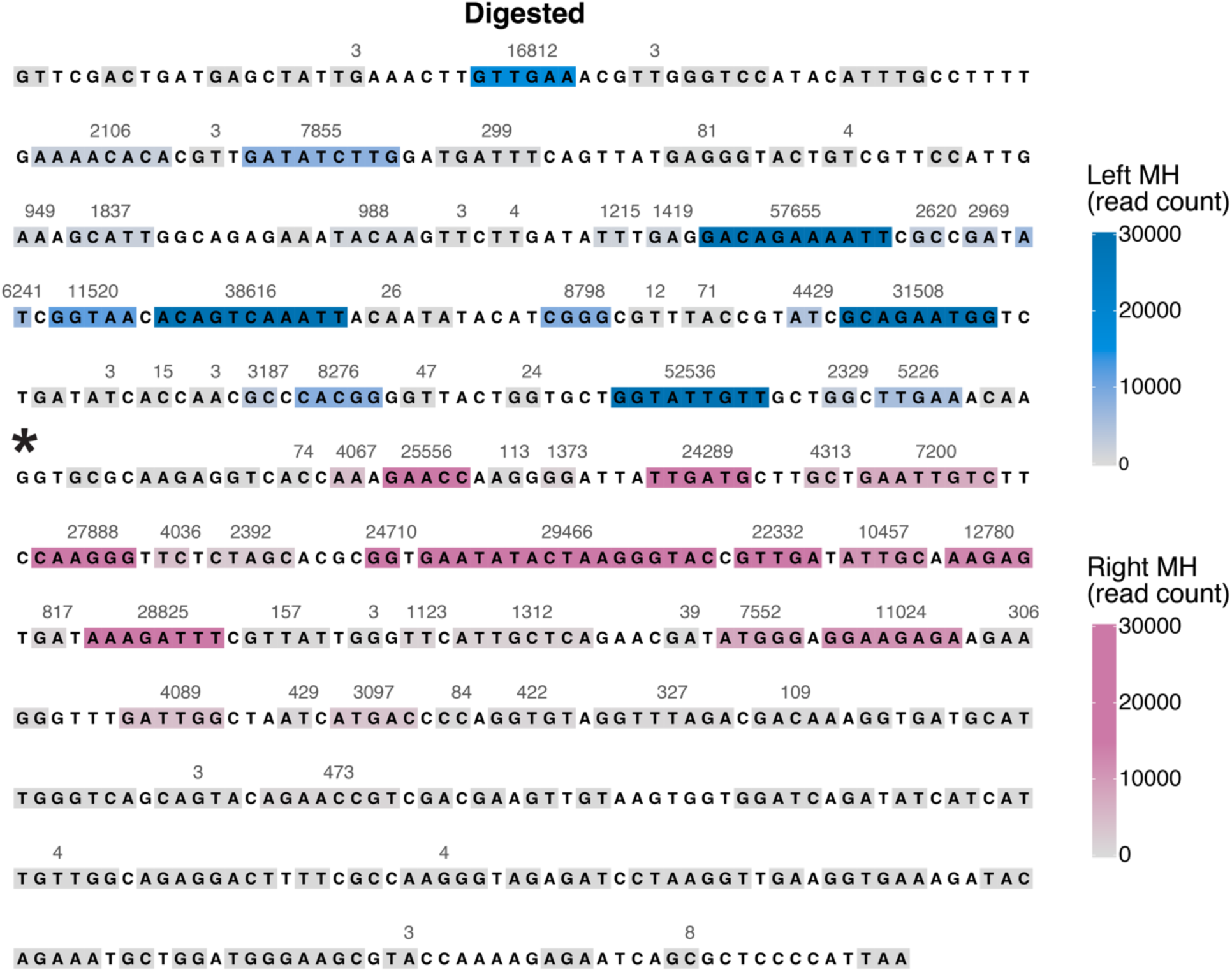
Frequency of usage of MHs between *Kl-URA3* and *Sc-ura3-52\** to form ICTS events in the *Bst*EII and *Fsp*I digested sample. Grey boxes indicate unused MHs, blue boxes indicate MHs used to jump “in” to *Sc-ura3-52\** and pink boxes indicate MHs used to jump “out” of *Sc-ura3-52\**. Darker colors indicate more frequent usage. Black asterisk indicates the introduced -1 frameshift.

**Figure S5:**
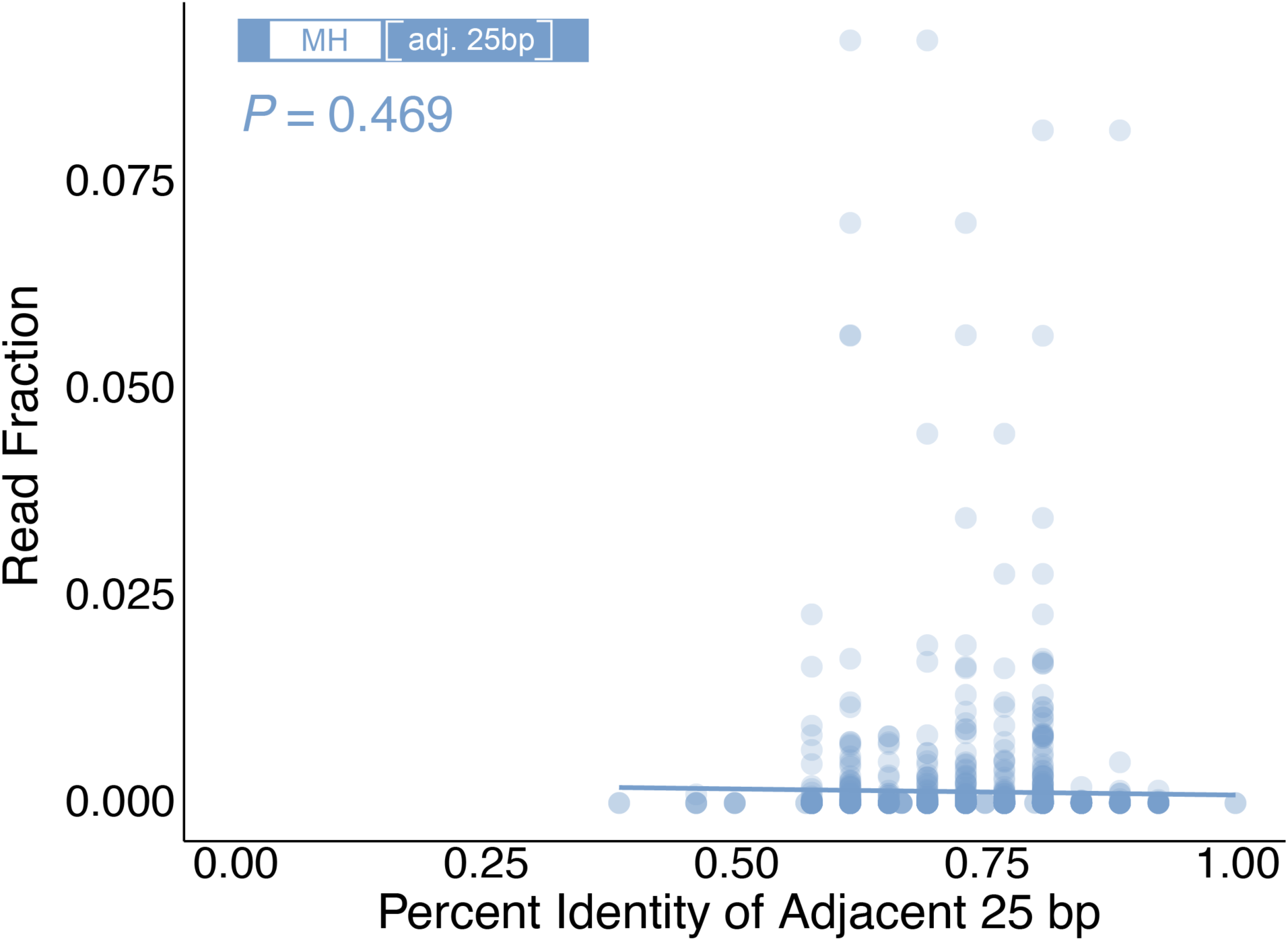
Comparison of frequency of specific MH usage and homeology in the adjacent 25 bp across ICTS events enriched via *Bst*EII and *Fsp*I digestion. A linear regression was fit to the read fractions vs. percent homeology values. A one-sample two-tailed t-test calculates the probability that the slope of the line is different from 0.

## Supplementary Table Legends

**Table S1**

Statistics of experimental details and mutations identified in each dataset described in the manuscript.

**Table S2**

Data from Liu and Zhang (8) describing spontaneous mutations acquired during growth on a variety of media types.

**Table S3**

Strains used in this study.

T**able S4**

Primers used for amplification and sequencing of DNA extracted from colonies described in the methods section.

## Additional Methods

Strains and plasmids. The strain used for the sequencing pools, yQW904, was derived from WH50 by using Cas9-mediated gene editing to delete G409 and change G411 to C so that the otherwise in-frame open reading frame contained a 1-bp deletion. Strain yQW892 contains a 32-bp deletion, *Kl-ura3-32*Δ*1*, at *HMR* that was previously described (6). Strain YCM1 contains *Kl-ura3-32*Δ*2*, an alternate 32bp deletion (ATATACATCGGGCGTTTACCGTATCGCAGAAT) located 109 bp 5’ of the deletion in *Kl-ura3-32*Δ*1*.

tNS2921 is strain yQW892 plus 100bp of a randomly generated DNA sequence, TCAAAGCCAGCTTTAGGATGTACATTATATCGTTCATGCCTAGGCGTAGCGTGGTGCACAC GTAGATTTTACAATCTACTAAGGTCCTTTGGACCGAACA, designated *100BP*, inserted between the end of the *Kl-URA3* open reading frame and *HMR Z*. This was constructed by Cas9-mediated gene editing. tNS2922 was constructed by cutting pNSU325 with *Bam*HI and transplacing *100BP*::*NATMX* into tNS2921 at the end of the *Sc-ura3* open reading frame. pNSU325 is pUC18 with *100BP*::*NatMX* inserted between 400-bp of *Sc-ura3* terminal ORF DNA and 400-bp of downstream DNA. pNSU326 is analogous to pNSU325 containing an altered *100BP* sequence, tcTaaCccTgcCttaggatgGacattataAcgCtcCtgcctTggTgtaCcgtggtgGacGcgtagaGtAtacaatctTctTag TtcctttggGccTaaca, where the upper-case letters represent the positions of mismatched nucleotides.

Construction of Libraries. Gene conversion was induced at *MAT*α in a strain carrying *HMR::Kl-URA3* and Sc-ura3-52* and Ura^-^ mutants were collected and pooled. Three libraries were prepared for sequencing. Library 1 was prepared by growing 5 cultures starting from single colonies in YP-lactate overnight and plating on 5-FOA+galactose plates. These were incubated at 30°C until Ura^-^ colonies appeared. Because the mutation rate associated with DSB repair is approximately 1000-fold higher than spontaneous mutations, each 5-FOA resistant colony that arises likely represents an independent repair-replication associated event. A total of approximately 33,000 colonies were collected. Genomic DNA was prepared and was used as a template for PCR amplifying the *MAT* locus. These PCR products were diluted and subsequently used as templates for PCR amplification using primers specific for *Kl-ura3* that were tagged with Illumina barcodes (**<u>Table S4</u>**). These were used in NGS amplicon sequencing.

Library 2 was prepared from cells plated on YPD and grown until individual colonies formed. These were replica plated to YPgal plates, incubated overnight and subsequently replica plated to 5-FOA plates. After incubation at 30°C, 1100 Ura^-^ papillae were isolated and pooled. Genomic DNA was purified and used for the PCR amplification of *Kl-ura3* using a *MAT*-specific primer to the right of *MAT Z* and a primer from *Kl-URA3* (**<u>Table S4</u>**). The DNA was submitted for PacBio sequencing.

Library 3 was started from two sets of Ura^-^ cells plated on 10 YPD plates each and grown to single colonies at a density of approximately 1000 colonies/plate. Both sets were replica plated to YPgal plates and incubated at 30°C. The colonies were replica-plated to 5-FOA plates and incubated at 30°C until individual papillae could be seen growing from the imprints. The papillae were pooled and genomic DNA was purified and used for 48 PCR reactions using primers specific to the *MAT*-distal region and the *MAT*a*1* promoter sequence upstream of *Kl-URA3*. The PCR products from each set were pooled, purified and submitted for PacBio sequencing. Each set was analyzed separately and eventually averaged together. In addition, a PCR product was digested with the enzymes *Bst*II and *Fsp*I, whose recognition sites start at nucleotide positions 408 and 417 in *Kl-URA3* and are absent in *Sc-URA3*. This pool is enriched for events that remove the restriction sites by ID or ICTS events. These 2 sets were distinguished by barcoding (**<u>Table S4</u>**).

## References

1. Haber, J. E. A Life Investigating Pathways That Repair Broken Chromosomes. Annu Rev Genet 50, 1–28 (2016).

2. Ira, G., Satory, D. & Haber, J. E. Conservative inheritance of newly synthesized DNA in double-strand break-induced gene conversion. Mol Cell Biol 26, 9424–9429 (2006).

3. Symington, L. S., Rothstein, R. & Lisby, M. Mechanisms and regulation of mitotic recombination in Saccharomyces cerevisiae. Genetics 198, 795–835 (2014).

4. Lee, C.-S. & Haber, J. E. Mating-type Gene Switching in Saccharomyces cerevisiae. Microbiol Spectr 3, MDNA3–0013–2014 (2015).

5. Hicks, W. M., Kim, M. & Haber, J. E. Increased mutagenesis and unique mutation signature associated with mitotic gene conversion. Science 329, 82–85 (2010).

6. Tsaponina, O. & Haber, J. E. Frequent Interchromosomal Template Switches during Gene Conversion in S. cerevisiae. Mol Cell 55, 615–625 (2014).

7. Sugawara, N., Towne, M. J., Lovett, S. T. & Haber, J. E. Spontaneous and double-strand break repair-associated quasipalindrome and frameshift mutagenesis in budding yeast: role of mismatch repair. Genetics 227, (2024).

8. Liu, H. & Zhang, J. The rate and molecular spectrum of mutation are selectively maintained in yeast. Nat Commun 12, 4044 (2021).

9. Hastings, P. J., Ira, G. & Lupski, J. R. A microhomology-mediated break-induced replication model for the origin of human copy number variation. PLoS Genet 5, e1000327 (2009).

10. Payen, C., Koszul, R., Dujon, B. & Fischer, G. Segmental duplications arise from Pol32-dependent repair of broken forks through two alternative replication-based mechanisms. PLoS Genet 4, e1000175 (2008).

11. Duan, Z. et al. A three-dimensional model of the yeast genome. Nature 465, 363–367 (2010).

12. Anand, R., Beach, A., Li, K. & Haber, J. Rad51-mediated double-strand break repair and mismatch correction of divergent substrates. Nature 544, 377–380 (2017).

13. Reyes, G. X. et al. Ligation of newly replicated DNA controls the timing of DNA mismatch repair. Curr Biol 31, 1268–1276.e6 (2021).

14. Hombauer, H., Srivatsan, A., Putnam, C. D. & Kolodner, R. D. Mismatch repair, but not heteroduplex rejection, is temporally coupled to DNA replication. Science 334, 1713– 1716 (2011).

15. Hicks, W. M., Yamaguchi, M. & Haber, J. E. Real-time analysis of double-strand DNA break repair by homologous recombination. Proc Natl Acad Sci U S A 108, 3108–3115 (2011).

16. Garg, P., Stith C. M., Sabouri N., Johansson E., Burgers P. M. Idling by DNA polymerase delta maintains a ligatable nick during lagging-strand DNA replication. Genes Dev. 18, 2764–73 (2004).

17. Mehta, A., Beach, A., Haber, J. E. Homology Requirements and Competition between Gene Conversion and Break-Induced Replication during Double-Strand Break Repair. Mol Cell 65, 515–526 (2017).

18. Wang, X. et al. Role of DNA replication proteins in double-strand break-induced recombination in Saccharomyces cerevisiae. Mol Cell Biol 24, 6891–6899 (2004).

19. Ira, G., Haber, J. E. Characterization of RAD51-independent break-induced replication that acts preferentially with short homologous sequences. Mol Cell Biol. 22, 6384–92 (2002).

20. Coïc, E., Martin, J., Ryu, T., Tay S. Y., Kondev, J., Haber, J. E. Dynamics of homology searching during gene conversion in Saccharomyces cerevisiae revealed by donor competition. Genetics 189, 1225–33 (2011).

21. Fishman-Lobell, J., Haber, J. E. Removal of nonhomologous DNA ends in double-strand break recombination: the role of the yeast ultraviolet repair gene RAD1. Science 16, 258, 480–4 (1992).

22. Shah, S. S., Hartono, S. R., Chédin, F. & Heyer, W.-D. Bisulfite treatment and single-molecule real-time sequencing reveal D-loop length, position, and distribution. Elife 9, (2020).

23. Smith, T. F. & Waterman, M. S. Identification of common molecular subsequences. J Mol Biol 147, 195–197 (1981).

24. Drier, Y. et al. Somatic rearrangements across cancer reveal classes of samples with distinct patterns of DNA breakage and rearrangement-induced hypermutability. Genome Res 23, 228–235 (2013).

