## Supplementary material for "Mutations and structural variants arising during double-strand break repair": Table S1

| **Statistic** | **Set 1** | **Set 2ᵃ** | **Set 3** |
| --- | --- | --- | --- |
| **Colonies collected for sequencing** | 33,000 | 1,100 | 50,000 |
| Total Reads | 993,577 | 224,844 | 319,766 |
| WT Reads | 64,342 | 8,138 | 21,125 |
| **Percentage non-WT readsᵇ** |  |  |  |
| ICTS events | 10.40% | 8.40% | 8.00% |
| ICTS events that did not copy the -1 insertion at *Sc-Ura3-52** | 0.43% | 0.30% | 0.00% |
| MH-bounded deletions | 0.92% | 0.95% | 0.55% |
| MH-bounded TDs | 0.11% | 0.09% | 0.04% |
| MH-bounded deletions:TDs | 8.12 | 10.66 | 13.22 |
| SNVs | 50.38% | 49.97% | 39.64% |
| Homonucleotide (+1) insertions | 9.46% | 7.00% | 11.65% |
| Homonucleotide (-1) deletions | 28.10% | 31.90% | 41.10% |
| Other (+1) insertions | 0.81% | 1.78% | 5.60% |
| Other (-1) deletions | 2.18% | 1.77% | 1.24% |
| 2-3bp insertions | 0.40% | 0.37% | 0.84% |
| Ty-LTR insertions | 0.07% | 0.03% | 0.00% |
| **Event Microhomology (MH)** |  |  |  |
| Mean MH length of ICTS events | 7.72 bp | 7.98 bp | 7.37 bp |
| Mean MH length of deletions | 6.04 bp | 5.93 bp | 5.39 bp |
| Mean MH length of TDs | 4.92 bp | 3.78 bp | 3.87 bp |
| **Event Length** |  |  |  |
| Mean MH-bounded deletion length | 57.2 bp | 59.7 bp | 42.6 bp |
| Mean TD length | 19.9 bp | 21.2 bp | 51.1 bp |
| Mean ICTS event length | 165 bp | 149 bp | 179 bp |
| ᵃ averaged from two technical replicates |  |  |  |
| ᵇ percentages sum to >100% due to small minority of sequences with multiple events | | |  |
