## Supplementary material for "Mutations and structural variants arising during double-strand break repair": Table S2

| Sample | Genomic Position | Ancestral Allele | Mutant Allele |
| --- | --- | --- | --- |
| Cu-1 | ref\|NC_001134\|:792564 | G | T |
| Cu-1 | ref\|NC_001145\|:224059 | G | A |
| Cu-1 | ref\|NC_001145\|:922658 | A | AATCGCTGCCGGTGTTGCCGCC |
| Cu-1 | ref\|NC_001146\|:570996 | C | T |
| Cu-1 | ref\|NC_001147\|:581592 | T | C |
| Cu-1 | ref\|NC_001147\|:787559 | C | T |
| Cu-1 | ref\|NC_001147\|:902240 | A | G |
| Cu-1 | ref\|NC_001148\|:879037 | CT | C |
| Cu-10 | ref\|NC_001136\|:601075 | TCTACTGAAGCTCCAACTGATACTA | T |
| Cu-10 | ref\|NC_001136\|:1183637 | C | T |
| Cu-10 | ref\|NC_001139\|:562003 | G | A |
| Cu-10 | ref\|NC_001143\|:521153 | C | G |
| Cu-10 | ref\|NC_001144\|:239395 | C | A |
| Cu-10 | ref\|NC_001146\|:198070 | T | C |
| Cu-10 | ref\|NC_001147\|:428054 | C | T |
| Cu-10 | ref\|NC_001147\|:1049319 | C | T |
| Cu-10 | ref\|NC_001148\|:197730 | ATATTATTATTGT | A |
| Cu-11 | ref\|NC_001134\|:317172 | T | C |
| Cu-11 | ref\|NC_001137\|:449460 | T | A |
| Cu-11 | ref\|NC_001140\|:557061 | C | T |
| Cu-11 | ref\|NC_001142\|:402004 | GCTGTAGCGGCTGGAACTGGCA | G |
| Cu-11 | ref\|NC_001144\|:5807 | C | G |
| Cu-11 | ref\|NC_001144\|:1050621 | A | C |
| Cu-12 | ref\|NC_001140\|:336612 | C | A |
| Cu-12 | ref\|NC_001145\|:112090 | A | G |
| Cu-13 | ref\|NC_001136\|:148508 | C | T |
| Cu-13 | ref\|NC_001136\|:287278 | G | C |
| Cu-13 | ref\|NC_001139\|:454459 | G | T |
| Cu-13 | ref\|NC_001142\|:171081 | G | T |
| Cu-13 | ref\|NC_001142\|:192984 | C | T |
| Cu-14 | ref\|NC_001134\|:685180 | G | C |
| Cu-14 | ref\|NC_001136\|:527515 | G | A |
| Cu-14 | ref\|NC_001136\|:527518 | T | C |
| Cu-14 | ref\|NC_001136\|:527523 | A | G |
| Cu-14 | ref\|NC_001136\|:527524 | A | G |
| Cu-14 | ref\|NC_001136\|:527530 | A | C |
| Cu-14 | ref\|NC_001138\|:181256 | C | T |
| Cu-14 | ref\|NC_001145\|:536375 | C | A |
| Cu-14 | ref\|NC_001146\|:233883 | G | A |
| Cu-14 | ref\|NC_001146\|:299392 | C | T |
| Cu-14 | ref\|NC_001148\|:510556 | T | C |
| Cu-15 | ref\|NC_001136\|:33328 | C | CAT |
| Cu-15 | ref\|NC_001138\|:4853 | C | A |
| Cu-15 | ref\|NC_001138\|:234483 | A | T |
| Cu-15 | ref\|NC_001139\|:1052640 | A | C |
| Cu-16 | ref\|NC_001134\|:353634 | A | ATG |
| Cu-16 | ref\|NC_001136\|:167020 | C | T |
| Cu-16 | ref\|NC_001136\|:353889 | C | A |
| Cu-16 | ref\|NC_001136\|:1092308 | G | T |
| Cu-16 | ref\|NC_001141\|:72235 | G | T |
| Cu-16 | ref\|NC_001147\|:182206 | G | A |
| Cu-16 | ref\|NC_001147\|:835339 | C | A |
| Cu-17 | ref\|NC_001134\|:60621 | C | G |
| Cu-17 | ref\|NC_001134\|:100577 | A | G |
| Cu-17 | ref\|NC_001136\|:414297 | A | C |
| Cu-17 | ref\|NC_001136\|:514945 | T | C |
| Cu-17 | ref\|NC_001141\|:292280 | CT | C |
| Cu-17 | ref\|NC_001142\|:653111 | G | A |
| Cu-17 | ref\|NC_001142\|:653125 | A | T |
| Cu-17 | ref\|NC_001143\|:645231 | A | T |
| Cu-17 | ref\|NC_001145\|:401138 | T | G |
| Cu-17 | ref\|NC_001148\|:240508 | A | G |
| Cu-18 | ref\|NC_001136\|:1292850 | A | T |
| Cu-18 | ref\|NC_001141\|:75210 | A | G |
| Cu-18 | ref\|NC_001142\|:77573 | G | C |
| Cu-18 | ref\|NC_001148\|:572227 | T | A |
| Cu-19 | ref\|NC_001133\|:39 | C | CCCA |
| Cu-19 | ref\|NC_001134\|:541458 | G | GTCA |
| Cu-19 | ref\|NC_001142\|:652858 | C | T |
| Cu-19 | ref\|NC_001143\|:82840 | G | T |
| Cu-19 | ref\|NC_001143\|:590986 | G | T |
| Cu-19 | ref\|NC_001148\|:485565 | C | A |
| Cu-20 | ref\|NC_001136\|:1524886 | T | A |
| Cu-20 | ref\|NC_001138\|:267538 | G | T |
| Cu-20 | ref\|NC_001139\|:920086 | T | TA |
| Cu-20 | ref\|NC_001140\|:2227 | T | C |
| Cu-20 | ref\|NC_001140\|:2233 | C | T |
| Cu-20 | ref\|NC_001140\|:54314 | G | A |
| Cu-20 | ref\|NC_001145\|:60408 | T | C |
| Cu-20 | ref\|NC_001147\|:768667 | GACGACGACGACGACGACGACA | G |
| Cu-21 | ref\|NC_001134\|:189915 | T | A |
| Cu-21 | ref\|NC_001142\|:235018 | T | C |
| Cu-21 | ref\|NC_001146\|:377515 | C | G |
| Cu-21 | ref\|NC_001147\|:664964 | C | T |
| Cu-21 | ref\|NC_001148\|:329960 | C | A |
| Cu-22 | ref\|NC_001134\|:589544 | AT | A |
| Cu-22 | ref\|NC_001135\|:23852 | T | G |
| Cu-22 | ref\|NC_001136\|:116375 | C | A |
| Cu-22 | ref\|NC_001137\|:52224 | G | C |
| Cu-22 | ref\|NC_001139\|:731538 | C | T |
| Cu-22 | ref\|NC_001144\|:800924 | C | A |
| Cu-22 | ref\|NC_001147\|:943888 | C | T |
| Cu-22 | ref\|NC_001148\|:598076 | C | A |
| Cu-22 | ref\|NC_001148\|:659910 | C | A |
| Cu-23 | ref\|NC_001136\|:1384725 | G | A |
| Cu-23 | ref\|NC_001147\|:81705 | C | A |
| Cu-23 | ref\|NC_001147\|:862270 | G | A |
| Cu-23 | ref\|NC_001148\|:820689 | T | C |
| Cu-24 | ref\|NC_001136\|:1521818 | C | CA |
| Cu-24 | ref\|NC_001137\|:520563 | A | T |
| Cu-24 | ref\|NC_001138\|:100861 | C | G |
| Cu-24 | ref\|NC_001139\|:990226 | A | C |
| Cu-24 | ref\|NC_001143\|:579981 | T | C |
| Cu-3 | ref\|NC_001133\|:30046 | G | A |
| Cu-3 | ref\|NC_001140\|:236877 | G | A |
| Cu-3 | ref\|NC_001143\|:436 | G | C |
| Cu-3 | ref\|NC_001146\|:290022 | CGAAGACATCGGTGAAGACATCGGT | C |
| Cu-4 | ref\|NC_001139\|:391500 | C | T |
| Cu-4 | ref\|NC_001139\|:459333 | A | C |
| Cu-4 | ref\|NC_001142\|:390778 | T | C |
| Cu-4 | ref\|NC_001147\|:606323 | C | A |
| Cu-5 | ref\|NC_001133\|:14680 | A | G |
| Cu-5 | ref\|NC_001136\|:831708 | A | C |
| Cu-5 | ref\|NC_001140\|:227350 | T | G |
| Cu-5 | ref\|NC_001148\|:636149 | A | G |
| Cu-6 | ref\|NC_001134\|:422944 | C | A |
| Cu-6 | ref\|NC_001137\|:517395 | C | T |
| Cu-6 | ref\|NC_001143\|:59653 | G | C |
| Cu-6 | ref\|NC_001143\|:638299 | G | GA |
| Cu-6 | ref\|NC_001147\|:15863 | G | T |
| Cu-6 | ref\|NC_001147\|:763727 | G | A |
| Cu-7 | ref\|NC_001140\|:255443 | C | T |
| Cu-7 | ref\|NC_001140\|:292170 | G | T |
| Cu-7 | ref\|NC_001144\|:1032878 | T | G |
| Cu-7 | ref\|NC_001146\|:338845 | T | G |
| Cu-7 | ref\|NC_001147\|:63072 | TCATCATCATCATCAC | T |
| Cu-8 | ref\|NC_001134\|:464130 | CTGTGCT | C |
| Cu-8 | ref\|NC_001139\|:621818 | A | T |
| Cu-8 | ref\|NC_001139\|:621820 | GA | G |
| Cu-8 | ref\|NC_001140\|:35699 | C | T |
| Cu-8 | ref\|NC_001145\|:649418 | T | C |
| Cu-8 | ref\|NC_001148\|:198004 | CA | C |
| Cu-8 | ref\|NC_001148\|:719218 | A | G |
| Cu-9 | ref\|NC_001134\|:664747 | T | G |
| Cu-9 | ref\|NC_001136\|:56985 | C | CATATATATATATATATATAAATATATAGG  T |
| Cu-9 | ref\|NC_001136\|:287464 | T | C |
| Cu-9 | ref\|NC_001138\|:254633 | C | T |
| Cu-9 | ref\|NC_001139\|:652799 | G | A |
| Cu-9 | ref\|NC_001142\|:638150 | C | G |
| Cu-9 | ref\|NC_001143\|:142376 | G | A |
| Cu-9 | ref\|NC_001143\|:142388 | T | A |
| Cu-9 | ref\|NC_001143\|:142389 | T | G |
| Cu-9 | ref\|NC_001143\|:142397 | A | G |
| Cu-9 | ref\|NC_001143\|:142409 | A | G |
| Cu-9 | ref\|NC_001144\|:372641 | T | C |
| Cu-9 | ref\|NC_001145\|:96971 | C | A |
| Li-1 | ref\|NC_001134\|:802048 | A | G |
| Li-1 | ref\|NC_001135\|:185424 | G | A |
| Li-1 | ref\|NC_001136\|:55244 | C | A |
| Li-1 | ref\|NC_001136\|:260200 | C | T |
| Li-1 | ref\|NC_001136\|:631867 | T | C |
| Li-1 | ref\|NC_001136\|:643890 | A | T |
| Li-1 | ref\|NC_001136\|:1325734 | G | T |
| Li-1 | ref\|NC_001137\|:452036 | G | T |
| Li-1 | ref\|NC_001139\|:71983 | G | T |
| Li-1 | ref\|NC_001139\|:78988 | CATATATATATTATGTATATATATAT | C |
| Li-1 | ref\|NC_001139\|:958520 | T | C |
| Li-1 | ref\|NC_001140\|:97079 | T | C |
| Li-1 | ref\|NC_001141\|:132416 | C | A |
| Li-1 | ref\|NC_001141\|:316349 | C | T |
| Li-1 | ref\|NC_001142\|:8285 | T | C |
| Li-1 | ref\|NC_001142\|:263078 | C | T |
| Li-1 | ref\|NC_001143\|:26997 | G | T |
| Li-1 | ref\|NC_001143\|:556417 | A | AAT |
| Li-1 | ref\|NC_001143\|:666237 | AG | A |
| Li-1 | ref\|NC_001143\|:666239 | T | C |
| Li-1 | ref\|NC_001144\|:133704 | A | G |
| Li-1 | ref\|NC_001145\|:748756 | G | T |
| Li-1 | ref\|NC_001146\|:374062 | G | GA |
| Li-1 | ref\|NC_001146\|:424816 | T | TA |
| Li-1 | ref\|NC_001146\|:722744 | T | C |
| Li-1 | ref\|NC_001147\|:14469 | T | C |
| Li-1 | ref\|NC_001147\|:53371 | C | A |
| Li-1 | ref\|NC_001148\|:347675 | A | AG |
| Li-10 | ref\|NC_001134\|:199834 | C | A |
| Li-10 | ref\|NC_001134\|:307654 | A | AT |
| Li-10 | ref\|NC_001136\|:675160 | C | A |
| Li-10 | ref\|NC_001136\|:1218967 | C | A |
| Li-10 | ref\|NC_001139\|:864371 | A | C |
| Li-10 | ref\|NC_001141\|:417349 | T | C |
| Li-10 | ref\|NC_001142\|:271258 | T | C |
| Li-10 | ref\|NC_001142\|:391566 | C | T |
| Li-10 | ref\|NC_001144\|:396588 | C | T |
| Li-10 | ref\|NC_001146\|:154199 | T | C |
| Li-10 | ref\|NC_001147\|:334452 | A | T |
| Li-11 | ref\|NC_001137\|:139405 | C | T |
| Li-11 | ref\|NC_001143\|:364323 | GAATATGAATATA | G |
| Li-11 | ref\|NC_001144\|:85392 | G | A |
| Li-11 | ref\|NC_001145\|:414507 | A | G |
| Li-11 | ref\|NC_001146\|:167479 | A | G |
| Li-11 | ref\|NC_001148\|:599733 | A | G |
| Li-12 | ref\|NC_001133\|:152624 | C | T |
| Li-12 | ref\|NC_001136\|:154666 | T | C |
| Li-12 | ref\|NC_001138\|:123211 | CT | C |
| Li-12 | ref\|NC_001139\|:51130 | A | G |
| Li-12 | ref\|NC_001139\|:549197 | A | G |
| Li-12 | ref\|NC_001140\|:520253 | A | G |
| Li-12 | ref\|NC_001141\|:40588 | G | T |
| Li-12 | ref\|NC_001142\|:74293 | T | C |
| Li-12 | ref\|NC_001145\|:290831 | G | C |
| Li-12 | ref\|NC_001148\|:11889 | G | A |
| Li-12 | ref\|NC_001148\|:541565 | A | G |
| Li-13 | ref\|NC_001134\|:554380 | T | C |
| Li-13 | ref\|NC_001135\|:300779 | G | A |
| Li-13 | ref\|NC_001136\|:913300 | G | A |
| Li-13 | ref\|NC_001136\|:1037221 | T | C |
| Li-13 | ref\|NC_001136\|:1213343 | T | G |
| Li-13 | ref\|NC_001138\|:179122 | G | A |
| Li-13 | ref\|NC_001139\|:533083 | C | T |
| Li-13 | ref\|NC_001139\|:695349 | T | C |
| Li-13 | ref\|NC_001140\|:197 | C | T |
| Li-13 | ref\|NC_001141\|:267671 | GGGCATCCAGCTCCGGTAGGAGCT  GGATTGCTCTTCTTC | G |
| Li-13 | ref\|NC_001142\|:350005 | C | G |
| Li-13 | ref\|NC_001143\|:149727 | A | G |
| Li-13 | ref\|NC_001147\|:830554 | C | T |
| Li-13 | ref\|NC_001147\|:896828 | TTTCTTCTCTTTCTTCTCTTTCTTGG  ACTTCTTTTCC | T |
| Li-13 | ref\|NC_001148\|:280980 | A | G |
| Li-13 | ref\|NC_001148\|:829370 | A | G |
| Li-14 | ref\|NC_001134\|:50433 | C | A |
| Li-14 | ref\|NC_001134\|:588002 | T | A |
| Li-14 | ref\|NC_001134\|:658496 | C | A |
| Li-14 | ref\|NC_001135\|:82820 | T | G |
| Li-14 | ref\|NC_001136\|:408905 | G | A |
| Li-14 | ref\|NC_001136\|:443942 | G | GA |
| Li-14 | ref\|NC_001136\|:732973 | C | T |
| Li-14 | ref\|NC_001138\|:125127 | A | G |
| Li-14 | ref\|NC_001139\|:988894 | C | CCAG |
| Li-14 | ref\|NC_001139\|:1020193 | G | A |
| Li-14 | ref\|NC_001140\|:366962 | C | A |
| Li-14 | ref\|NC_001142\|:500547 | C | A |
| Li-14 | ref\|NC_001143\|:644007 | A | T |
| Li-14 | ref\|NC_001144\|:317770 | T | C |
| Li-14 | ref\|NC_001144\|:667275 | C | T |
| Li-14 | ref\|NC_001144\|:784444 | G | T |
| Li-14 | ref\|NC_001144\|:916488 | T | G |
| Li-14 | ref\|NC_001145\|:906074 | G | A |
| Li-14 | ref\|NC_001146\|:108957 | T | C |
| Li-14 | ref\|NC_001147\|:636324 | G | T |
| Li-14 | ref\|NC_001147\|:848167 | T | G |
| Li-15 | ref\|NC_001134\|:778042 | G | T |
| Li-15 | ref\|NC_001136\|:40379 | G | C |
| Li-15 | ref\|NC_001136\|:206222 | T | A |
| Li-15 | ref\|NC_001136\|:332972 | G | GA |
| Li-15 | ref\|NC_001136\|:717707 | C | A |
| Li-15 | ref\|NC_001136\|:1372154 | C | T |
| Li-15 | ref\|NC_001137\|:183606 | C | T |
| Li-15 | ref\|NC_001138\|:6428 | G | T |
| Li-15 | ref\|NC_001139\|:138890 | G | T |
| Li-15 | ref\|NC_001139\|:567717 | T | C |
| Li-15 | ref\|NC_001139\|:991287 | C | A |
| Li-15 | ref\|NC_001139\|:1037328 | G | GA |
| Li-15 | ref\|NC_001140\|:97829 | C | A |
| Li-15 | ref\|NC_001140\|:378120 | GTGTTTT | G |
| Li-15 | ref\|NC_001141\|:149542 | G | T |
| Li-15 | ref\|NC_001142\|:167492 | C | G |
| Li-15 | ref\|NC_001142\|:393708 | C | CT |
| Li-15 | ref\|NC_001144\|:698633 | C | T |
| Li-15 | ref\|NC_001144\|:1019875 | C | T |
| Li-15 | ref\|NC_001145\|:760792 | G | T |
| Li-15 | ref\|NC_001146\|:325286 | A | G |
| Li-15 | ref\|NC_001146\|:720276 | CTGTTGCTGCTGTTGT | C |
| Li-15 | ref\|NC_001147\|:227300 | C | CGAT |
| Li-15 | ref\|NC_001147\|:768824 | C | A |
| Li-15 | ref\|NC_001147\|:1058301 | T | C |
| Li-15 | ref\|NC_001148\|:242623 | C | T |
| Li-15 | ref\|NC_001148\|:920298 | G | A |
| Li-16 | ref\|NC_001134\|:694353 | T | C |
| Li-16 | ref\|NC_001135\|:4577 | A | G |
| Li-16 | ref\|NC_001136\|:105881 | A | T |
| Li-16 | ref\|NC_001136\|:207677 | C | T |
| Li-16 | ref\|NC_001136\|:215204 | G | A |
| Li-16 | ref\|NC_001136\|:383937 | A | G |
| Li-16 | ref\|NC_001136\|:1320113 | C | A |
| Li-16 | ref\|NC_001136\|:1379400 | G | A |
| Li-16 | ref\|NC_001136\|:1389525 | G | GA |
| Li-16 | ref\|NC_001137\|:357337 | G | A |
| Li-16 | ref\|NC_001139\|:278147 | T | G |
| Li-16 | ref\|NC_001139\|:832444 | C | CAT |
| Li-16 | ref\|NC_001139\|:1014908 | A | G |
| Li-16 | ref\|NC_001141\|:386182 | A | G |
| Li-16 | ref\|NC_001141\|:432657 | T | G |
| Li-16 | ref\|NC_001142\|:366490 | G | C |
| Li-16 | ref\|NC_001143\|:357984 | TA | T |
| Li-16 | ref\|NC_001144\|:393645 | G | A |
| Li-16 | ref\|NC_001144\|:736702 | C | A |
| Li-16 | ref\|NC_001144\|:923672 | G | T |
| Li-16 | ref\|NC_001145\|:103182 | T | C |
| Li-16 | ref\|NC_001145\|:158567 | CCACAGCA | C |
| Li-16 | ref\|NC_001145\|:158575 | A | T |
| Li-16 | ref\|NC_001145\|:158577 | G | GTTGCTGT |
| Li-16 | ref\|NC_001145\|:158578 | A | G |
| Li-16 | ref\|NC_001145\|:389559 | G | A |
| Li-16 | ref\|NC_001146\|:56333 | A | T |
| Li-16 | ref\|NC_001146\|:709447 | G | T |
| Li-16 | ref\|NC_001147\|:136354 | GAAAAATTTTTCCCATATATTTG  AAAA | G |
| Li-16 | ref\|NC_001147\|:824412 | G | A |
| Li-17 | ref\|NC_001136\|:992864 | G | T |
| Li-17 | ref\|NC_001136\|:1246935 | C | A |
| Li-17 | ref\|NC_001137\|:378373 | T | C |
| Li-17 | ref\|NC_001138\|:30604 | A | G |
| Li-17 | ref\|NC_001139\|:832639 | G | A |
| Li-17 | ref\|NC_001139\|:1017221 | A | C |
| Li-17 | ref\|NC_001141\|:20351 | T | C |
| Li-17 | ref\|NC_001141\|:28342 | C | T |
| Li-17 | ref\|NC_001142\|:139760 | A | T |
| Li-17 | ref\|NC_001142\|:493501 | C | A |
| Li-17 | ref\|NC_001143\|:162195 | G | A |
| Li-17 | ref\|NC_001143\|:434875 | C | T |
| Li-17 | ref\|NC_001144\|:342885 | A | G |
| Li-17 | ref\|NC_001145\|:260696 | A | T |
| Li-17 | ref\|NC_001146\|:150930 | T | G |
| Li-17 | ref\|NC_001146\|:661755 | A | AAT |
| Li-17 | ref\|NC_001146\|:708532 | C | A |
| Li-17 | ref\|NC_001147\|:535487 | G | GA |
| Li-17 | ref\|NC_001148\|:835364 | T | A |
| Li-18 | ref\|NC_001134\|:696778 | G | A |
| Li-18 | ref\|NC_001134\|:721265 | G | GA |
| Li-18 | ref\|NC_001135\|:9758 | T | A |
| Li-18 | ref\|NC_001136\|:155732 | T | TA |
| Li-18 | ref\|NC_001136\|:220080 | G | T |
| Li-18 | ref\|NC_001136\|:374731 | C | T |
| Li-18 | ref\|NC_001136\|:818976 | T | A |
| Li-18 | ref\|NC_001136\|:942410 | A | AT |
| Li-18 | ref\|NC_001136\|:1315128 | A | G |
| Li-18 | ref\|NC_001139\|:113786 | A | G |
| Li-18 | ref\|NC_001139\|:286663 | C | A |
| Li-18 | ref\|NC_001139\|:298593 | G | T |
| Li-18 | ref\|NC_001139\|:744885 | T | A |
| Li-18 | ref\|NC_001139\|:990811 | A | G |
| Li-18 | ref\|NC_001139\|:1024090 | G | A |
| Li-18 | ref\|NC_001140\|:316327 | A | T |
| Li-18 | ref\|NC_001140\|:528484 | C | A |
| Li-18 | ref\|NC_001141\|:165202 | A | T |
| Li-18 | ref\|NC_001142\|:592450 | GT | G |
| Li-18 | ref\|NC_001143\|:118299 | T | A |
| Li-18 | ref\|NC_001143\|:632965 | T | C |
| Li-18 | ref\|NC_001144\|:128859 | G | A |
| Li-18 | ref\|NC_001144\|:243453 | A | AT |
| Li-18 | ref\|NC_001145\|:96680 | C | G |
| Li-18 | ref\|NC_001145\|:372437 | TA | T |
| Li-18 | ref\|NC_001145\|:405864 | A | T |
| Li-18 | ref\|NC_001145\|:431003 | T | TA |
| Li-18 | ref\|NC_001146\|:459271 | T | C |
| Li-18 | ref\|NC_001146\|:616706 | C | A |
| Li-18 | ref\|NC_001147\|:78640 | C | T |
| Li-18 | ref\|NC_001147\|:221897 | C | T |
| Li-18 | ref\|NC_001147\|:714497 | T | TA |
| Li-18 | ref\|NC_001147\|:884768 | C | T |
| Li-18 | ref\|NC_001147\|:925870 | A | C |
| Li-19 | ref\|NC_001136\|:401714 | G | T |
| Li-19 | ref\|NC_001136\|:1008355 | C | A |
| Li-19 | ref\|NC_001137\|:79234 | A | T |
| Li-19 | ref\|NC_001137\|:538998 | C | T |
| Li-19 | ref\|NC_001139\|:526371 | A | G |
| Li-19 | ref\|NC_001141\|:438521 | G | A |
| Li-19 | ref\|NC_001142\|:703431 | C | A |
| Li-19 | ref\|NC_001145\|:166044 | G | T |
| Li-19 | ref\|NC_001145\|:811636 | GTTTGGACTGGGGCTGAAT | G |
| Li-19 | ref\|NC_001146\|:660702 | C | A |
| Li-19 | ref\|NC_001147\|:991665 | G | A |
| Li-2 | ref\|NC_001135\|:313479 | T | TA |
| Li-2 | ref\|NC_001136\|:149113 | A | AAAAT |
| Li-2 | ref\|NC_001136\|:360004 | T | A |
| Li-2 | ref\|NC_001136\|:850650 | T | C |
| Li-2 | ref\|NC_001136\|:1241094 | T | TA |
| Li-2 | ref\|NC_001137\|:35429 | A | G |
| Li-2 | ref\|NC_001139\|:87586 | C | CT |
| Li-2 | ref\|NC_001139\|:191611 | C | T |
| Li-2 | ref\|NC_001139\|:775328 | G | A |
| Li-2 | ref\|NC_001140\|:332165 | G | T |
| Li-2 | ref\|NC_001141\|:251677 | G | GA |
| Li-2 | ref\|NC_001142\|:117487 | C | A |
| Li-2 | ref\|NC_001142\|:219946 | T | C |
| Li-2 | ref\|NC_001142\|:737876 | C | A |
| Li-2 | ref\|NC_001143\|:240481 | G | A |
| Li-2 | ref\|NC_001143\|:653468 | A | T |
| Li-2 | ref\|NC_001144\|:232706 | C | A |
| Li-2 | ref\|NC_001144\|:932258 | A | AGATGTACG |
| Li-2 | ref\|NC_001144\|:1057912 | T | A |
| Li-2 | ref\|NC_001147\|:362723 | T | C |
| Li-2 | ref\|NC_001147\|:579826 | C | A |
| Li-2 | ref\|NC_001147\|:600687 | A | G |
| Li-2 | ref\|NC_001147\|:600689 | C | G |
| Li-2 | ref\|NC_001147\|:600690 | A | C |
| Li-2 | ref\|NC_001148\|:449277 | A | G |
| Li-2 | ref\|NC_001148\|:620276 | A | G |
| Li-20 | ref\|NC_001136\|:773766 | A | G |
| Li-20 | ref\|NC_001136\|:1074887 | A | C |
| Li-20 | ref\|NC_001136\|:1308549 | G | A |
| Li-20 | ref\|NC_001139\|:886496 | A | G |
| Li-20 | ref\|NC_001139\|:1059027 | A | G |
| Li-20 | ref\|NC_001140\|:300040 | T | C |
| Li-20 | ref\|NC_001143\|:182668 | G | GT |
| Li-20 | ref\|NC_001144\|:149790 | G | A |
| Li-20 | ref\|NC_001146\|:56333 | A | T |
| Li-20 | ref\|NC_001148\|:219677 | G | T |
| Li-20 | ref\|NC_001148\|:829704 | T | G |
| Li-21 | ref\|NC_001134\|:526831 | G | GA |
| Li-21 | ref\|NC_001135\|:191617 | A | G |
| Li-21 | ref\|NC_001136\|:435770 | C | A |
| Li-21 | ref\|NC_001136\|:707062 | G | A |
| Li-21 | ref\|NC_001136\|:1498784 | T | C |
| Li-21 | ref\|NC_001139\|:1039367 | G | GA |
| Li-21 | ref\|NC_001142\|:570788 | T | A |
| Li-21 | ref\|NC_001142\|:620606 | T | C |
| Li-21 | ref\|NC_001143\|:167518 | A | AAT |
| Li-21 | ref\|NC_001143\|:169000 | A | G |
| Li-21 | ref\|NC_001144\|:31638 | C | A |
| Li-21 | ref\|NC_001144\|:320784 | T | C |
| Li-21 | ref\|NC_001144\|:675761 | T | C |
| Li-21 | ref\|NC_001145\|:811237 | G | GGTT |
| Li-21 | ref\|NC_001146\|:69054 | A | G |
| Li-21 | ref\|NC_001146\|:94900 | A | T |
| Li-21 | ref\|NC_001146\|:417802 | TC | T |
| Li-21 | ref\|NC_001146\|:778892 | T | C |
| Li-21 | ref\|NC_001147\|:40189 | A | G |
| Li-21 | ref\|NC_001147\|:1032474 | G | T |
| Li-21 | ref\|NC_001148\|:114361 | C | T |
| Li-21 | ref\|NC_001148\|:188912 | T | C |
| Li-21 | ref\|NC_001148\|:353870 | C | T |
| Li-21 | ref\|NC_001148\|:481366 | G | GA |
| Li-22 | ref\|NC_001134\|:471073 | A | G |
| Li-22 | ref\|NC_001135\|:83153 | C | G |
| Li-22 | ref\|NC_001136\|:194843 | T | C |
| Li-22 | ref\|NC_001136\|:285692 | A | G |
| Li-22 | ref\|NC_001136\|:1362773 | A | AT |
| Li-22 | ref\|NC_001137\|:91544 | C | A |
| Li-22 | ref\|NC_001137\|:513034 | T | C |
| Li-22 | ref\|NC_001139\|:96645 | C | T |
| Li-22 | ref\|NC_001139\|:353056 | G | A |
| Li-22 | ref\|NC_001139\|:396376 | C | T |
| Li-22 | ref\|NC_001142\|:108832 | T | C |
| Li-22 | ref\|NC_001142\|:420815 | T | C |
| Li-22 | ref\|NC_001143\|:144647 | A | ACTGCTGCTGTTTCTAAGAAATCCACCG |
| Li-22 | ref\|NC_001144\|:88851 | C | A |
| Li-22 | ref\|NC_001144\|:115304 | A | G |
| Li-22 | ref\|NC_001145\|:294448 | C | T |
| Li-22 | ref\|NC_001145\|:483541 | A | C |
| Li-22 | ref\|NC_001146\|:476455 | G | T |
| Li-22 | ref\|NC_001147\|:227289 | A | AATGATGATGACGATGATGATGATG |
| Li-22 | ref\|NC_001147\|:675097 | A | G |
| Li-22 | ref\|NC_001147\|:891184 | C | T |
| Li-22 | ref\|NC_001148\|:664678 | TATAAAAGAAA | T |
| Li-23 | ref\|NC_001134\|:136718 | C | A |
| Li-23 | ref\|NC_001134\|:391501 | G | A |
| Li-23 | ref\|NC_001143\|:17007 | G | A |
| Li-23 | ref\|NC_001143\|:113257 | T | C |
| Li-23 | ref\|NC_001143\|:145645 | T | A |
| Li-23 | ref\|NC_001144\|:511640 | A | G |
| Li-23 | ref\|NC_001144\|:585845 | C | A |
| Li-23 | ref\|NC_001146\|:49643 | A | T |
| Li-23 | ref\|NC_001147\|:740252 | T | C |
| Li-23 | ref\|NC_001147\|:811288 | C | T |
| Li-23 | ref\|NC_001148\|:715059 | A | G |
| Li-23 | ref\|NC_001148\|:754378 | T | TG |
| Li-24 | ref\|NC_001134\|:94837 | T | C |
| Li-24 | ref\|NC_001134\|:424994 | T | C |
| Li-24 | ref\|NC_001134\|:743401 | C | T |
| Li-24 | ref\|NC_001135\|:208053 | A | G |
| Li-24 | ref\|NC_001136\|:134120 | G | A |
| Li-24 | ref\|NC_001136\|:278306 | T | G |
| Li-24 | ref\|NC_001136\|:306326 | G | T |
| Li-24 | ref\|NC_001136\|:605315 | A | G |
| Li-24 | ref\|NC_001136\|:683113 | C | G |
| Li-24 | ref\|NC_001136\|:823703 | C | A |
| Li-24 | ref\|NC_001136\|:900276 | G | T |
| Li-24 | ref\|NC_001136\|:1352378 | T | TA |
| Li-24 | ref\|NC_001136\|:1378481 | A | G |
| Li-24 | ref\|NC_001137\|:555985 | G | A |
| Li-24 | ref\|NC_001138\|:32114 | T | A |
| Li-24 | ref\|NC_001138\|:165630 | A | G |
| Li-24 | ref\|NC_001139\|:112795 | T | C |
| Li-24 | ref\|NC_001139\|:458106 | T | C |
| Li-24 | ref\|NC_001140\|:443891 | T | G |
| Li-24 | ref\|NC_001141\|:172917 | T | C |
| Li-24 | ref\|NC_001141\|:439846 | T | C |
| Li-24 | ref\|NC_001142\|:27064 | G | A |
| Li-24 | ref\|NC_001143\|:26340 | G | T |
| Li-24 | ref\|NC_001143\|:281244 | C | A |
| Li-24 | ref\|NC_001143\|:422678 | T | G |
| Li-24 | ref\|NC_001144\|:209471 | A | T |
| Li-24 | ref\|NC_001144\|:1037799 | A | G |
| Li-24 | ref\|NC_001145\|:11475 | T | G |
| Li-24 | ref\|NC_001145\|:402563 | G | T |
| Li-24 | ref\|NC_001145\|:420858 | C | T |
| Li-24 | ref\|NC_001145\|:445790 | T | C |
| Li-24 | ref\|NC_001145\|:456590 | A | G |
| Li-24 | ref\|NC_001145\|:874864 | A | G |
| Li-24 | ref\|NC_001146\|:186163 | G | T |
| Li-24 | ref\|NC_001146\|:418915 | G | GTA |
| Li-24 | ref\|NC_001147\|:75717 | G | A |
| Li-24 | ref\|NC_001147\|:203362 | A | T |
| Li-24 | ref\|NC_001147\|:783091 | C | T |
| Li-24 | ref\|NC_001148\|:430325 | C | A |
| Li-24 | ref\|NC_001148\|:843356 | A | T |
| Li-3 | ref\|NC_001134\|:446010 | G | A |
| Li-3 | ref\|NC_001136\|:1384907 | C | T |
| Li-3 | ref\|NC_001136\|:1392610 | C | A |
| Li-3 | ref\|NC_001136\|:1448455 | C | T |
| Li-3 | ref\|NC_001136\|:1479266 | C | T |
| Li-3 | ref\|NC_001138\|:94191 | G | C |
| Li-3 | ref\|NC_001139\|:44728 | G | T |
| Li-3 | ref\|NC_001139\|:110451 | G | C |
| Li-3 | ref\|NC_001139\|:161861 | G | A |
| Li-3 | ref\|NC_001139\|:889933 | C | A |
| Li-3 | ref\|NC_001140\|:251625 | C | A |
| Li-3 | ref\|NC_001141\|:112290 | G | A |
| Li-3 | ref\|NC_001141\|:152655 | G | A |
| Li-3 | ref\|NC_001141\|:200816 | T | A |
| Li-3 | ref\|NC_001142\|:42711 | C | T |
| Li-3 | ref\|NC_001143\|:331877 | A | T |
| Li-3 | ref\|NC_001144\|:824697 | G | A |
| Li-3 | ref\|NC_001145\|:694163 | T | TA |
| Li-3 | ref\|NC_001145\|:891007 | C | T |
| Li-3 | ref\|NC_001146\|:59156 | C | T |
| Li-3 | ref\|NC_001146\|:68162 | A | G |
| Li-3 | ref\|NC_001147\|:209654 | C | A |
| Li-3 | ref\|NC_001147\|:682694 | G | A |
| Li-3 | ref\|NC_001147\|:756280 | T | C |
| Li-4 | ref\|NC_001134\|:175476 | C | T |
| Li-4 | ref\|NC_001134\|:808065 | C | A |
| Li-4 | ref\|NC_001136\|:94640 | T | G |
| Li-4 | ref\|NC_001136\|:781521 | A | T |
| Li-4 | ref\|NC_001136\|:864020 | A | G |
| Li-4 | ref\|NC_001136\|:1290576 | T | A |
| Li-4 | ref\|NC_001141\|:324294 | TA | T |
| Li-4 | ref\|NC_001142\|:418340 | G | C |
| Li-4 | ref\|NC_001143\|:120751 | T | G |
| Li-4 | ref\|NC_001143\|:370697 | G | A |
| Li-4 | ref\|NC_001143\|:616704 | T | C |
| Li-4 | ref\|NC_001144\|:63675 | C | T |
| Li-4 | ref\|NC_001145\|:340488 | T | C |
| Li-4 | ref\|NC_001145\|:463162 | C | A |
| Li-4 | ref\|NC_001145\|:580165 | G | A |
| Li-4 | ref\|NC_001146\|:632588 | G | T |
| Li-4 | ref\|NC_001147\|:634216 | C | A |
| Li-4 | ref\|NC_001148\|:402757 | C | A |
| Li-4 | ref\|NC_001148\|:639909 | G | A |
| Li-4 | ref\|NC_001148\|:684436 | A | AT |
| Li-4 | ref\|NC_001148\|:939365 | A | T |
| Li-5 | ref\|NC_001134\|:758907 | C | T |
| Li-5 | ref\|NC_001135\|:304964 | G | A |
| Li-5 | ref\|NC_001136\|:461179 | C | T |
| Li-5 | ref\|NC_001136\|:592363 | G | A |
| Li-5 | ref\|NC_001136\|:1381423 | C | T |
| Li-5 | ref\|NC_001137\|:130736 | A | AT |
| Li-5 | ref\|NC_001137\|:449492 | G | A |
| Li-5 | ref\|NC_001139\|:1084303 | G | T |
| Li-5 | ref\|NC_001140\|:10418 | A | T |
| Li-5 | ref\|NC_001140\|:210848 | A | C |
| Li-5 | ref\|NC_001141\|:147631 | T | C |
| Li-5 | ref\|NC_001141\|:355926 | T | TA |
| Li-5 | ref\|NC_001142\|:34976 | G | A |
| Li-5 | ref\|NC_001142\|:34977 | G | T |
| Li-5 | ref\|NC_001142\|:34980 | C | G |
| Li-5 | ref\|NC_001142\|:291655 | G | T |
| Li-5 | ref\|NC_001142\|:541478 | CAGAAAATCCAGATTTGTACAGAA | C |
| Li-5 | ref\|NC_001143\|:77348 | A | G |
| Li-5 | ref\|NC_001143\|:209032 | A | G |
| Li-5 | ref\|NC_001144\|:405688 | G | A |
| Li-5 | ref\|NC_001145\|:330117 | T | G |
| Li-5 | ref\|NC_001146\|:308452 | C | A |
| Li-5 | ref\|NC_001147\|:526158 | G | A |
| Li-5 | ref\|NC_001147\|:855699 | TCCACGTCCTCGCCCTCGA | T |
| Li-6 | ref\|NC_001134\|:581558 | A | AT |
| Li-6 | ref\|NC_001136\|:430897 | A | G |
| Li-6 | ref\|NC_001136\|:595656 | C | A |
| Li-6 | ref\|NC_001136\|:1080944 | GTGTATGTAATGTTAAGCGCTAAATA | G |
| Li-6 | ref\|NC_001136\|:1332435 | A | G |
| Li-6 | ref\|NC_001137\|:382541 | A | T |
| Li-6 | ref\|NC_001137\|:421843 | C | CT |
| Li-6 | ref\|NC_001138\|:6856 | T | G |
| Li-6 | ref\|NC_001138\|:39970 | T | A |
| Li-6 | ref\|NC_001139\|:166464 | T | C |
| Li-6 | ref\|NC_001140\|:463379 | T | C |
| Li-6 | ref\|NC_001140\|:550134 | T | G |
| Li-6 | ref\|NC_001143\|:631098 | T | TA |
| Li-6 | ref\|NC_001144\|:1064339 | A | G |
| Li-6 | ref\|NC_001144\|:1064375 | T | A |
| Li-6 | ref\|NC_001144\|:1064396 | T | C |
| Li-6 | ref\|NC_001145\|:685893 | A | T |
| Li-6 | ref\|NC_001146\|:45766 | G | GT |
| Li-6 | ref\|NC_001148\|:30494 | C | T |
| Li-7 | ref\|NC_001134\|:777106 | G | A |
| Li-7 | ref\|NC_001136\|:271119 | G | A |
| Li-7 | ref\|NC_001136\|:514621 | G | T |
| Li-7 | ref\|NC_001136\|:851614 | G | C |
| Li-7 | ref\|NC_001136\|:1226057 | C | A |
| Li-7 | ref\|NC_001137\|:389638 | G | A |
| Li-7 | ref\|NC_001137\|:421909 | GATCT | G |
| Li-7 | ref\|NC_001137\|:433468 | C | G |
| Li-7 | ref\|NC_001138\|:101479 | CT | C |
| Li-7 | ref\|NC_001139\|:245641 | TCAACAACAACAACAACAA | T |
| Li-7 | ref\|NC_001139\|:909466 | A | T |
| Li-7 | ref\|NC_001142\|:253113 | C | G |
| Li-7 | ref\|NC_001142\|:351648 | T | C |
| Li-7 | ref\|NC_001142\|:506570 | G | A |
| Li-7 | ref\|NC_001143\|:162494 | G | T |
| Li-7 | ref\|NC_001143\|:634952 | C | A |
| Li-7 | ref\|NC_001145\|:682812 | G | A |
| Li-7 | ref\|NC_001146\|:129414 | T | TTA |
| Li-7 | ref\|NC_001146\|:132660 | C | A |
| Li-7 | ref\|NC_001146\|:224755 | C | T |
| Li-7 | ref\|NC_001147\|:56397 | A | T |
| Li-7 | ref\|NC_001147\|:398534 | A | G |
| Li-7 | ref\|NC_001147\|:963994 | G | A |
| Li-7 | ref\|NC_001148\|:371470 | C | G |
| Li-9 | ref\|NC_001133\|:194369 | A | T |
| Li-9 | ref\|NC_001135\|:62898 | C | A |
| Li-9 | ref\|NC_001135\|:303527 | A | G |
| Li-9 | ref\|NC_001136\|:633969 | C | A |
| Li-9 | ref\|NC_001136\|:1513804 | G | A |
| Li-9 | ref\|NC_001138\|:190621 | T | TG |
| Li-9 | ref\|NC_001139\|:824279 | A | C |
| Li-9 | ref\|NC_001140\|:378120 | G | GTGTTTTTGTTTTTGTTTTTGTTTT |
| Li-9 | ref\|NC_001140\|:528882 | A | AT |
| Li-9 | ref\|NC_001142\|:342037 | A | G |
| Li-9 | ref\|NC_001142\|:637260 | C | A |
| Li-9 | ref\|NC_001144\|:823442 | GGAGAGAGAGAGAGAGAGAGAG  AGAGAGAGAGA | G |
| Li-9 | ref\|NC_001146\|:291633 | T | C |
| Li-9 | ref\|NC_001146\|:365595 | T | C |
| Li-9 | ref\|NC_001146\|:427977 | T | C |
| Li-9 | ref\|NC_001147\|:258421 | G | T |
| Li-9 | ref\|NC_001147\|:354155 | A | C |
| Li-9 | ref\|NC_001147\|:545172 | G | A |
| Na-1 | ref\|NC_001133\|:43067 | G | T |
| Na-1 | ref\|NC_001136\|:592539 | ATGCAAATGCAACTGCAAATGCAAC | A |
| Na-1 | ref\|NC_001136\|:1349609 | T | TA |
| Na-1 | ref\|NC_001139\|:287219 | G | GA |
| Na-1 | ref\|NC_001139\|:637574 | A | G |
| Na-1 | ref\|NC_001142\|:54126 | C | A |
| Na-1 | ref\|NC_001144\|:194782 | G | T |
| Na-1 | ref\|NC_001144\|:730759 | C | A |
| Na-1 | ref\|NC_001144\|:1065041 | T | G |
| Na-1 | ref\|NC_001145\|:266490 | A | C |
| Na-1 | ref\|NC_001147\|:63078 | TCATCATCAC | T |
| Na-1 | ref\|NC_001148\|:242379 | C | A |
| Na-1 | ref\|NC_001148\|:876066 | T | G |
| Na-10 | ref\|NC_001136\|:971649 | C | A |
| Na-10 | ref\|NC_001137\|:399034 | T | G |
| Na-10 | ref\|NC_001141\|:76855 | C | A |
| Na-10 | ref\|NC_001146\|:303870 | G | T |
| Na-11 | ref\|NC_001137\|:571010 | G | A |
| Na-11 | ref\|NC_001139\|:115468 | G | T |
| Na-11 | ref\|NC_001139\|:481367 | G | C |
| Na-11 | ref\|NC_001139\|:682392 | C | T |
| Na-11 | ref\|NC_001143\|:380937 | GCTGCTGTTGCTGCAA | G |
| Na-11 | ref\|NC_001143\|:594316 | G | C |
| Na-11 | ref\|NC_001144\|:55778 | T | A |
| Na-11 | ref\|NC_001145\|:505006 | C | T |
| Na-11 | ref\|NC_001146\|:516614 | T | C |
| Na-11 | ref\|NC_001147\|:789539 | G | GA |
| Na-11 | ref\|NC_001148\|:465845 | G | T |
| Na-12 | ref\|NC_001134\|:457098 | G | A |
| Na-12 | ref\|NC_001135\|:264574 | C | A |
| Na-12 | ref\|NC_001136\|:386154 | CA | C |
| Na-12 | ref\|NC_001136\|:643209 | A | C |
| Na-12 | ref\|NC_001136\|:1418045 | C | T |
| Na-12 | ref\|NC_001137\|:258930 | G | A |
| Na-12 | ref\|NC_001138\|:95460 | AT | A |
| Na-12 | ref\|NC_001138\|:212294 | C | A |
| Na-12 | ref\|NC_001139\|:20016 | TA | T |
| Na-12 | ref\|NC_001139\|:94089 | T | A |
| Na-12 | ref\|NC_001140\|:401677 | T | A |
| Na-12 | ref\|NC_001141\|:224491 | C | CT |
| Na-12 | ref\|NC_001143\|:21369 | C | A |
| Na-12 | ref\|NC_001144\|:571048 | C | T |
| Na-12 | ref\|NC_001145\|:245959 | C | CT |
| Na-12 | ref\|NC_001146\|:71382 | G | GT |
| Na-12 | ref\|NC_001146\|:96496 | C | A |
| Na-12 | ref\|NC_001147\|:213812 | A | AT |
| Na-12 | ref\|NC_001147\|:680801 | A | AT |
| Na-12 | ref\|NC_001147\|:717918 | C | A |
| Na-12 | ref\|NC_001147\|:824938 | C | A |
| Na-12 | ref\|NC_001147\|:899844 | G | T |
| Na-13 | ref\|NC_001133\|:49 | C | CA |
| Na-13 | ref\|NC_001134\|:650261 | C | CAT |
| Na-13 | ref\|NC_001135\|:98731 | G | A |
| Na-13 | ref\|NC_001135\|:142730 | C | A |
| Na-13 | ref\|NC_001135\|:201268 | G | T |
| Na-13 | ref\|NC_001136\|:148065 | G | A |
| Na-13 | ref\|NC_001136\|:1443188 | G | A |
| Na-13 | ref\|NC_001138\|:55234 | TC | T |
| Na-13 | ref\|NC_001140\|:277095 | G | T |
| Na-13 | ref\|NC_001143\|:645059 | C | A |
| Na-13 | ref\|NC_001146\|:10201 | G | A |
| Na-13 | ref\|NC_001148\|:428348 | G | T |
| Na-13 | ref\|NC_001148\|:599170 | G | A |
| Na-14 | ref\|NC_001133\|:49847 | T | C |
| Na-14 | ref\|NC_001136\|:257621 | G | GA |
| Na-14 | ref\|NC_001136\|:836535 | C | T |
| Na-14 | ref\|NC_001136\|:1006776 | C | A |
| Na-14 | ref\|NC_001136\|:1212805 | A | G |
| Na-14 | ref\|NC_001137\|:185991 | A | G |
| Na-14 | ref\|NC_001139\|:298511 | T | G |
| Na-14 | ref\|NC_001139\|:326735 | A | AT |
| Na-14 | ref\|NC_001139\|:340278 | C | T |
| Na-14 | ref\|NC_001139\|:594202 | G | C |
| Na-14 | ref\|NC_001140\|:445130 | C | A |
| Na-14 | ref\|NC_001144\|:179891 | G | A |
| Na-14 | ref\|NC_001144\|:699735 | G | A |
| Na-14 | ref\|NC_001146\|:34671 | C | T |
| Na-14 | ref\|NC_001147\|:867894 | C | T |
| Na-14 | ref\|NC_001147\|:929597 | G | T |
| Na-14 | ref\|NC_001147\|:1036136 | C | CTGT |
| Na-14 | ref\|NC_001147\|:1078302 | C | CT |
| Na-14 | ref\|NC_001148\|:701092 | G | A |
| Na-14 | ref\|NC_001148\|:746633 | C | A |
| Na-15 | ref\|NC_001135\|:312415 | C | T |
| Na-15 | ref\|NC_001136\|:1243687 | CTGT | C |
| Na-15 | ref\|NC_001136\|:1403816 | C | A |
| Na-15 | ref\|NC_001139\|:774391 | G | T |
| Na-15 | ref\|NC_001139\|:1030704 | C | A |
| Na-15 | ref\|NC_001142\|:247804 | G | T |
| Na-15 | ref\|NC_001145\|:371798 | A | AT |
| Na-15 | ref\|NC_001146\|:95729 | A | C |
| Na-15 | ref\|NC_001146\|:125095 | T | C |
| Na-15 | ref\|NC_001146\|:216974 | C | A |
| Na-15 | ref\|NC_001146\|:765735 | C | A |
| Na-15 | ref\|NC_001147\|:427962 | A | ATCGTCGTCG |
| Na-15 | ref\|NC_001148\|:165800 | G | GTATATATAGACATACACACACACATATA |
| Na-16 | ref\|NC_001134\|:569222 | C | A |
| Na-16 | ref\|NC_001135\|:308240 | C | CT |
| Na-16 | ref\|NC_001136\|:117 | C | T |
| Na-16 | ref\|NC_001136\|:23198 | A | G |
| Na-16 | ref\|NC_001136\|:83576 | C | A |
| Na-16 | ref\|NC_001136\|:1451068 | C | A |
| Na-16 | ref\|NC_001136\|:1517680 | G | GT |
| Na-16 | ref\|NC_001138\|:95067 | G | GAT |
| Na-16 | ref\|NC_001139\|:736084 | C | A |
| Na-16 | ref\|NC_001142\|:108949 | C | A |
| Na-16 | ref\|NC_001142\|:123772 | G | A |
| Na-16 | ref\|NC_001144\|:864879 | G | T |
| Na-16 | ref\|NC_001146\|:467812 | G | T |
| Na-16 | ref\|NC_001147\|:711615 | G | GT |
| Na-16 | ref\|NC_001147\|:1045053 | T | G |
| Na-16 | ref\|NC_001148\|:265844 | A | G |
| Na-16 | ref\|NC_001148\|:356678 | A | T |
| Na-17 | ref\|NC_001136\|:66048 | G | GA |
| Na-17 | ref\|NC_001136\|:1314268 | C | A |
| Na-17 | ref\|NC_001137\|:490211 | T | TTGTTCA |
| Na-17 | ref\|NC_001139\|:82693 | G | A |
| Na-17 | ref\|NC_001139\|:663630 | G | T |
| Na-17 | ref\|NC_001139\|:729387 | T | A |
| Na-17 | ref\|NC_001139\|:729518 | T | G |
| Na-17 | ref\|NC_001139\|:1037082 | G | T |
| Na-17 | ref\|NC_001141\|:424346 | A | T |
| Na-17 | ref\|NC_001141\|:427307 | C | CT |
| Na-17 | ref\|NC_001142\|:161835 | T | TA |
| Na-17 | ref\|NC_001143\|:74602 | C | A |
| Na-17 | ref\|NC_001143\|:435731 | T | C |
| Na-17 | ref\|NC_001144\|:636416 | T | TA |
| Na-17 | ref\|NC_001145\|:244094 | G | T |
| Na-17 | ref\|NC_001146\|:243612 | G | T |
| Na-17 | ref\|NC_001148\|:286097 | A | G |
| Na-17 | ref\|NC_001148\|:385381 | C | A |
| Na-18 | ref\|NC_001136\|:19003 | T | TA |
| Na-18 | ref\|NC_001137\|:100112 | C | A |
| Na-18 | ref\|NC_001139\|:825331 | G | T |
| Na-18 | ref\|NC_001141\|:169788 | T | TC |
| Na-18 | ref\|NC_001143\|:527014 | T | A |
| Na-18 | ref\|NC_001143\|:536453 | C | A |
| Na-18 | ref\|NC_001143\|:642295 | G | T |
| Na-18 | ref\|NC_001147\|:369051 | G | T |
| Na-18 | ref\|NC_001147\|:732908 | AACC | A |
| Na-18 | ref\|NC_001148\|:408413 | G | T |
| Na-19 | ref\|NC_001134\|:761301 | C | A |
| Na-19 | ref\|NC_001135\|:12333 | A | AAT |
| Na-19 | ref\|NC_001136\|:705666 | C | T |
| Na-19 | ref\|NC_001136\|:1003976 | A | C |
| Na-19 | ref\|NC_001136\|:1423170 | C | A |
| Na-19 | ref\|NC_001137\|:297233 | G | T |
| Na-19 | ref\|NC_001139\|:174596 | A | C |
| Na-19 | ref\|NC_001139\|:908890 | GT | G |
| Na-19 | ref\|NC_001140\|:530031 | G | T |
| Na-19 | ref\|NC_001143\|:660577 | C | CA |
| Na-19 | ref\|NC_001144\|:844060 | C | A |
| Na-19 | ref\|NC_001145\|:234617 | G | GA |
| Na-19 | ref\|NC_001147\|:525904 | C | A |
| Na-19 | ref\|NC_001148\|:165903 | CCTT | C |
| Na-2 | ref\|NC_001134\|:51498 | G | T |
| Na-2 | ref\|NC_001135\|:2235 | G | T |
| Na-2 | ref\|NC_001136\|:357044 | G | GA |
| Na-2 | ref\|NC_001136\|:731266 | C | T |
| Na-2 | ref\|NC_001136\|:818116 | T | TC |
| Na-2 | ref\|NC_001142\|:47467 | G | GT |
| Na-2 | ref\|NC_001142\|:55159 | C | T |
| Na-2 | ref\|NC_001142\|:246504 | A | C |
| Na-2 | ref\|NC_001142\|:533014 | T | G |
| Na-2 | ref\|NC_001144\|:489466 | A | G |
| Na-2 | ref\|NC_001146\|:710634 | G | A |
| Na-20 | ref\|NC_001134\|:307442 | T | TA |
| Na-20 | ref\|NC_001134\|:386726 | GGAA | G |
| Na-20 | ref\|NC_001134\|:681870 | T | TA |
| Na-20 | ref\|NC_001135\|:300882 | T | C |
| Na-20 | ref\|NC_001136\|:331323 | C | T |
| Na-20 | ref\|NC_001136\|:439033 | C | A |
| Na-20 | ref\|NC_001136\|:797407 | C | T |
| Na-20 | ref\|NC_001136\|:1346136 | A | G |
| Na-20 | ref\|NC_001139\|:40640 | C | A |
| Na-20 | ref\|NC_001139\|:78853 | G | T |
| Na-20 | ref\|NC_001139\|:231256 | C | T |
| Na-20 | ref\|NC_001139\|:393488 | G | C |
| Na-20 | ref\|NC_001140\|:11865 | C | CT |
| Na-20 | ref\|NC_001140\|:225000 | C | T |
| Na-20 | ref\|NC_001140\|:252331 | C | A |
| Na-20 | ref\|NC_001141\|:66107 | G | GT |
| Na-20 | ref\|NC_001141\|:101227 | AGTT | A |
| Na-20 | ref\|NC_001142\|:110763 | T | C |
| Na-20 | ref\|NC_001142\|:317080 | G | GTA |
| Na-20 | ref\|NC_001142\|:426329 | T | G |
| Na-20 | ref\|NC_001142\|:723631 | A | T |
| Na-20 | ref\|NC_001143\|:121603 | C | A |
| Na-20 | ref\|NC_001143\|:536453 | C | A |
| Na-20 | ref\|NC_001145\|:426216 | CT | C |
| Na-20 | ref\|NC_001145\|:705350 | T | G |
| Na-20 | ref\|NC_001146\|:688619 | T | G |
| Na-20 | ref\|NC_001147\|:131263 | G | A |
| Na-20 | ref\|NC_001147\|:624205 | C | T |
| Na-20 | ref\|NC_001147\|:912827 | C | A |
| Na-20 | ref\|NC_001148\|:12909 | C | CT |
| Na-20 | ref\|NC_001148\|:319892 | CA | C |
| Na-20 | ref\|NC_001148\|:536707 | T | TTTC |
| Na-21 | ref\|NC_001134\|:117303 | T | A |
| Na-21 | ref\|NC_001136\|:38423 | C | A |
| Na-21 | ref\|NC_001136\|:427112 | T | A |
| Na-21 | ref\|NC_001136\|:665217 | A | AT |
| Na-21 | ref\|NC_001136\|:679129 | T | A |
| Na-21 | ref\|NC_001136\|:750638 | C | T |
| Na-21 | ref\|NC_001137\|:98421 | C | A |
| Na-21 | ref\|NC_001141\|:33181 | C | T |
| Na-21 | ref\|NC_001143\|:168995 | T | TAATA |
| Na-21 | ref\|NC_001147\|:8683 | T | A |
| Na-21 | ref\|NC_001147\|:539586 | G | A |
| Na-22 | ref\|NC_001134\|:541458 | G | GTCA |
| Na-22 | ref\|NC_001134\|:678281 | G | T |
| Na-22 | ref\|NC_001134\|:775032 | C | T |
| Na-22 | ref\|NC_001135\|:83153 | C | G |
| Na-22 | ref\|NC_001136\|:3263 | C | A |
| Na-22 | ref\|NC_001136\|:195056 | C | A |
| Na-22 | ref\|NC_001136\|:484308 | A | G |
| Na-22 | ref\|NC_001136\|:785156 | C | A |
| Na-22 | ref\|NC_001136\|:1308607 | T | C |
| Na-22 | ref\|NC_001136\|:1516080 | G | A |
| Na-22 | ref\|NC_001137\|:468982 | C | CT |
| Na-22 | ref\|NC_001139\|:748809 | T | G |
| Na-22 | ref\|NC_001142\|:71276 | G | GA |
| Na-22 | ref\|NC_001144\|:129214 | C | CA |
| Na-22 | ref\|NC_001144\|:624650 | T | A |
| Na-22 | ref\|NC_001144\|:841499 | G | T |
| Na-22 | ref\|NC_001144\|:932259 | G | T |
| Na-22 | ref\|NC_001144\|:1066450 | C | T |
| Na-22 | ref\|NC_001145\|:130895 | A | AGT |
| Na-22 | ref\|NC_001145\|:762085 | C | A |
| Na-22 | ref\|NC_001146\|:18775 | G | A |
| Na-22 | ref\|NC_001146\|:650099 | G | C |
| Na-22 | ref\|NC_001146\|:751566 | C | T |
| Na-22 | ref\|NC_001147\|:971014 | A | G |
| Na-22 | ref\|NC_001148\|:520952 | T | TATAATA |
| Na-22 | ref\|NC_001148\|:776934 | T | TTTTC |
| Na-23 | ref\|NC_001133\|:38777 | C | A |
| Na-23 | ref\|NC_001134\|:56139 | A | G |
| Na-23 | ref\|NC_001134\|:680588 | T | G |
| Na-23 | ref\|NC_001136\|:258461 | T | C |
| Na-23 | ref\|NC_001136\|:639207 | C | G |
| Na-23 | ref\|NC_001136\|:690502 | G | T |
| Na-23 | ref\|NC_001139\|:669746 | G | T |
| Na-23 | ref\|NC_001139\|:1077212 | A | T |
| Na-23 | ref\|NC_001140\|:139906 | C | T |
| Na-23 | ref\|NC_001142\|:256810 | C | CAT |
| Na-23 | ref\|NC_001142\|:618031 | A | AT |
| Na-23 | ref\|NC_001143\|:227752 | C | A |
| Na-23 | ref\|NC_001144\|:437594 | G | A |
| Na-23 | ref\|NC_001144\|:656938 | C | A |
| Na-23 | ref\|NC_001145\|:156819 | C | A |
| Na-23 | ref\|NC_001146\|:290022 | CGAAGACATCGGT | C |
| Na-23 | ref\|NC_001147\|:549366 | C | T |
| Na-24 | ref\|NC_001134\|:407170 | G | T |
| Na-24 | ref\|NC_001136\|:42002 | C | T |
| Na-24 | ref\|NC_001136\|:80724 | C | T |
| Na-24 | ref\|NC_001136\|:830928 | C | A |
| Na-24 | ref\|NC_001136\|:1273155 | T | C |
| Na-24 | ref\|NC_001139\|:173516 | T | TG |
| Na-24 | ref\|NC_001139\|:203076 | C | T |
| Na-24 | ref\|NC_001140\|:66399 | G | A |
| Na-24 | ref\|NC_001142\|:405874 | A | G |
| Na-24 | ref\|NC_001144\|:1007265 | G | C |
| Na-24 | ref\|NC_001146\|:777869 | G | A |
| Na-24 | ref\|NC_001147\|:594379 | G | A |
| Na-24 | ref\|NC_001148\|:485926 | T | TCACA |
| Na-24 | ref\|NC_001148\|:799409 | C | A |
| Na-24 | ref\|NC_001148\|:908978 | C | A |
| Na-3 | ref\|NC_001134\|:733289 | A | T |
| Na-3 | ref\|NC_001136\|:797149 | C | A |
| Na-3 | ref\|NC_001139\|:502764 | C | CT |
| Na-3 | ref\|NC_001142\|:588442 | C | A |
| Na-3 | ref\|NC_001143\|:661329 | G | T |
| Na-3 | ref\|NC_001145\|:705604 | T | G |
| Na-3 | ref\|NC_001146\|:98905 | G | T |
| Na-3 | ref\|NC_001146\|:372894 | C | A |
| Na-3 | ref\|NC_001147\|:848307 | G | GT |
| Na-3 | ref\|NC_001148\|:105500 | AT | A |
| Na-4 | ref\|NC_001136\|:1475501 | T | G |
| Na-4 | ref\|NC_001137\|:359426 | G | T |
| Na-4 | ref\|NC_001140\|:411057 | T | C |
| Na-4 | ref\|NC_001143\|:531857 | G | T |
| Na-4 | ref\|NC_001144\|:199814 | C | T |
| Na-4 | ref\|NC_001144\|:908097 | C | T |
| Na-4 | ref\|NC_001146\|:509211 | G | T |
| Na-4 | ref\|NC_001146\|:724094 | C | T |
| Na-4 | ref\|NC_001147\|:916 | G | T |
| Na-4 | ref\|NC_001147\|:1005297 | G | A |
| Na-4 | ref\|NC_001148\|:348398 | G | GA |
| Na-5 | ref\|NC_001134\|:429501 | A | G |
| Na-5 | ref\|NC_001134\|:749407 | C | T |
| Na-5 | ref\|NC_001134\|:772403 | G | T |
| Na-5 | ref\|NC_001136\|:54046 | T | TA |
| Na-5 | ref\|NC_001136\|:71600 | C | A |
| Na-5 | ref\|NC_001137\|:81191 | GTTGTTATTATTATTATTA | G |
| Na-5 | ref\|NC_001139\|:67689 | G | A |
| Na-5 | ref\|NC_001139\|:215260 | G | GA |
| Na-5 | ref\|NC_001141\|:109645 | G | T |
| Na-5 | ref\|NC_001144\|:865993 | G | T |
| Na-5 | ref\|NC_001146\|:29703 | T | C |
| Na-5 | ref\|NC_001147\|:12726 | T | A |
| Na-5 | ref\|NC_001147\|:620467 | A | G |
| Na-5 | ref\|NC_001148\|:339856 | G | T |
| Na-5 | ref\|NC_001148\|:936712 | G | A |
| Na-6 | ref\|NC_001134\|:541608 | T | TTTA |
| Na-6 | ref\|NC_001134\|:714132 | T | TTCTTCGTCTTCG |
| Na-6 | ref\|NC_001135\|:67752 | C | G |
| Na-6 | ref\|NC_001136\|:447179 | G | T |
| Na-6 | ref\|NC_001136\|:691224 | G | GA |
| Na-6 | ref\|NC_001136\|:762336 | A | T |
| Na-6 | ref\|NC_001137\|:514642 | C | T |
| Na-6 | ref\|NC_001139\|:532378 | G | T |
| Na-6 | ref\|NC_001139\|:541813 | T | C |
| Na-6 | ref\|NC_001139\|:669888 | C | A |
| Na-6 | ref\|NC_001139\|:965160 | C | A |
| Na-6 | ref\|NC_001140\|:565 | C | T |
| Na-6 | ref\|NC_001141\|:248934 | AT | A |
| Na-6 | ref\|NC_001142\|:416084 | C | A |
| Na-6 | ref\|NC_001142\|:548426 | C | G |
| Na-6 | ref\|NC_001143\|:604319 | T | G |
| Na-6 | ref\|NC_001143\|:662546 | G | T |
| Na-6 | ref\|NC_001144\|:599939 | C | T |
| Na-6 | ref\|NC_001144\|:898743 | C | A |
| Na-6 | ref\|NC_001144\|:978657 | C | A |
| Na-6 | ref\|NC_001145\|:339947 | A | C |
| Na-6 | ref\|NC_001147\|:420953 | T | C |
| Na-6 | ref\|NC_001147\|:622232 | G | A |
| Na-6 | ref\|NC_001148\|:44428 | G | GT |
| Na-6 | ref\|NC_001148\|:154750 | ATC | A |
| Na-7 | ref\|NC_001134\|:229142 | C | T |
| Na-7 | ref\|NC_001137\|:517521 | A | C |
| Na-7 | ref\|NC_001138\|:224651 | CCTT | C |
| Na-7 | ref\|NC_001139\|:415228 | C | CA |
| Na-7 | ref\|NC_001139\|:727060 | T | TA |
| Na-7 | ref\|NC_001143\|:420495 | G | T |
| Na-7 | ref\|NC_001143\|:644998 | G | GA |
| Na-7 | ref\|NC_001144\|:806443 | CTGT | C |
| Na-7 | ref\|NC_001145\|:734978 | T | C |
| Na-7 | ref\|NC_001146\|:164924 | C | T |
| Na-7 | ref\|NC_001147\|:602488 | A | AT |
| Na-7 | ref\|NC_001148\|:280246 | A | C |
| Na-7 | ref\|NC_001148\|:783275 | C | A |
| Na-8 | ref\|NC_001133\|:172008 | G | GA |
| Na-8 | ref\|NC_001134\|:347603 | G | A |
| Na-8 | ref\|NC_001134\|:719695 | C | G |
| Na-8 | ref\|NC_001136\|:144003 | C | A |
| Na-8 | ref\|NC_001136\|:1408890 | A | G |
| Na-8 | ref\|NC_001137\|:527499 | C | A |
| Na-8 | ref\|NC_001140\|:60273 | G | T |
| Na-8 | ref\|NC_001144\|:333391 | G | T |
| Na-8 | ref\|NC_001144\|:536513 | G | T |
| Na-8 | ref\|NC_001144\|:1054376 | G | T |
| Na-8 | ref\|NC_001145\|:513261 | G | T |
| Na-8 | ref\|NC_001147\|:617735 | A | T |
| Na-8 | ref\|NC_001147\|:655439 | TA | T |
| Na-8 | ref\|NC_001147\|:803670 | C | T |
| Na-8 | ref\|NC_001148\|:168215 | T | G |
| Na-8 | ref\|NC_001148\|:270872 | T | TG |
| Na-8 | ref\|NC_001148\|:474027 | CA | C |
| Na-9 | ref\|NC_001134\|:72395 | G | A |
| Na-9 | ref\|NC_001136\|:246433 | T | C |
| Na-9 | ref\|NC_001136\|:355508 | C | CT |
| Na-9 | ref\|NC_001136\|:1376066 | G | A |
| Na-9 | ref\|NC_001137\|:11309 | C | A |
| Na-9 | ref\|NC_001137\|:511754 | T | G |
| Na-9 | ref\|NC_001138\|:80279 | T | A |
| Na-9 | ref\|NC_001138\|:107244 | C | A |
| Na-9 | ref\|NC_001139\|:482120 | TAAC | T |
| Na-9 | ref\|NC_001139\|:1050205 | A | C |
| Na-9 | ref\|NC_001140\|:518678 | G | A |
| Na-9 | ref\|NC_001142\|:245109 | T | G |
| Na-9 | ref\|NC_001143\|:176286 | A | C |
| Na-9 | ref\|NC_001143\|:577899 | G | T |
| Na-9 | ref\|NC_001145\|:434662 | C | T |
| Na-9 | ref\|NC_001146\|:262637 | C | T |
| YNB-1 | ref\|NC_001134\|:36513 | T | G |
| YNB-1 | ref\|NC_001136\|:24249 | T | C |
| YNB-1 | ref\|NC_001136\|:779770 | C | T |
| YNB-1 | ref\|NC_001136\|:1431415 | G | T |
| YNB-1 | ref\|NC_001137\|:100197 | C | A |
| YNB-1 | ref\|NC_001138\|:226632 | G | T |
| YNB-1 | ref\|NC_001148\|:198911 | G | A |
| YNB-10 | ref\|NC_001136\|:1268404 | C | A |
| YNB-10 | ref\|NC_001136\|:1390505 | A | C |
| YNB-10 | ref\|NC_001137\|:32959 | C | T |
| YNB-10 | ref\|NC_001142\|:623890 | G | T |
| YNB-10 | ref\|NC_001145\|:247423 | G | T |
| YNB-10 | ref\|NC_001148\|:530890 | G | T |
| YNB-11 | ref\|NC_001135\|:180552 | A | T |
| YNB-11 | ref\|NC_001139\|:927431 | T | C |
| YNB-11 | ref\|NC_001145\|:490129 | A | C |
| YNB-11 | ref\|NC_001148\|:205738 | G | T |
| YNB-12 | ref\|NC_001134\|:681668 | A | T |
| YNB-12 | ref\|NC_001136\|:244108 | G | T |
| YNB-12 | ref\|NC_001136\|:307127 | T | C |
| YNB-12 | ref\|NC_001136\|:575820 | G | A |
| YNB-12 | ref\|NC_001136\|:1138438 | G | T |
| YNB-12 | ref\|NC_001137\|:492341 | GA | G |
| YNB-12 | ref\|NC_001138\|:249537 | G | T |
| YNB-12 | ref\|NC_001139\|:240071 | T | G |
| YNB-12 | ref\|NC_001139\|:501500 | A | T |
| YNB-12 | ref\|NC_001140\|:318125 | T | G |
| YNB-12 | ref\|NC_001143\|:190651 | C | A |
| YNB-12 | ref\|NC_001143\|:379921 | GA | G |
| YNB-12 | ref\|NC_001144\|:179740 | A | C |
| YNB-12 | ref\|NC_001146\|:311609 | G | T |
| YNB-12 | ref\|NC_001146\|:399073 | C | A |
| YNB-12 | ref\|NC_001147\|:358297 | C | A |
| YNB-13 | ref\|NC_001140\|:16567 | T | C |
| YNB-13 | ref\|NC_001146\|:167625 | C | T |
| YNB-13 | ref\|NC_001147\|:236286 | G | GCAAGCACAAGCA |
| YNB-13 | ref\|NC_001148\|:406795 | G | T |
| YNB-14 | ref\|NC_001142\|:139864 | C | G |
| YNB-14 | ref\|NC_001142\|:393836 | A | T |
| YNB-14 | ref\|NC_001142\|:641182 | G | A |
| YNB-14 | ref\|NC_001146\|:312805 | T | G |
| YNB-14 | ref\|NC_001146\|:702574 | A | G |
| YNB-14 | ref\|NC_001147\|:822412 | A | G |
| YNB-14 | ref\|NC_001148\|:369402 | G | A |
| YNB-14 | ref\|NC_001148\|:636509 | C | A |
| YNB-15 | ref\|NC_001134\|:156469 | G | A |
| YNB-15 | ref\|NC_001134\|:740400 | A | C |
| YNB-15 | ref\|NC_001136\|:1242577 | G | A |
| YNB-15 | ref\|NC_001137\|:305250 | T | A |
| YNB-15 | ref\|NC_001138\|:76169 | AT | A |
| YNB-15 | ref\|NC_001139\|:679899 | T | C |
| YNB-15 | ref\|NC_001140\|:450837 | C | A |
| YNB-15 | ref\|NC_001144\|:498569 | C | A |
| YNB-15 | ref\|NC_001144\|:875444 | G | T |
| YNB-15 | ref\|NC_001144\|:1031695 | G | C |
| YNB-16 | ref\|NC_001136\|:149113 | A | AAAATAAATAAAT |
| YNB-16 | ref\|NC_001136\|:1282640 | T | C |
| YNB-16 | ref\|NC_001139\|:290682 | G | A |
| YNB-16 | ref\|NC_001140\|:381232 | T | G |
| YNB-16 | ref\|NC_001142\|:200136 | T | A |
| YNB-16 | ref\|NC_001142\|:252575 | G | A |
| YNB-16 | ref\|NC_001142\|:504909 | A | G |
| YNB-16 | ref\|NC_001142\|:693833 | C | A |
| YNB-16 | ref\|NC_001145\|:266104 | C | A |
| YNB-16 | ref\|NC_001146\|:631844 | TG | T |
| YNB-16 | ref\|NC_001147\|:581360 | A | C |
| YNB-16 | ref\|NC_001148\|:97284 | T | C |
| YNB-17 | ref\|NC_001136\|:806170 | G | A |
| YNB-17 | ref\|NC_001137\|:71441 | G | A |
| YNB-17 | ref\|NC_001139\|:693473 | A | C |
| YNB-17 | ref\|NC_001139\|:882470 | G | T |
| YNB-17 | ref\|NC_001142\|:3537 | C | A |
| YNB-17 | ref\|NC_001144\|:1039201 | A | T |
| YNB-17 | ref\|NC_001145\|:178957 | G | A |
| YNB-17 | ref\|NC_001145\|:579498 | G | T |
| YNB-17 | ref\|NC_001148\|:695367 | C | T |
| YNB-18 | ref\|NC_001133\|:226044 | G | T |
| YNB-18 | ref\|NC_001134\|:402294 | G | T |
| YNB-18 | ref\|NC_001134\|:464217 | CGCTTGT | C |
| YNB-18 | ref\|NC_001136\|:687284 | C | A |
| YNB-18 | ref\|NC_001137\|:351630 | TG | T |
| YNB-18 | ref\|NC_001139\|:243372 | A | T |
| YNB-18 | ref\|NC_001139\|:739200 | C | A |
| YNB-18 | ref\|NC_001142\|:192938 | C | T |
| YNB-18 | ref\|NC_001144\|:213640 | T | G |
| YNB-18 | ref\|NC_001144\|:1065086 | T | G |
| YNB-18 | ref\|NC_001145\|:85499 | A | C |
| YNB-18 | ref\|NC_001145\|:381205 | G | T |
| YNB-18 | ref\|NC_001146\|:369180 | G | T |
| YNB-18 | ref\|NC_001148\|:536501 | G | T |
| YNB-19 | ref\|NC_001133\|:9794 | T | G |
| YNB-19 | ref\|NC_001136\|:303010 | AT | A |
| YNB-19 | ref\|NC_001138\|:167855 | A | AT |
| YNB-19 | ref\|NC_001139\|:889860 | T | C |
| YNB-19 | ref\|NC_001140\|:541009 | C | T |
| YNB-19 | ref\|NC_001142\|:525346 | G | T |
| YNB-19 | ref\|NC_001144\|:59747 | T | TCTATATTGAAAAGTAAAC |
| YNB-19 | ref\|NC_001145\|:816162 | G | T |
| YNB-19 | ref\|NC_001146\|:574263 | A | G |
| YNB-2 | ref\|NC_001139\|:328564 | A | G |
| YNB-2 | ref\|NC_001142\|:592317 | C | G |
| YNB-2 | ref\|NC_001143\|:290334 | G | T |
| YNB-2 | ref\|NC_001144\|:643138 | G | C |
| YNB-2 | ref\|NC_001146\|:428892 | T | G |
| YNB-2 | ref\|NC_001148\|:898820 | C | A |
| YNB-20 | ref\|NC_001134\|:235501 | G | C |
| YNB-20 | ref\|NC_001134\|:290421 | G | T |
| YNB-20 | ref\|NC_001136\|:1407766 | G | T |
| YNB-20 | ref\|NC_001139\|:703178 | G | A |
| YNB-20 | ref\|NC_001141\|:277131 | A | C |
| YNB-20 | ref\|NC_001143\|:61204 | G | T |
| YNB-20 | ref\|NC_001143\|:567597 | G | T |
| YNB-20 | ref\|NC_001145\|:31785 | C | A |
| YNB-20 | ref\|NC_001146\|:499567 | T | A |
| YNB-20 | ref\|NC_001148\|:879177 | T | C |
| YNB-21 | ref\|NC_001135\|:137354 | C | T |
| YNB-21 | ref\|NC_001136\|:349669 | A | G |
| YNB-21 | ref\|NC_001136\|:954363 | C | G |
| YNB-21 | ref\|NC_001137\|:6917 | T | G |
| YNB-21 | ref\|NC_001137\|:367860 | T | G |
| YNB-21 | ref\|NC_001138\|:209252 | T | G |
| YNB-21 | ref\|NC_001139\|:1074114 | A | G |
| YNB-21 | ref\|NC_001140\|:168780 | G | GTA |
| YNB-21 | ref\|NC_001140\|:190310 | G | T |
| YNB-21 | ref\|NC_001141\|:43046 | A | G |
| YNB-21 | ref\|NC_001143\|:296871 | G | A |
| YNB-21 | ref\|NC_001144\|:44335 | G | T |
| YNB-21 | ref\|NC_001144\|:769867 | T | C |
| YNB-21 | ref\|NC_001146\|:110236 | C | A |
| YNB-21 | ref\|NC_001146\|:620025 | C | T |
| YNB-21 | ref\|NC_001147\|:240370 | G | T |
| YNB-21 | ref\|NC_001147\|:381141 | A | G |
| YNB-22 | ref\|NC_001134\|:303639 | A | G |
| YNB-22 | ref\|NC_001136\|:205078 | C | G |
| YNB-22 | ref\|NC_001136\|:367645 | G | A |
| YNB-22 | ref\|NC_001138\|:89056 | G | T |
| YNB-22 | ref\|NC_001141\|:336701 | T | TC |
| YNB-22 | ref\|NC_001142\|:163481 | T | C |
| YNB-22 | ref\|NC_001143\|:76503 | C | A |
| YNB-22 | ref\|NC_001144\|:839370 | C | G |
| YNB-22 | ref\|NC_001147\|:109202 | A | G |
| YNB-22 | ref\|NC_001148\|:645855 | C | T |
| YNB-23 | ref\|NC_001134\|:536463 | C | A |
| YNB-23 | ref\|NC_001138\|:88203 | G | C |
| YNB-23 | ref\|NC_001139\|:438890 | A | G |
| YNB-23 | ref\|NC_001139\|:1073218 | C | A |
| YNB-23 | ref\|NC_001140\|:100822 | C | T |
| YNB-23 | ref\|NC_001140\|:406128 | G | T |
| YNB-23 | ref\|NC_001144\|:80469 | G | T |
| YNB-23 | ref\|NC_001148\|:869777 | C | A |
| YNB-24 | ref\|NC_001136\|:154529 | T | G |
| YNB-24 | ref\|NC_001140\|:54812 | G | T |
| YNB-24 | ref\|NC_001141\|:209302 | T | G |
| YNB-24 | ref\|NC_001142\|:541188 | C | G |
| YNB-24 | ref\|NC_001143\|:228293 | G | T |
| YNB-24 | ref\|NC_001143\|:345087 | T | A |
| YNB-24 | ref\|NC_001143\|:394234 | C | T |
| YNB-24 | ref\|NC_001147\|:328501 | T | G |
| YNB-24 | ref\|NC_001148\|:161621 | T | G |
| YNB-24 | ref\|NC_001148\|:500061 | G | A |
| YNB-3 | ref\|NC_001133\|:89511 | T | A |
| YNB-3 | ref\|NC_001136\|:816702 | T | C |
| YNB-3 | ref\|NC_001137\|:290430 | T | C |
| YNB-3 | ref\|NC_001139\|:1063533 | G | T |
| YNB-3 | ref\|NC_001141\|:377845 | G | A |
| YNB-3 | ref\|NC_001143\|:333255 | G | C |
| YNB-3 | ref\|NC_001144\|:510369 | G | A |
| YNB-3 | ref\|NC_001146\|:331978 | G | A |
| YNB-4 | ref\|NC_001134\|:656811 | C | T |
| YNB-4 | ref\|NC_001140\|:79802 | G | T |
| YNB-4 | ref\|NC_001142\|:588389 | G | C |
| YNB-4 | ref\|NC_001145\|:266009 | T | C |
| YNB-4 | ref\|NC_001147\|:1057787 | C | T |
| YNB-5 | ref\|NC_001135\|:309046 | A | G |
| YNB-5 | ref\|NC_001138\|:162289 | G | T |
| YNB-5 | ref\|NC_001141\|:37437 | C | A |
| YNB-5 | ref\|NC_001142\|:645988 | A | G |
| YNB-5 | ref\|NC_001144\|:63509 | C | T |
| YNB-5 | ref\|NC_001145\|:621036 | G | A |
| YNB-6 | ref\|NC_001134\|:217318 | A | T |
| YNB-6 | ref\|NC_001134\|:401436 | G | A |
| YNB-6 | ref\|NC_001136\|:527743 | A | G |
| YNB-6 | ref\|NC_001139\|:559566 | T | A |
| YNB-6 | ref\|NC_001139\|:1067326 | C | A |
| YNB-6 | ref\|NC_001143\|:212990 | C | T |
| YNB-6 | ref\|NC_001144\|:605220 | C | T |
| YNB-6 | ref\|NC_001144\|:639803 | T | TAAG |
| YNB-6 | ref\|NC_001144\|:929562 | G | C |
| YNB-6 | ref\|NC_001145\|:251661 | G | GA |
| YNB-6 | ref\|NC_001147\|:66012 | C | A |
| YNB-6 | ref\|NC_001147\|:87072 | T | G |
| YNB-6 | ref\|NC_001147\|:590687 | A | C |
| YNB-6 | ref\|NC_001148\|:169529 | G | C |
| YNB-7 | ref\|NC_001135\|:134884 | C | A |
| YNB-7 | ref\|NC_001137\|:82945 | A | G |
| YNB-7 | ref\|NC_001140\|:241326 | C | A |
| YNB-7 | ref\|NC_001140\|:255906 | A | C |
| YNB-7 | ref\|NC_001142\|:554036 | A | T |
| YNB-7 | ref\|NC_001143\|:16601 | C | A |
| YNB-7 | ref\|NC_001145\|:337497 | C | G |
| YNB-7 | ref\|NC_001147\|:487511 | C | A |
| YNB-7 | ref\|NC_001147\|:537711 | G | A |
| YNB-7 | ref\|NC_001148\|:639646 | A | AG |
| YNB-8 | ref\|NC_001134\|:359742 | G | T |
| YNB-8 | ref\|NC_001134\|:647638 | G | A |
| YNB-8 | ref\|NC_001135\|:174931 | G | A |
| YNB-8 | ref\|NC_001140\|:518781 | G | A |
| YNB-8 | ref\|NC_001144\|:862911 | C | T |
| YNB-8 | ref\|NC_001147\|:193675 | C | CTT |
| YNB-9 | ref\|NC_001134\|:721799 | C | T |
| YNB-9 | ref\|NC_001139\|:500550 | T | C |
| YNB-9 | ref\|NC_001139\|:669031 | C | T |
| YNB-9 | ref\|NC_001141\|:372368 | C | A |
| YNB-9 | ref\|NC_001145\|:730711 | A | C |
| YNB-9 | ref\|NC_001146\|:490489 | C | T |
| YPD-1 | ref\|NC_001134\|:667718 | A | G |
| YPD-1 | ref\|NC_001136\|:331549 | G | C |
| YPD-10 | ref\|NC_001134\|:761769 | G | A |
| YPD-10 | ref\|NC_001136\|:1414938 | G | A |
| YPD-10 | ref\|NC_001136\|:1464052 | G | C |
| YPD-10 | ref\|NC_001138\|:6707 | T | C |
| YPD-10 | ref\|NC_001139\|:435270 | G | A |
| YPD-10 | ref\|NC_001140\|:35246 | A | T |
| YPD-10 | ref\|NC_001147\|:778437 | A | T |
| YPD-11 | ref\|NC_001136\|:741915 | G | T |
| YPD-11 | ref\|NC_001139\|:297781 | A | C |
| YPD-11 | ref\|NC_001142\|:635208 | C | A |
| YPD-12 | ref\|NC_001136\|:393709 | G | A |
| YPD-12 | ref\|NC_001136\|:1246397 | G | T |
| YPD-12 | ref\|NC_001138\|:210626 | G | A |
| YPD-12 | ref\|NC_001139\|:789636 | C | A |
| YPD-12 | ref\|NC_001140\|:149740 | A | G |
| YPD-12 | ref\|NC_001146\|:704606 | C | A |
| YPD-12 | ref\|NC_001148\|:866449 | C | T |
| YPD-13 | ref\|NC_001139\|:127918 | T | C |
| YPD-13 | ref\|NC_001144\|:427944 | C | T |
| YPD-14 | ref\|NC_001134\|:672300 | C | G |
| YPD-14 | ref\|NC_001136\|:375741 | G | T |
| YPD-14 | ref\|NC_001136\|:488899 | A | C |
| YPD-14 | ref\|NC_001136\|:784729 | C | T |
| YPD-14 | ref\|NC_001141\|:364925 | C | G |
| YPD-14 | ref\|NC_001147\|:923456 | C | T |
| YPD-15 | ref\|NC_001139\|:192401 | C | A |
| YPD-15 | ref\|NC_001143\|:53397 | C | G |
| YPD-15 | ref\|NC_001143\|:98981 | C | A |
| YPD-15 | ref\|NC_001145\|:30871 | G | T |
| YPD-15 | ref\|NC_001148\|:737914 | T | C |
| YPD-16 | ref\|NC_001134\|:495787 | G | A |
| YPD-16 | ref\|NC_001136\|:427438 | CCGTGGTGGTGCTCGCGGTGGTTC  CAGAGGTGGCTTCGGTGGTAGAGG  CGGTTCT | C |
| YPD-16 | ref\|NC_001137\|:337452 | GA | G |
| YPD-16 | ref\|NC_001139\|:103493 | T | G |
| YPD-16 | ref\|NC_001139\|:501621 | A | T |
| YPD-16 | ref\|NC_001139\|:1042628 | A | G |
| YPD-17 | ref\|NC_001136\|:833384 | G | A |
| YPD-17 | ref\|NC_001139\|:541931 | T | C |
| YPD-17 | ref\|NC_001142\|:21094 | G | T |
| YPD-17 | ref\|NC_001145\|:529382 | C | A |
| YPD-18 | ref\|NC_001147\|:132023 | G | T |
| YPD-18 | ref\|NC_001148\|:403753 | C | T |
| YPD-19 | ref\|NC_001134\|:499445 | A | T |
| YPD-19 | ref\|NC_001139\|:157665 | C | T |
| YPD-19 | ref\|NC_001145\|:725589 | C | T |
| YPD-2 | ref\|NC_001134\|:139670 | T | C |
| YPD-2 | ref\|NC_001140\|:217011 | T | G |
| YPD-2 | ref\|NC_001143\|:533956 | GA | G |
| YPD-2 | ref\|NC_001144\|:374173 | C | T |
| YPD-2 | ref\|NC_001144\|:526416 | T | A |
| YPD-2 | ref\|NC_001146\|:31515 | A | C |
| YPD-20 | ref\|NC_001134\|:270163 | A | C |
| YPD-20 | ref\|NC_001136\|:319071 | T | A |
| YPD-20 | ref\|NC_001136\|:1237213 | A | C |
| YPD-20 | ref\|NC_001136\|:1268587 | G | T |
| YPD-20 | ref\|NC_001141\|:69872 | C | A |
| YPD-20 | ref\|NC_001144\|:1021040 | C | A |
| YPD-21 | ref\|NC_001136\|:779244 | G | T |
| YPD-21 | ref\|NC_001136\|:1455173 | C | T |
| YPD-21 | ref\|NC_001140\|:414656 | G | C |
| YPD-21 | ref\|NC_001145\|:544030 | C | T |
| YPD-21 | ref\|NC_001148\|:350543 | A | G |
| YPD-21 | ref\|NC_001148\|:490736 | A | C |
| YPD-22 | ref\|NC_001136\|:836873 | G | T |
| YPD-22 | ref\|NC_001139\|:370644 | A | C |
| YPD-22 | ref\|NC_001139\|:650055 | G | T |
| YPD-22 | ref\|NC_001143\|:162487 | C | A |
| YPD-22 | ref\|NC_001144\|:318088 | A | T |
| YPD-22 | ref\|NC_001144\|:394006 | T | A |
| YPD-22 | ref\|NC_001144\|:622172 | C | T |
| YPD-22 | ref\|NC_001147\|:340289 | TA | T |
| YPD-23 | ref\|NC_001140\|:230268 | A | G |
| YPD-23 | ref\|NC_001143\|:364030 | T | G |
| YPD-23 | ref\|NC_001147\|:616886 | C | A |
| YPD-24 | ref\|NC_001141\|:130960 | T | C |
| YPD-24 | ref\|NC_001145\|:283910 | T | G |
| YPD-24 | ref\|NC_001147\|:698839 | C | A |
| YPD-24 | ref\|NC_001147\|:978149 | CA | C |
| YPD-24 | ref\|NC_001147\|:1020404 | T | C |
| YPD-3 | ref\|NC_001133\|:34907 | G | A |
| YPD-3 | ref\|NC_001140\|:487918 | C | A |
| YPD-3 | ref\|NC_001143\|:244107 | A | G |
| YPD-3 | ref\|NC_001144\|:284407 | T | C |
| YPD-3 | ref\|NC_001144\|:997886 | G | T |
| YPD-4 | ref\|NC_001137\|:553904 | A | C |
| YPD-4 | ref\|NC_001140\|:415705 | G | T |
| YPD-4 | ref\|NC_001143\|:593175 | G | A |
| YPD-4 | ref\|NC_001148\|:39131 | C | G |
| YPD-5 | ref\|NC_001139\|:1054123 | T | A |
| YPD-5 | ref\|NC_001142\|:404098 | A | G |
| YPD-5 | ref\|NC_001145\|:65915 | C | CTGTTGTCCATTCTGTTGTTGTTGT |
| YPD-5 | ref\|NC_001146\|:736000 | G | T |
| YPD-6 | ref\|NC_001135\|:289129 | C | G |
| YPD-6 | ref\|NC_001136\|:1187503 | A | T |
| YPD-6 | ref\|NC_001141\|:209110 | C | A |
| YPD-6 | ref\|NC_001142\|:690754 | G | A |
| YPD-6 | ref\|NC_001144\|:683769 | T | C |
| YPD-6 | ref\|NC_001144\|:971150 | G | T |
| YPD-6 | ref\|NC_001146\|:768348 | G | T |
| YPD-7 | ref\|NC_001136\|:91781 | C | A |
| YPD-7 | ref\|NC_001137\|:191857 | A | G |
| YPD-7 | ref\|NC_001138\|:268805 | T | A |
| YPD-7 | ref\|NC_001139\|:467706 | C | CAAT |
| YPD-7 | ref\|NC_001142\|:73626 | C | T |
| YPD-7 | ref\|NC_001143\|:464183 | T | C |
| YPD-7 | ref\|NC_001143\|:607226 | A | G |
| YPD-7 | ref\|NC_001144\|:789707 | G | T |
| YPD-7 | ref\|NC_001147\|:684203 | T | C |
| YPD-8 | ref\|NC_001134\|:714132 | T | TTCTTCGTCTTCG |
| YPD-8 | ref\|NC_001141\|:248934 | A | AT |
| YPD-8 | ref\|NC_001144\|:179412 | C | A |
| YPD-8 | ref\|NC_001144\|:302229 | A | T |
| YPD-8 | ref\|NC_001145\|:232727 | C | G |
| YPD-8 | ref\|NC_001147\|:638908 | G | GA |
| YPD-8 | ref\|NC_001148\|:311394 | C | T |
| YPD-9 | ref\|NC_001136\|:149113 | A | AAAATAAAT |
| YPD-9 | ref\|NC_001136\|:427494 | G | A |
| YPD-9 | ref\|NC_001138\|:26021 | C | T |
| YPD-9 | ref\|NC_001139\|:951418 | T | C |
| YPD-9 | ref\|NC_001140\|:413011 | C | A |
| YPD-9 | ref\|NC_001141\|:265583 | C | T |
| YPL-1 | ref\|NC_001133\|:62383 | G | A |
| YPL-1 | ref\|NC_001136\|:71151 | G | A |
| YPL-1 | ref\|NC_001136\|:762550 | C | T |
| YPL-1 | ref\|NC_001139\|:804320 | C | G |
| YPL-1 | ref\|NC_001143\|:400076 | A | G |
| YPL-1 | ref\|NC_001143\|:456920 | C | G |
| YPL-1 | ref\|NC_001146\|:590997 | C | G |
| YPL-1 | ref\|NC_001147\|:787093 | A | C |
| YPL-10 | ref\|NC_001133\|:227367 | C | T |
| YPL-10 | ref\|NC_001136\|:1428255 | A | C |
| YPL-10 | ref\|NC_001138\|:137325 | T | TA |
| YPL-10 | ref\|NC_001138\|:192593 | C | T |
| YPL-10 | ref\|NC_001143\|:225322 | G | A |
| YPL-10 | ref\|NC_001145\|:618174 | A | C |
| YPL-11 | ref\|NC_001146\|:77959 | G | T |
| YPL-12 | ref\|NC_001136\|:741945 | TA | T |
| YPL-12 | ref\|NC_001138\|:130184 | G | A |
| YPL-12 | ref\|NC_001140\|:175997 | C | T |
| YPL-12 | ref\|NC_001144\|:1052059 | T | TA |
| YPL-12 | ref\|NC_001146\|:374220 | G | A |
| YPL-12 | ref\|NC_001148\|:878943 | G | C |
| YPL-13 | ref\|NC_001134\|:761255 | C | A |
| YPL-13 | ref\|NC_001136\|:846842 | C | A |
| YPL-13 | ref\|NC_001141\|:49999 | T | G |
| YPL-13 | ref\|NC_001141\|:367578 | C | T |
| YPL-13 | ref\|NC_001143\|:576631 | T | G |
| YPL-13 | ref\|NC_001148\|:397845 | T | G |
| YPL-14 | ref\|NC_001142\|:448126 | TA | T |
| YPL-14 | ref\|NC_001143\|:542789 | C | A |
| YPL-14 | ref\|NC_001145\|:65144 | C | A |
| YPL-14 | ref\|NC_001145\|:548187 | G | A |
| YPL-14 | ref\|NC_001146\|:288926 | G | T |
| YPL-14 | ref\|NC_001147\|:384323 | A | AT |
| YPL-15 | ref\|NC_001134\|:368557 | C | G |
| YPL-15 | ref\|NC_001134\|:674619 | T | A |
| YPL-15 | ref\|NC_001137\|:4186 | T | A |
| YPL-15 | ref\|NC_001139\|:418158 | G | A |
| YPL-15 | ref\|NC_001141\|:84917 | C | T |
| YPL-15 | ref\|NC_001142\|:80308 | A | C |
| YPL-15 | ref\|NC_001142\|:619554 | A | AT |
| YPL-15 | ref\|NC_001144\|:388254 | G | A |
| YPL-15 | ref\|NC_001146\|:353375 | A | C |
| YPL-15 | ref\|NC_001147\|:254470 | C | T |
| YPL-15 | ref\|NC_001147\|:343810 | C | T |
| YPL-16 | ref\|NC_001136\|:202886 | C | T |
| YPL-16 | ref\|NC_001136\|:619346 | C | T |
| YPL-16 | ref\|NC_001136\|:1121148 | T | G |
| YPL-16 | ref\|NC_001137\|:230418 | G | A |
| YPL-16 | ref\|NC_001139\|:218418 | C | A |
| YPL-16 | ref\|NC_001139\|:511347 | T | C |
| YPL-16 | ref\|NC_001140\|:192841 | G | A |
| YPL-16 | ref\|NC_001144\|:20982 | T | C |
| YPL-16 | ref\|NC_001145\|:479722 | T | G |
| YPL-16 | ref\|NC_001146\|:100669 | G | T |
| YPL-16 | ref\|NC_001147\|:135684 | C | A |
| YPL-16 | ref\|NC_001148\|:448887 | C | T |
| YPL-17 | ref\|NC_001133\|:172832 | G | A |
| YPL-17 | ref\|NC_001134\|:583520 | G | A |
| YPL-17 | ref\|NC_001136\|:323237 | T | G |
| YPL-17 | ref\|NC_001137\|:69277 | C | A |
| YPL-17 | ref\|NC_001139\|:431421 | C | CTAT |
| YPL-17 | ref\|NC_001139\|:1050029 | G | A |
| YPL-17 | ref\|NC_001140\|:424516 | T | G |
| YPL-17 | ref\|NC_001145\|:277560 | T | A |
| YPL-17 | ref\|NC_001147\|:841782 | T | G |
| YPL-18 | ref\|NC_001133\|:83992 | A | C |
| YPL-18 | ref\|NC_001139\|:466245 | T | C |
| YPL-18 | ref\|NC_001140\|:441858 | C | T |
| YPL-18 | ref\|NC_001143\|:15562 | G | C |
| YPL-18 | ref\|NC_001145\|:263584 | T | A |
| YPL-18 | ref\|NC_001145\|:873039 | C | A |
| YPL-19 | ref\|NC_001136\|:1491343 | G | C |
| YPL-19 | ref\|NC_001137\|:208031 | G | T |
| YPL-19 | ref\|NC_001138\|:237730 | C | T |
| YPL-19 | ref\|NC_001143\|:42335 | C | CTTA |
| YPL-19 | ref\|NC_001143\|:392249 | C | T |
| YPL-19 | ref\|NC_001144\|:226869 | T | C |
| YPL-19 | ref\|NC_001144\|:1014007 | C | T |
| YPL-19 | ref\|NC_001144\|:1042503 | A | T |
| YPL-19 | ref\|NC_001146\|:166168 | T | G |
| YPL-19 | ref\|NC_001147\|:413234 | G | T |
| YPL-2 | ref\|NC_001134\|:106253 | G | T |
| YPL-2 | ref\|NC_001136\|:286978 | C | T |
| YPL-2 | ref\|NC_001139\|:524238 | C | A |
| YPL-2 | ref\|NC_001139\|:677458 | A | G |
| YPL-2 | ref\|NC_001144\|:924055 | C | G |
| YPL-2 | ref\|NC_001147\|:309087 | T | TTA |
| YPL-20 | ref\|NC_001136\|:1107455 | C | T |
| YPL-20 | ref\|NC_001138\|:64632 | A | G |
| YPL-20 | ref\|NC_001139\|:469242 | A | ATTG |
| YPL-20 | ref\|NC_001140\|:556108 | G | A |
| YPL-20 | ref\|NC_001142\|:411876 | G | T |
| YPL-20 | ref\|NC_001147\|:556043 | G | T |
| YPL-20 | ref\|NC_001148\|:858004 | T | G |
| YPL-21 | ref\|NC_001139\|:290083 | A | C |
| YPL-21 | ref\|NC_001146\|:400624 | T | G |
| YPL-21 | ref\|NC_001147\|:434293 | T | A |
| YPL-22 | ref\|NC_001134\|:57711 | C | G |
| YPL-22 | ref\|NC_001134\|:308139 | T | C |
| YPL-22 | ref\|NC_001134\|:513077 | T | G |
| YPL-22 | ref\|NC_001134\|:724042 | C | G |
| YPL-22 | ref\|NC_001135\|:80863 | G | A |
| YPL-22 | ref\|NC_001136\|:76042 | A | C |
| YPL-22 | ref\|NC_001136\|:144889 | C | T |
| YPL-22 | ref\|NC_001136\|:151861 | T | A |
| YPL-22 | ref\|NC_001136\|:758617 | T | G |
| YPL-22 | ref\|NC_001136\|:1266191 | G | T |
| YPL-22 | ref\|NC_001139\|:761668 | A | G |
| YPL-22 | ref\|NC_001139\|:762435 | C | T |
| YPL-22 | ref\|NC_001145\|:660464 | C | A |
| YPL-22 | ref\|NC_001148\|:145531 | G | A |
| YPL-23 | ref\|NC_001134\|:16872 | T | C |
| YPL-23 | ref\|NC_001134\|:675236 | C | T |
| YPL-23 | ref\|NC_001136\|:1276500 | T | C |
| YPL-23 | ref\|NC_001136\|:1328320 | G | C |
| YPL-23 | ref\|NC_001136\|:1491806 | AT | A |
| YPL-23 | ref\|NC_001136\|:1491811 | G | A |
| YPL-23 | ref\|NC_001138\|:148600 | A | C |
| YPL-23 | ref\|NC_001145\|:633059 | G | C |
| YPL-23 | ref\|NC_001146\|:400624 | T | G |
| YPL-23 | ref\|NC_001148\|:80097 | G | A |
| YPL-24 | ref\|NC_001136\|:1404790 | C | A |
| YPL-24 | ref\|NC_001140\|:497254 | C | A |
| YPL-24 | ref\|NC_001143\|:229783 | G | A |
| YPL-3 | ref\|NC_001134\|:64321 | C | A |
| YPL-3 | ref\|NC_001134\|:433570 | C | G |
| YPL-3 | ref\|NC_001137\|:477342 | C | A |
| YPL-3 | ref\|NC_001138\|:163241 | G | A |
| YPL-3 | ref\|NC_001142\|:55615 | A | C |
| YPL-3 | ref\|NC_001147\|:297061 | C | A |
| YPL-3 | ref\|NC_001147\|:1060154 | T | TTCA |
| YPL-4 | ref\|NC_001136\|:336148 | C | CT |
| YPL-4 | ref\|NC_001136\|:441919 | G | A |
| YPL-4 | ref\|NC_001136\|:592539 | A | ATGCAAATGCAAC |
| YPL-4 | ref\|NC_001136\|:704251 | G | T |
| YPL-4 | ref\|NC_001136\|:1245263 | T | C |
| YPL-4 | ref\|NC_001137\|:361149 | C | A |
| YPL-4 | ref\|NC_001139\|:799442 | C | T |
| YPL-4 | ref\|NC_001144\|:204652 | C | T |
| YPL-4 | ref\|NC_001144\|:637244 | T | C |
| YPL-4 | ref\|NC_001144\|:889450 | A | C |
| YPL-4 | ref\|NC_001148\|:156322 | A | C |
| YPL-4 | ref\|NC_001148\|:156324 | A | T |
| YPL-4 | ref\|NC_001148\|:156325 | A | T |
| YPL-4 | ref\|NC_001148\|:156326 | A | T |
| YPL-4 | ref\|NC_001148\|:156328 | G | T |
| YPL-5 | ref\|NC_001148\|:188106 | C | A |
| YPL-6 | ref\|NC_001138\|:181571 | A | T |
| YPL-6 | ref\|NC_001142\|:115223 | G | T |
| YPL-6 | ref\|NC_001142\|:115224 | A | T |
| YPL-6 | ref\|NC_001142\|:126763 | G | A |
| YPL-6 | ref\|NC_001142\|:525350 | G | A |
| YPL-6 | ref\|NC_001142\|:532582 | C | T |
| YPL-6 | ref\|NC_001144\|:630675 | G | T |
| YPL-6 | ref\|NC_001145\|:578659 | A | T |
| YPL-6 | ref\|NC_001145\|:651721 | G | A |
| YPL-6 | ref\|NC_001145\|:827736 | C | T |
| YPL-7 | ref\|NC_001134\|:482749 | A | G |
| YPL-7 | ref\|NC_001134\|:497982 | A | T |
| YPL-7 | ref\|NC_001134\|:594871 | GAAATTTTTTTGAAAATAATTGTG  TTTGCTTATCTAGAGCTTTATT | G |
| YPL-7 | ref\|NC_001135\|:276735 | C | A |
| YPL-7 | ref\|NC_001139\|:387559 | T | G |
| YPL-7 | ref\|NC_001139\|:442139 | T | TGAAGTAGAGGACGAAGTAGAGGAC |
| YPL-7 | ref\|NC_001140\|:122590 | A | C |
| YPL-7 | ref\|NC_001142\|:679808 | G | C |
| YPL-7 | ref\|NC_001147\|:988946 | A | G |
| YPL-7 | ref\|NC_001148\|:581943 | C | T |
| YPL-8 | ref\|NC_001133\|:100665 | C | A |
| YPL-8 | ref\|NC_001133\|:227401 | T | C |
| YPL-8 | ref\|NC_001134\|:710348 | C | A |
| YPL-8 | ref\|NC_001136\|:266003 | C | A |
| YPL-8 | ref\|NC_001136\|:906197 | GA | G |
| YPL-8 | ref\|NC_001137\|:372006 | GA | G |
| YPL-8 | ref\|NC_001139\|:31966 | G | A |
| YPL-8 | ref\|NC_001139\|:522124 | G | C |
| YPL-8 | ref\|NC_001145\|:110268 | G | T |
| YPL-8 | ref\|NC_001146\|:391913 | T | A |
| YPL-8 | ref\|NC_001147\|:408639 | C | T |
| YPL-8 | ref\|NC_001147\|:718985 | T | A |
| YPL-8 | ref\|NC_001148\|:369002 | G | A |
| YPL-8 | ref\|NC_001148\|:662906 | T | A |
| YPL-9 | ref\|NC_001140\|:461189 | T | G |
| YPL-9 | ref\|NC_001143\|:437443 | G | C |
| YPL-9 | ref\|NC_001147\|:110791 | C | CTAA |
| YPL-9 | ref\|NC_001148\|:65048 | T | G |
| YPX-1 | ref\|NC_001134\|:454676 | G | C |
| YPX-1 | ref\|NC_001139\|:295680 | A | C |
| YPX-1 | ref\|NC_001139\|:736869 | C | A |
| YPX-1 | ref\|NC_001144\|:859379 | C | A |
| YPX-1 | ref\|NC_001145\|:17858 | T | C |
| YPX-1 | ref\|NC_001147\|:102528 | T | G |
| YPX-1 | ref\|NC_001147\|:727380 | T | G |
| YPX-1 | ref\|NC_001147\|:822924 | GTGTTGTTGTTGT | G |
| YPX-1 | ref\|NC_001147\|:969147 | C | T |
| YPX-10 | ref\|NC_001135\|:290355 | C | T |
| YPX-10 | ref\|NC_001136\|:806559 | A | G |
| YPX-10 | ref\|NC_001137\|:304345 | C | T |
| YPX-10 | ref\|NC_001139\|:534985 | T | G |
| YPX-10 | ref\|NC_001140\|:86934 | G | T |
| YPX-10 | ref\|NC_001140\|:286246 | A | T |
| YPX-10 | ref\|NC_001143\|:181779 | A | G |
| YPX-10 | ref\|NC_001144\|:514982 | G | A |
| YPX-10 | ref\|NC_001147\|:1037517 | C | CAAT |
| YPX-11 | ref\|NC_001136\|:288306 | C | T |
| YPX-11 | ref\|NC_001136\|:289552 | A | C |
| YPX-11 | ref\|NC_001138\|:7950 | G | T |
| YPX-11 | ref\|NC_001138\|:95581 | T | C |
| YPX-11 | ref\|NC_001139\|:1292 | G | T |
| YPX-11 | ref\|NC_001139\|:80135 | C | T |
| YPX-11 | ref\|NC_001141\|:246420 | G | T |
| YPX-12 | ref\|NC_001136\|:653939 | C | T |
| YPX-12 | ref\|NC_001139\|:443944 | C | A |
| YPX-12 | ref\|NC_001144\|:129080 | T | G |
| YPX-12 | ref\|NC_001144\|:941127 | G | A |
| YPX-12 | ref\|NC_001147\|:453088 | CA | C |
| YPX-12 | ref\|NC_001147\|:1053620 | A | T |
| YPX-12 | ref\|NC_001148\|:883291 | C | A |
| YPX-13 | ref\|NC_001136\|:447998 | T | C |
| YPX-13 | ref\|NC_001139\|:337954 | A | G |
| YPX-13 | ref\|NC_001140\|:221382 | GA | G |
| YPX-13 | ref\|NC_001144\|:1046308 | A | G |
| YPX-13 | ref\|NC_001145\|:171114 | G | C |
| YPX-13 | ref\|NC_001145\|:633790 | A | G |
| YPX-13 | ref\|NC_001145\|:698996 | A | C |
| YPX-13 | ref\|NC_001148\|:934453 | G | A |
| YPX-14 | ref\|NC_001136\|:166116 | T | C |
| YPX-14 | ref\|NC_001136\|:541173 | T | A |
| YPX-14 | ref\|NC_001146\|:10807 | C | A |
| YPX-14 | ref\|NC_001146\|:418383 | A | C |
| YPX-15 | ref\|NC_001135\|:224795 | A | G |
| YPX-15 | ref\|NC_001136\|:194077 | A | G |
| YPX-15 | ref\|NC_001139\|:134849 | T | C |
| YPX-15 | ref\|NC_001145\|:238330 | G | GATA |
| YPX-15 | ref\|NC_001145\|:888784 | A | AT |
| YPX-16 | ref\|NC_001136\|:1292458 | C | T |
| YPX-16 | ref\|NC_001137\|:32058 | G | A |
| YPX-16 | ref\|NC_001139\|:120043 | T | C |
| YPX-16 | ref\|NC_001144\|:980707 | G | T |
| YPX-16 | ref\|NC_001148\|:29974 | G | T |
| YPX-16 | ref\|NC_001148\|:435983 | A | G |
| YPX-17 | ref\|NC_001136\|:1262858 | G | C |
| YPX-17 | ref\|NC_001137\|:468096 | A | G |
| YPX-17 | ref\|NC_001138\|:83622 | A | C |
| YPX-17 | ref\|NC_001138\|:146191 | T | G |
| YPX-17 | ref\|NC_001142\|:283688 | T | TTA |
| YPX-17 | ref\|NC_001142\|:634490 | T | A |
| YPX-17 | ref\|NC_001144\|:737449 | C | T |
| YPX-17 | ref\|NC_001145\|:415088 | A | T |
| YPX-17 | ref\|NC_001147\|:27206 | C | G |
| YPX-17 | ref\|NC_001147\|:98002 | C | A |
| YPX-17 | ref\|NC_001147\|:626894 | A | C |
| YPX-18 | ref\|NC_001134\|:215205 | AT | A |
| YPX-18 | ref\|NC_001134\|:813066 | G | A |
| YPX-18 | ref\|NC_001134\|:813081 | G | A |
| YPX-18 | ref\|NC_001136\|:584737 | C | A |
| YPX-18 | ref\|NC_001136\|:823256 | G | GA |
| YPX-19 | ref\|NC_001135\|:72434 | C | T |
| YPX-19 | ref\|NC_001143\|:85785 | C | T |
| YPX-19 | ref\|NC_001143\|:114621 | A | C |
| YPX-19 | ref\|NC_001144\|:834527 | A | G |
| YPX-19 | ref\|NC_001144\|:837956 | G | GA |
| YPX-19 | ref\|NC_001145\|:25799 | A | C |
| YPX-19 | ref\|NC_001145\|:69881 | G | A |
| YPX-19 | ref\|NC_001147\|:496770 | G | A |
| YPX-2 | ref\|NC_001134\|:476572 | G | T |
| YPX-2 | ref\|NC_001135\|:316564 | TGTGTGG | T |
| YPX-2 | ref\|NC_001136\|:623840 | G | C |
| YPX-2 | ref\|NC_001138\|:222196 | ACTGGCGCTGGCG | A |
| YPX-20 | ref\|NC_001139\|:209409 | T | C |
| YPX-20 | ref\|NC_001147\|:544861 | A | G |
| YPX-21 | ref\|NC_001133\|:18355 | A | C |
| YPX-21 | ref\|NC_001141\|:210690 | C | T |
| YPX-21 | ref\|NC_001144\|:34586 | G | T |
| YPX-22 | ref\|NC_001134\|:556210 | G | GAT |
| YPX-22 | ref\|NC_001136\|:1367554 | C | G |
| YPX-22 | ref\|NC_001136\|:1426481 | G | A |
| YPX-22 | ref\|NC_001142\|:116397 | C | T |
| YPX-22 | ref\|NC_001144\|:43062 | T | G |
| YPX-22 | ref\|NC_001146\|:423497 | A | C |
| YPX-22 | ref\|NC_001146\|:734881 | C | T |
| YPX-23 | ref\|NC_001134\|:96571 | C | T |
| YPX-23 | ref\|NC_001139\|:1040639 | C | A |
| YPX-23 | ref\|NC_001143\|:233296 | T | G |
| YPX-23 | ref\|NC_001144\|:159144 | C | T |
| YPX-23 | ref\|NC_001144\|:837451 | G | T |
| YPX-23 | ref\|NC_001147\|:6330 | C | T |
| YPX-24 | ref\|NC_001136\|:639207 | C | G |
| YPX-24 | ref\|NC_001139\|:731558 | GC | G |
| YPX-24 | ref\|NC_001144\|:659022 | C | A |
| YPX-24 | ref\|NC_001144\|:1039918 | G | T |
| YPX-24 | ref\|NC_001146\|:471940 | A | G |
| YPX-3 | ref\|NC_001134\|:110592 | G | A |
| YPX-3 | ref\|NC_001134\|:608312 | G | A |
| YPX-3 | ref\|NC_001141\|:94754 | G | C |
| YPX-3 | ref\|NC_001143\|:456334 | C | A |
| YPX-3 | ref\|NC_001147\|:4157 | A | G |
| YPX-3 | ref\|NC_001147\|:87610 | T | A |
| YPX-3 | ref\|NC_001147\|:145760 | T | C |
| YPX-4 | ref\|NC_001136\|:1310950 | T | C |
| YPX-4 | ref\|NC_001138\|:95244 | ATTTTT | A |
| YPX-4 | ref\|NC_001138\|:95249 | T | TAAAAA |
| YPX-4 | ref\|NC_001139\|:957479 | G | A |
| YPX-4 | ref\|NC_001142\|:347962 | C | T |
| YPX-4 | ref\|NC_001143\|:491042 | A | AT |
| YPX-4 | ref\|NC_001147\|:293595 | G | T |
| YPX-5 | ref\|NC_001134\|:479153 | AG | A |
| YPX-5 | ref\|NC_001139\|:364421 | G | A |
| YPX-5 | ref\|NC_001144\|:577230 | T | C |
| YPX-5 | ref\|NC_001145\|:888537 | GTTC | G |
| YPX-5 | ref\|NC_001146\|:24868 | A | C |
| YPX-5 | ref\|NC_001147\|:43204 | T | G |
| YPX-5 | ref\|NC_001148\|:903842 | G | T |
| YPX-6 | ref\|NC_001136\|:217667 | C | A |
| YPX-6 | ref\|NC_001139\|:312313 | A | C |
| YPX-6 | ref\|NC_001148\|:123359 | G | A |
| YPX-7 | ref\|NC_001136\|:722221 | G | GA |
| YPX-7 | ref\|NC_001141\|:350478 | A | C |
| YPX-7 | ref\|NC_001143\|:568487 | C | A |
| YPX-8 | ref\|NC_001134\|:620983 | A | G |
| YPX-8 | ref\|NC_001136\|:556579 | G | T |
| YPX-8 | ref\|NC_001136\|:974316 | T | G |
| YPX-8 | ref\|NC_001136\|:1224227 | C | T |
| YPX-8 | ref\|NC_001136\|:1336742 | T | G |
| YPX-8 | ref\|NC_001139\|:696779 | C | T |
| YPX-8 | ref\|NC_001139\|:929653 | G | A |
| YPX-8 | ref\|NC_001140\|:35731 | G | T |
| YPX-8 | ref\|NC_001142\|:538548 | C | T |
| YPX-8 | ref\|NC_001143\|:347886 | G | A |
| YPX-8 | ref\|NC_001144\|:1006965 | G | A |
| YPX-8 | ref\|NC_001145\|:725360 | A | T |
| YPX-8 | ref\|NC_001147\|:446494 | A | G |
| YPX-9 | ref\|NC_001134\|:411007 | G | A |
| YPX-9 | ref\|NC_001137\|:174270 | C | A |
| YPX-9 | ref\|NC_001137\|:517141 | T | C |
| YPX-9 | ref\|NC_001142\|:44517 | G | A |
| YPX-9 | ref\|NC_001144\|:110811 | A | G |
| YPX-9 | ref\|NC_001145\|:553577 | G | A |
