## Supplementary material for "Mutations and structural variants arising during double-strand break repair": TableS4

| **Set** | **Forward** | **Reverse** | **Comments** |
| --- | --- | --- | --- |
| Set 1 | ACACTCTTTCCCTACACGACGCTCTTCCGATCT | GACTGGAGTTCAGACGTGTGCTCTTCCGATCT | Amplification of *Kl-ura3* |
| Set 1 | *ACACTCTTTCCCTACACGACGCTCTTCCGATCT*  ACAAGTTCTTGATATTTGAGGACAG | *GACTGGAGTTCAGACGTGTGCTCTTCCGATCT*  GCAATGAACCCAATAACGAAATC | NGS amplicon sequencing |
| Set 2, replicate 1 | TCAAGAAGGACAACATGTCCAC | *CTGTCT*CATTTGTCATCCGTCCCGTATAG | PacBio |
| Set 2, replicate 2 | TCAAGAAGGACAACATGTCCAC | *TAGTGG*CATTTGTCATCCGTCCCGTATAG | PacBio |
| Set 3 | *ACACACGCGAGACAG*ATATCAAAGAAAATCAAG  AAGGACAAC | *CGCATGACACGTGTGT*ATCCAACAACAACCTA  GAGTAATGG | PacBio |
| Digested Sample | *CTATACGTATATCTAT*TATCAAAGAAAATCAAGA  AGGACAAC | *GATATACGCGAGAGAG*ATCCAACAACAACCTA  GAGTAATGG | PacBio |
|  | Sequences used for barcoding are italicized. |  |  |
