## Supplementary material for "Mutations and structural variants arising during double-strand break repair": Table S3

| **Strain Name** | **Genotype** |
| --- | --- |
| tNS2921 | *ho∆ hmlΔ::ADE1 MAT****α*** *hmr****a****1::Kl-ura3-32Δ1,100BP sir3Δ::HphMX ade1 leu2 lys5 trp1::hisG ura3-52 ade3::GAL-HO* |
| tNS2922 | *ho∆ hmlΔ::ADE1 MAT****α*** *hmr****a****1::Kl-ura3-32Δ1,100BP sir3Δ::HphMX ade1 leu2 lys5 trp1::hisG ura3-100BP ade3::GAL-HO* |
| tNS2923 | *ho∆ hmlΔ::ADE1 MAT****α*** *hmr****a****1::Kl-ura3-32Δ1,100BP sir3Δ::HphMX ade1 leu2 lys5 trp1::hisG ura3-100BPmm ade3::GAL-HO* |
| TOY47 | *ho∆ hmlΔ::ADE1 MAT****α*** *hmr****a****1::Kl-ura3-32Δ1 ade1 leu2 lys5 trp1::hisG ade3::Gal-HO ura3Δ::Kl-ura3-804::NatMX* |
| WH50 | *ho∆ hmlΔ::ADE1 MAT****α*** *hmr****a****1::Kl-URA3 ade1 leu2 lys5 trp1::hisG ura3-52 ade3::Gal-HO* |
| YCM1 | *ho∆ hmlΔ::ADE1 MAT****α*** *hmr****a****1::Kl-ura3-32Δ2 sir3Δ::HphMX ade1 leu2 lys5 trp1::hisG ura3-52 ade3::GALHO* |
| yQW851 | *ho∆ hmlΔ::ADE1 MAT****α*** *hmr****a****1::Kl-ura3-32Δ1 ade1 leu2 lys5 trp1::hisG ade3::Gal::HO ura3::NatMX nej1::KanMX* |
| yQW852 | *ho∆ hmlΔ::ADE1 MAT****α*** *hmr****a****1::Kl-ura3-32Δ1 ade1 leu2 lys5 trp1::hisG ade3::Gal-HO ura3::NatMX nej1::KanMX Chr16-Kl-ura3-804* |
| yQW856 | *ho∆ hmlΔ::ADE1 MAT****α*** *hmr****a****1::Kl-ura3-32Δ1 ade1 leu2 lys5 trp1::hisG ade3::Gal-HO ura3::NatMX nej1::KanMX Chr16-Kl-ura3-132* |
| yQW857 | *ho∆ hmlΔ::ADE1 MAT****α*** *hmr****a****1::Kl-ura3-32Δ1 ade1 leu2 lys5 trp1::hisG ade3::Gal-HO ura3::NatMX nej1::KanMX Chr16-Kl-ura3-232* |
| yQW858 | *ho∆ hmlΔ::ADE1 MAT****α*** *hmr****a****1::Kl-ura3-32Δ1 ade1 leu2 lys5 trp1::hisG ade3::Gal-HO ura3::NatMX nej1::KanMX Chr16-Kl-ura3-332* |
| yQW859 | *ho∆ hmlΔ::ADE1 MAT****α*** *hmr****a****1::Kl-ura3-32Δ1 ade1 leu2 lys5 trp1::hisG ade3::Gal-HO ura3::NatMX nej1::KanMX Chr6-Kl-ura3-804* |
| yQW860 | *ho∆ hmlΔ::ADE1 MAT****α*** *hmr****a****1::Kl-ura3-32Δ1 ade1 leu2 lys5 trp1::hisG ade3::Gal-HO ura3::NatMX nej1::KanMX Chr7-Kl-ura3-804* |
| yQW861 | *ho∆ hmlΔ::ADE1 MAT****α*** *hmr****a****1::Kl-ura3-32Δ1 ade1 leu2 lys5 trp1::hisG ade3::Gal-HO ura3::NatMX nej1::KanMX Chr8-Kl-ura3-804* |
| yQW863 | *ho∆ hmlΔ::ADE1 MAT****α*** *hmr****a****1::Kl-ura3-32Δ1 ade1 leu2 lys5 trp1::hisG ade3::Gal-HO ura3::NatMX nej1::KanMX Chr6-Kl-ura3-132* |
| yQW864 | *ho∆ hmlΔ::ADE1 MAT****α*** *hmr****a****1::Kl-ura3-32Δ1 ade1 leu2 lys5 trp1::hisG ade3::Gal-HO ura3::NatMX nej1::KanMX Chr6-Kl-ura3-232* |
| yQW865 | *ho∆ hmlΔ::ADE1 MAT****α*** *hmr****a****1::Kl-ura3-32Δ1 ade1 leu2 lys5 trp1::hisG ade3::Gal-HO ura3::NatMX nej1::KanMX Chr6-Kl-ura3-332* |
| yQW866 | *ho∆ hmlΔ::ADE1 MAT****α*** *hmr****a****1::Kl-ura3-32Δ1 ade1 leu2 lys5 trp1::hisG ade3::Gal-HO ura3::NatMX nej1::KanMX Chr7-Kl-ura3-132* |
| yQW867 | *ho∆ hmlΔ::ADE1 MAT****α*** *hmr****a****1::Kl-ura3-32Δ1 ade1 leu2 lys5 trp1::hisG ade3::Gal-HO ura3::NatMX nej1::KanMX Chr7-Kl-ura3-232* |
| yQW868 | *ho∆ hmlΔ::ADE1 MAT****α*** *hmr****a****1::Kl-ura3-32Δ1 ade1 leu2 lys5 trp1::hisG ade3::Gal-HO ura3::NatMX nej1::KanMX Chr7-Kl-ura3-332* |
| yQW869 | *ho∆ hmlΔ::ADE1 MAT****α*** *hmr****a****1::Kl-ura3-32Δ1 ade1 leu2 lys5 trp1::hisG ade3::Gal-HO ura3::NatMX nej1::KanMX Chr8-Kl-ura3-132* |
| yQW870 | *ho∆ hmlΔ::ADE1 MAT****α*** *hmr****a****1::Kl-ura3-32Δ1 ade1 leu2 lys5 trp1::hisG ade3::Gal-HO ura3::NatMX nej1::KanMX Chr8-Kl-ura3-232* |
| yQW871 | *ho∆ hmlΔ::ADE1 MAT****α*** *hmr****a****1::Kl-ura3-32Δ1 ade1 leu2 lys5 trp1::hisG ade3::Gal-HO ura3::NatMX nej1::KanMX Chr8-Kl-ura3-332* |
| yQW892 and TOY043 | *ho∆ hmlΔ::ADE1 MAT****α*** *hmr****a****1::Kl-ura3-32Δ1 sir3Δ::HphMX ade1 leu2 lys5 trp1::hisG ura3-52 ade3::GALHO* |
| yQW904 | *ho∆ hmlΔ::ADE1 MAT****α*** *hmr****a****1::Kl-URA3 ade1 leu2 lys5 trp1::hisG ura3-52* ade3::Gal-HO* |

*Sc-ura3-52** contains a frameshift and point mutation that changes GGCGGCG to GGCGCC. *Kl-ura3-132* contains 132bp of *Kl-ura3* DNA (50bp+32bp+50bp) as a donor sequence (see text). *Kl-ura3-232, Kl-ura3-332,* and *Kl-ura3-804* respectively contain 232bp, 332bp and 804bp of *Kl-ura3* DNA. *Kl-ura3-32Δ2* contains a 32bp deletion located 5’ of *Kl-ura3-32Δ1* (see text). *Kl-ura3-100BP* contains 100bp of randomly generated DNA inserted at the end of the *Kl-ura3-32Δ1* open reading frame. *Kl-ura3-100BPmm* contains the same 100bp as *Kl-ura3-100BP* with mismatches (see text).
