## Supplemental Figures for "Mutations and structural variants arising during double-strand break repair"

### Supplementary Figures & Supplementary Figure Legends

Figure S1. SNVs and -1s are uniformly distributed across Kl-URA3, while MH sequences used for ICTS are central.

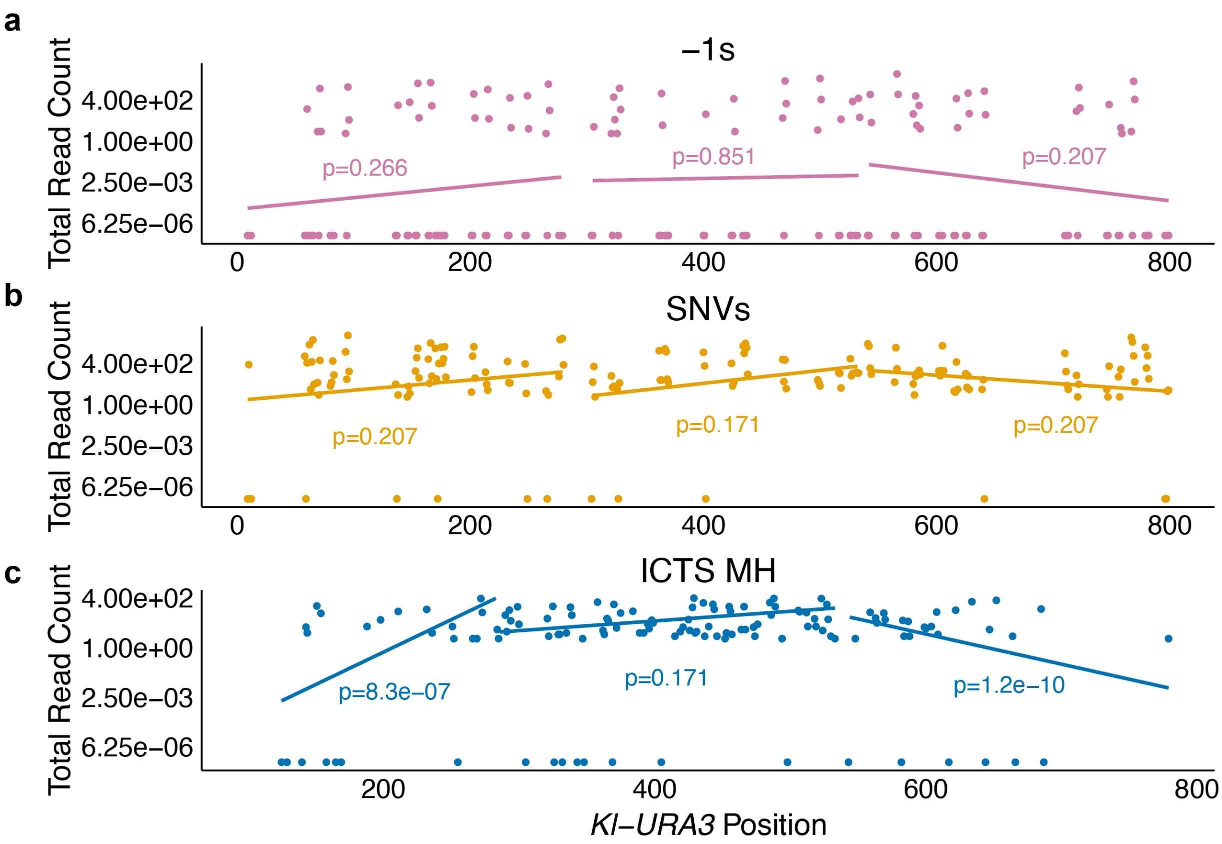


**Figure S1:**

SNVs and -1s are uniformly distributed across *Kl-URA3*, while ICTS MH sequences are exclusive to the center. **a** and **b**, The read counts summed across all three sets of SNVs (**a**) or -1 deletions (**b**) occurring within 3 bp homonucleotide runs. **c**, The read counts summed across all three sets of MHs used in the ICTS events were plotted vs the gene coordinate. Loci with no events are annotated with a read count of 0. The *Kl-URA3* sequence was split into thirds and a linear regression was fitted to the read counts of each event type within each third of the sequence. A one-sample two-tailed t-test calculates the probability that the slope of each line is different from 0. Annotated p-values were FDR-corrected. An annotated p-value less than 0.05 indicates that events of that type did not occur uniformly in the annotated region of *Kl-URA3*.

Figure S2. Frequency of MH usage between *Kl-URA3* and *Sc-ura3-52** to form ICTS events in Set 2.

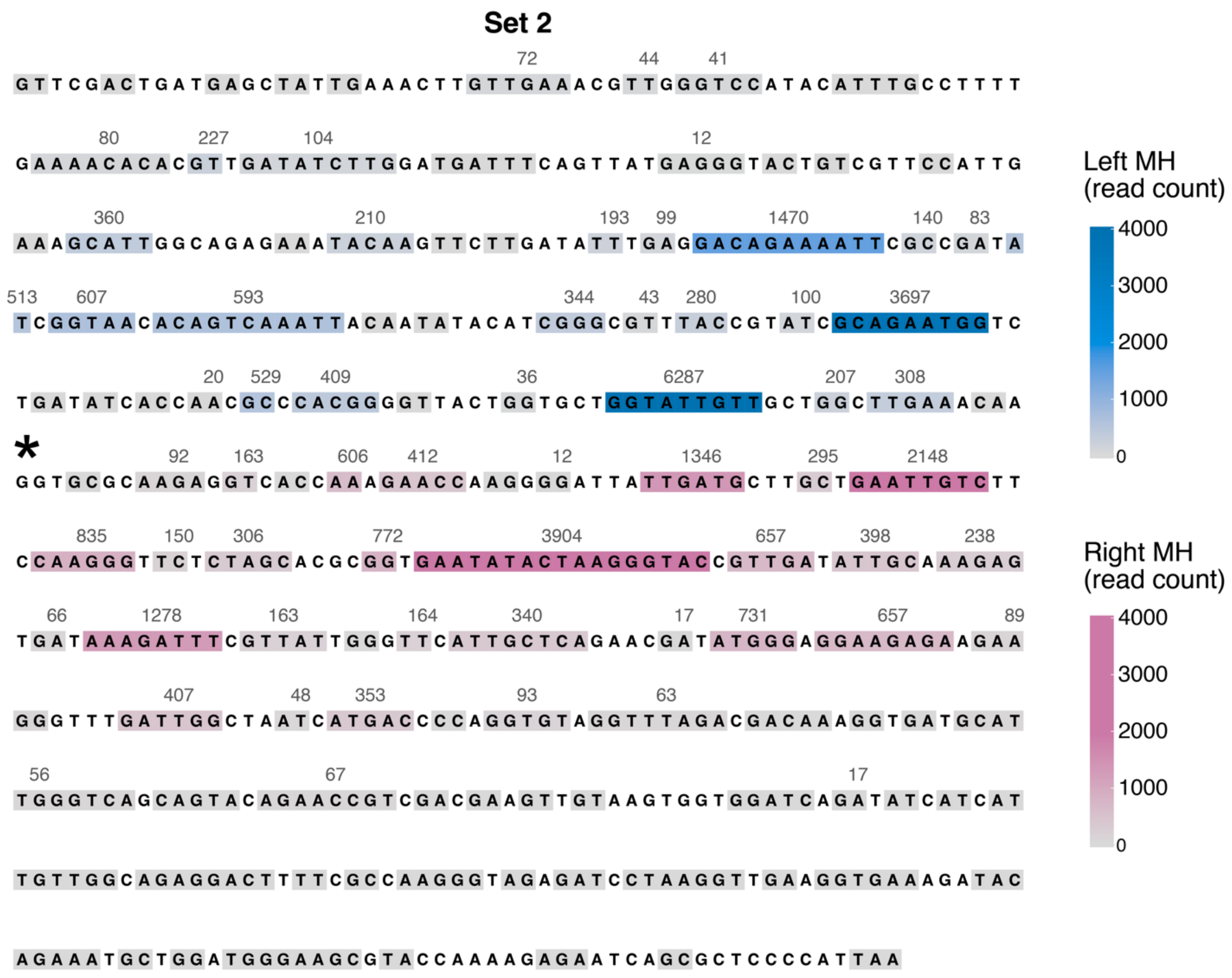


**Figure S2:**

Frequency of usage of MHs between *Kl-URA3* and *Sc-ura3-52** to form ICTS events in set 2. Grey boxes indicate unused MHs, blue boxes indicate MHs used to jump “in” to *Sc-ura3-52** and pink boxes indicate MHs used to jump “out” of *Sc-ura3-52**. Darker colors indicate more frequent usage. Black asterisk indicates the introduced -1 frameshift.

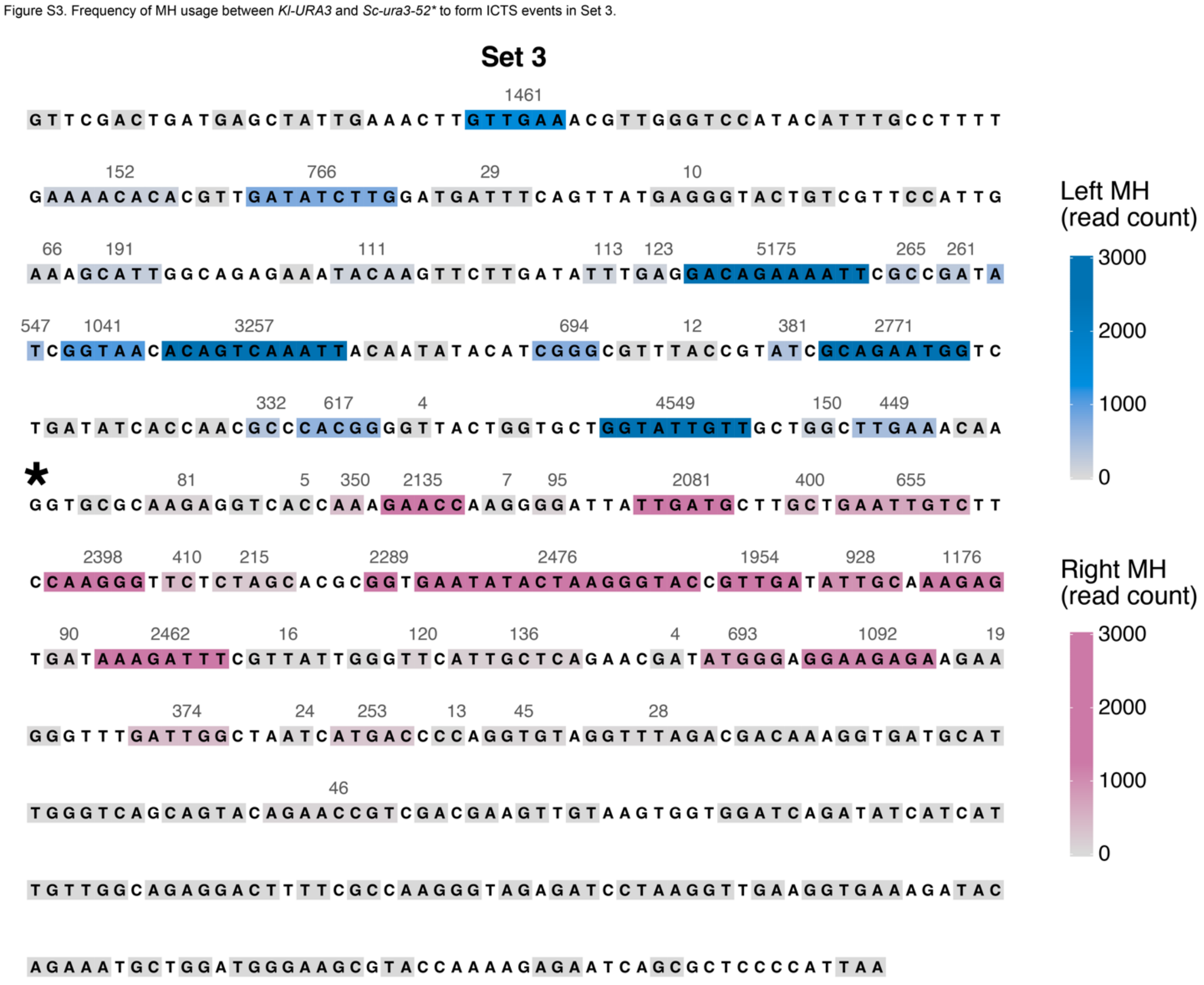


**Figure S3:**

Frequency of usage of MHs between *Kl-URA3* and *Sc-ura3-52** to form ICTS events in set 3. Grey boxes indicate unused MHs, blue boxes indicate MHs used to jump “in” to *Sc-ura3-52** and pink boxes indicate MHs used to jump “out” of *Sc-ura3-52**. Darker colors indicate more frequent usage. Black asterisk indicates the introduced -1 frameshift.

Figure S4. Microhomology usage in ICTS events enriched for intragenic deletions.

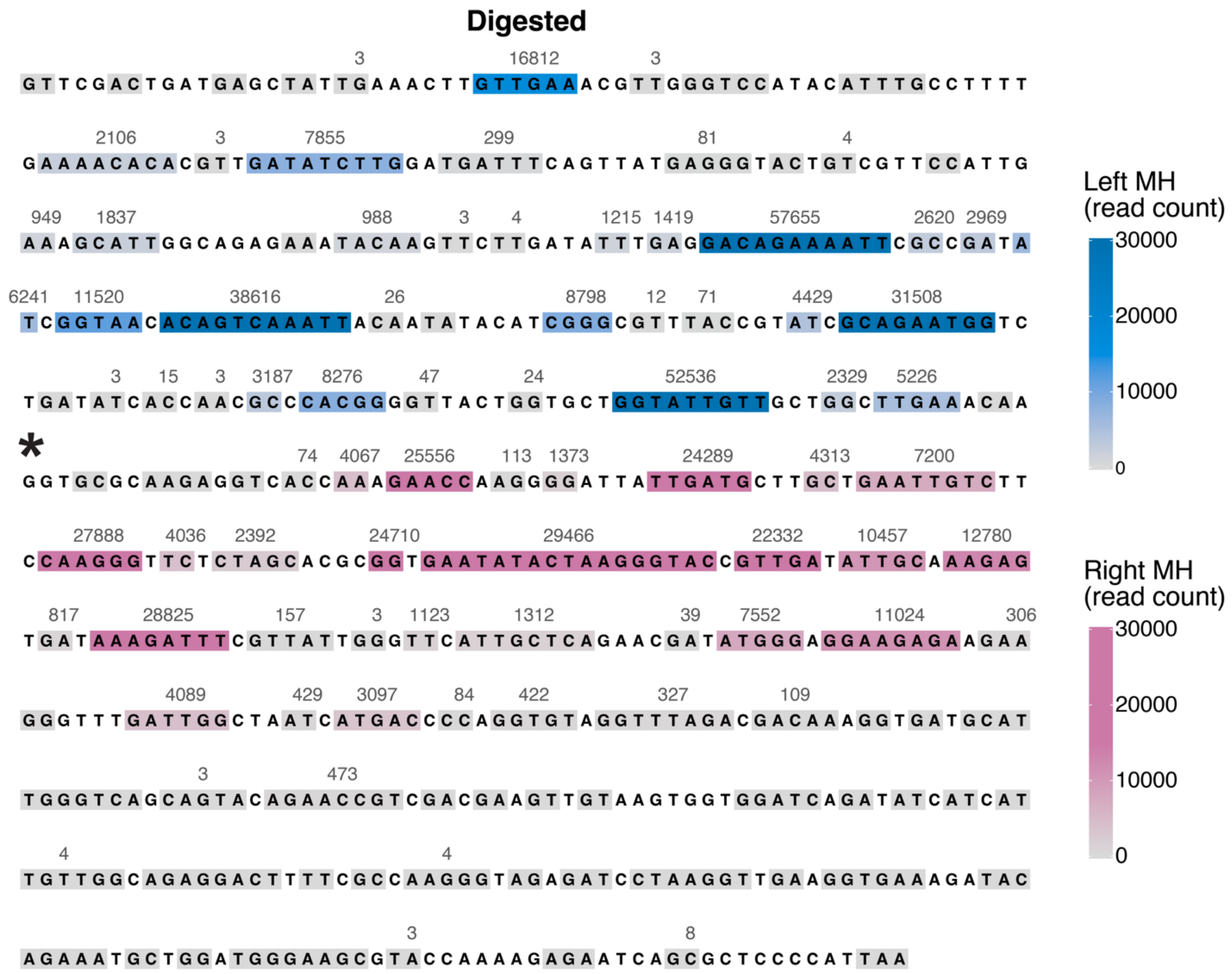


**Figure S4:**

Frequency of usage of MHs between *Kl-URA3* and *Sc-ura3-52** to form ICTS events in the *Bst*EII and *Fsp*I digested sample. Grey boxes indicate unused MHs, blue boxes indicate MHs used to jump “in” to *Sc-ura3-52** and pink boxes indicate MHs used to jump “out” of *Sc-ura3-52**. Darker colors indicate more frequent usage. Black asterisk indicates the introduced -1 frameshift.

Figure S5. Homeology within adjacent 25 bp does not play a role in MH choice for ICTS

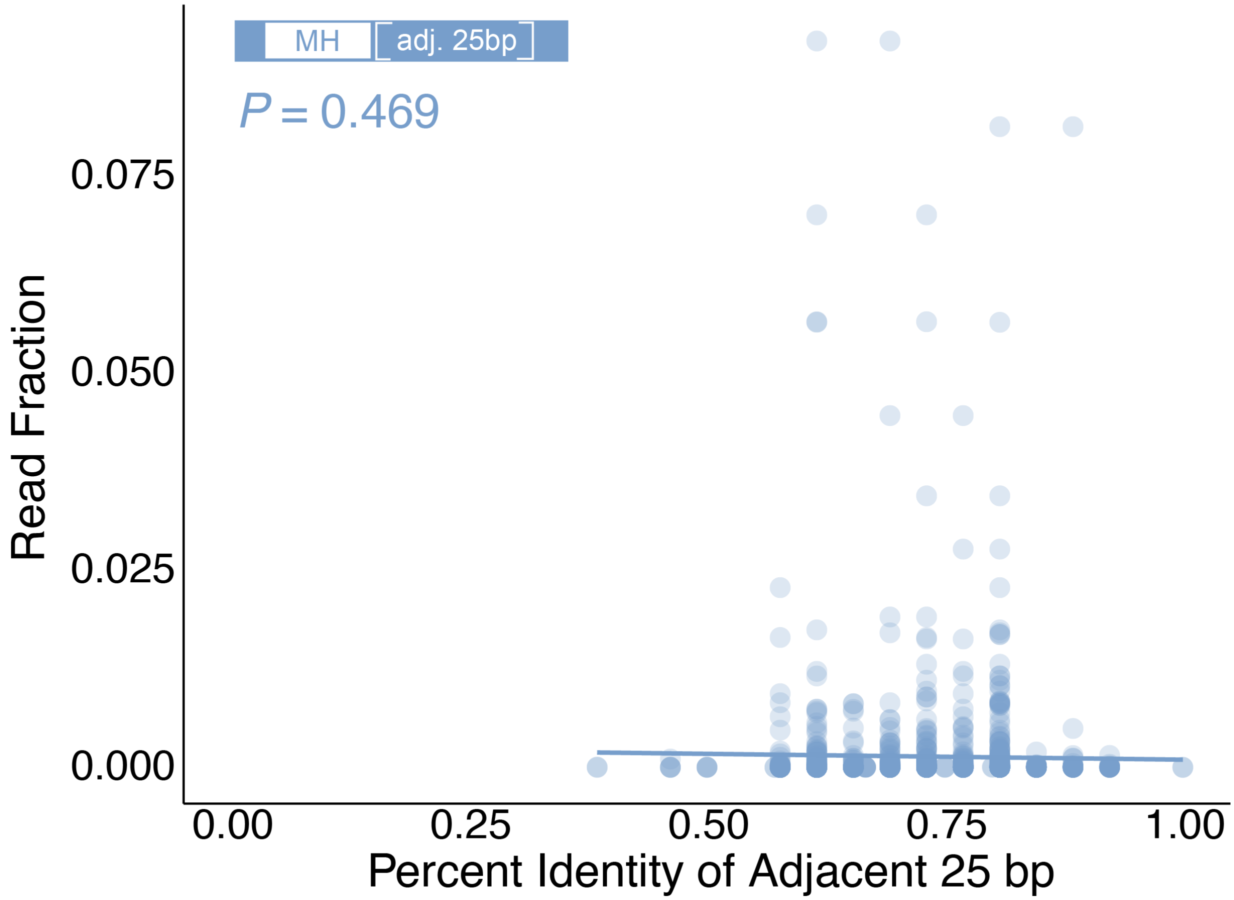


**Figure S5:**

Comparison of frequency of specific MH usage and homeology in the adjacent 25 bp across ICTS events enriched via *Bst*EII and *Fsp*I digestion. A linear regression was fit to the read fractions vs. percent homeology values. A one-sample two-tailed t-test calculates the probability that the slope of the line is different from 0.
